# A Complete Genome for the Common Marmoset

**DOI:** 10.64898/2026.03.25.713844

**Authors:** Prajna Hebbar, Tamara Potapova, Hailey Loucks, Karina Ray, Murillo F. Rodrigues, Fedor Ryabov, Joanna Malukiewicz, DongAhn Yoo, Leonardo de Lima, Annat Haber, Sonal Kumar, Swati Banerjee, Matthew Borchers, Gage H. Garcia, Joshua Gardner, Stephanie Hachem, Harrison Heath, Seung Kwon Ha, Mira Mastoras, Brandy McNulty, Julian Menendez, Katherine M. Munson, Karol Pal, JungEun Park, Simon Ploesch, Christian Roos, William E. Seligmann, Valery Shepelev, Catrina Spruce, Ivo Violich, Lutz Walter, Kateryna D. Makova, Amantha Thathiah, Stacey J. Sukoff Rizzo, Afonso C. Silva, Gregory W. Carter, Karen H. Miga, Evan E Eichler, Donald F. Conrad, Jennifer L. Gerton, Ivan Alexandrov, Benedict Paten

**Affiliations:** UC Santa Cruz Genomics Institute, University of California, Santa Cruz, Santa Cruz, CA 95060, USA; Stowers Institute for Medical Research, Kansas City, MO 64110, USA; Division of Genetics, Oregon National Primate Research Center, Beaverton, OR 97006, USA; Centre for Biomedical Research and Technology, HSE University, Moscow, Russia; The Center for Bio- and Medical Technologies, Moscow, Russia; Primate Genetics Laboratory, German Primate Center, Leibniz Institute for Primate Research, Göttingen, Germany; Instituto de Medicina Tropical de São Paulo, Universidade de São Paulo, São Paulo, SP, Brasil; Department of Genome Sciences, University of Washington School of Medicine, Seattle, WA 98195, USA; The Jackson Laboratory, Bar Harbor, Maine 04609, USA; Graduate School of Biomedical Sciences, Tufts University, Boston 02111, MA, USA; Department of Neurobiology, University of Pittsburgh School of Medicine, Pittsburgh, PA 15213, USA; Department of Biology, Penn State University, University Park, PA 16802, USA; Institute of Molecular Genetics, Moscow, Russia (currently retired); Howard Hughes Medical Institute, University of Washington, Seattle, WA 98195, USA; Department of Human Molecular Genetics and Biochemistry, Faculty of Medical and Health Sciences, Tel Aviv University, Israel

**Author notes:** Corresponding Author: Benedict Paten.

## Abstract

The common marmoset is a New World monkey (NWM) commonly used as a model organism to investigate questions in primate evolution and human disease, including Alzheimer’s and other neurodegenerative diseases, as well as neuropsychiatric disorders. Here we present the first telomere-to-telomere (T2T) reference genome for the common marmoset, adding over 88 Mb of sequence and resolving challenging genomic regions. An additional near-T2T assembly from a second unrelated individual yields a total of four high-quality haplotypes for analysis. The improved contiguity and accuracy of these assemblies enable unprecedented insights into complex and rapidly evolving genomic regions such as centromeres, sex chromosomes, ribosomal DNA (rDNA) structure, and the major histocompatibility complex (MHC). We fully resolved all marmoset centromeres, uncovering dimeric alpha satellites with chromosomal specificity and stratified inactive layers documenting ancestral centromere turnover. We assembled six acrocentric autosomes with gene-poor, satellite-rich short arms and provide evidence that most of them can harbor rDNA and all of them share large pseudo-homologous regions (PHRs). The Y chromosome, but not the X chromosome, carries active rDNA and PHRs, and the rDNA copy number is sexually dimorphic. Chromosomes that share PHRs also share closely related centromeric satellite DNA, supporting a model of ongoing recombinational exchange between heterologous chromosomes facilitated by rDNA. We discovered multiple novel, marmoset-specific MHC genes that are predicted to protect against pathogens encountered in its environment. Leveraging this complete reference, we further identified over 500 transcribed genes with transcript models or expansions specific to the marmoset lineage. Together with additional long-read marmoset assemblies, these genomes were used to construct a marmoset pangenome, providing a robust reference framework for short-read mapping across diverse individuals. This resource will improve the utility of the common marmoset as a biomedical model organism and fill key gaps in our understanding of primate evolution.

## Introduction

The common marmoset (*Callithrix jacchus*) has emerged as an indispensable model organism for understanding primate biology, neuroscience, and human disease. Native to the forests and semi-deserts of northeastern Brazil^1^, these small New World monkeys possess several attributes that make them particularly valuable for biomedical research-their compact size, relatively short generation time, high reproductive output, and their susceptibility to age-related, neurodegenerative human diseases, including Alzheimer’s disease^2^, and neuropsychiatric diseases, including anxiety and depression^3,4^. Unlike traditional rodent models, marmosets share critical physiological and neurological features with humans while offering practical advantages over larger primates. As such, they occupy a unique niche in translational research, bridging the gap between mice and macaques while providing insights into primate-specific evolutionary innovations^5–8^.

Despite the common marmoset’s growing prominence in biomedical research, the genomic resources available for this species have remained incomplete. The current reference genome assemblies contain numerous gaps and errors, particularly in structurally complex regions such as centromeres, telomeres, segmental duplications, satellite-rich short arms of acrocentric chromosomes, and the sex chromosomes^9–13^. These limitations have constrained investigations into fundamental questions about primate genome evolution, structural variation, and the genetic basis of lineage-specific adaptations. Incomplete assemblies have also hampered gene annotation, including in loci relevant to disease modeling. These issues have obscured our understanding of gene family evolution, immune system diversity, and chromosomal organization.

Recent advances in long-read sequencing technologies and telomere-to-telomere (T2T) assembly methods have revolutionized our ability to resolve these challenging genomic regions^14–16^. The successful completion of the human T2T genome and subsequent T2T assemblies for great apes have demonstrated that these problematic genomic regions can now be fully resolved, showing that these repetitive and duplicated sequences contain novel genes, satellites, and structural variants^17–21^. Similarly, long-read transcriptomic technologies such as IsoSeq complement these advances in assembly, enabling the recovery of missing gene models. These technological breakthroughs provide an opportunity to generate complete, gapless reference genomes and transcriptomes for non-human primates, enabling more accurate comparative genomics and functional studies.

Centromeres are essential for correct chromosome segregation and, in primates, are typically associated with tandemly repeated alpha-satellite DNA. Yet, their repetitive nature has made them resistant to assembly and largely excluded them from reference genomes. In primates, centromeric alpha satellites range from simple monomeric arrays to highly organized higher-order repeat (HOR) structures, with distinct architectures in great apes and Old World monkeys now well characterized thanks to T2T assemblies^18–20,22^. In contrast, centromere organization in New World monkeys remains largely uncharacterized at the sequence level. However, there is cytogenetic evidence that their alpha satellites are predominantly dimeric and lack the HOR structures typical of hominids^23,24^. Similarly, acrocentric chromosomes, those with centromeres positioned near one end, also pose unique assembly challenges because their short arms are usually composed almost entirely of satellite DNA and segmental duplications. In many species, these arms also harbor rDNA arrays that form nucleolar organizing regions (NORs). Marmosets carry several acrocentric chromosomes, and the sequence content and structural organization of most of their short arms have not been resolved.

Here we present a telomere-to-telomere reference genome for the common marmoset, adding over 88 megabases of previously unresolved sequence and achieving completeness and high accuracy. We generated an additional near-T2T assembly from a second unrelated individual, providing four high-quality haplotype assemblies that capture extensive structural and genic variation within the species. These genomes enabled the characterization of New World monkey centromere architecture, full resolution of acrocentric chromosome short arms, which revealed highly variable rDNA array configurations, a transcriptionally active Y-chromosome rDNA array, and underlying sex-specific rDNA copy number differences, and haplotype-resolved assemblies of the MHC region, uncovering lineage-specific gene family expansions. The Y chromosome contains novel ampliconic gene families not found in apes, and genome-wide analysis showed elevated segmental duplication enriched for immune-related and reproductively relevant genes^25^. Along with long-read transcriptome data, a multispecies primate alignment, and a marmoset pangenome graph, these assemblies provide a comprehensive foundation for using the marmoset as a model organism to research human health and primate evolution.

## Results

To generate these assemblies, we used sequencing data generated from fibroblasts from a male (calJac240) and a female (calJac220) marmoset, both living in a United States research colony. Genome-wide SNP-based principal component analysis (PCA) confirms that these two individuals are representative of the broader population of research marmosets registered with the Marmoset Coordinating Center at OHSU (Fig. **1A**) and a different population of research marmosets in the University of Pittsburgh colony (Fig. **S1**). We integrated multiple sequencing technologies and employed Verkko^26^ for genome assembly. We generated PacBio high-fidelity (HiFi) sequencing at >60x coverage alongside Oxford Nanopore Technologies (ONT) ultra-long reads (UL >100 kb) at 30x coverage (Table **S1**), using high-molecular-weight DNA extracted from each sample. The ultra-long reads were used to scaffold across complex repetitive regions, including centromeric satellites and segmental duplications (SDs), thereby resolving repeats. Hi-C sequencing data from both samples enabled chromosome-scale haplotype phasing^26,27^ (Table **1**). The more complete and accurate haplotype for each chromosome pair, as measured by quality value (QV), was assigned to the primary assembly for each donor, with preference given to haplotypes harboring more complete rDNA arrays. The primary assembly of calJac240, which is the T2T genome, includes both sex chromosomes^19^.

**Figure 1.**
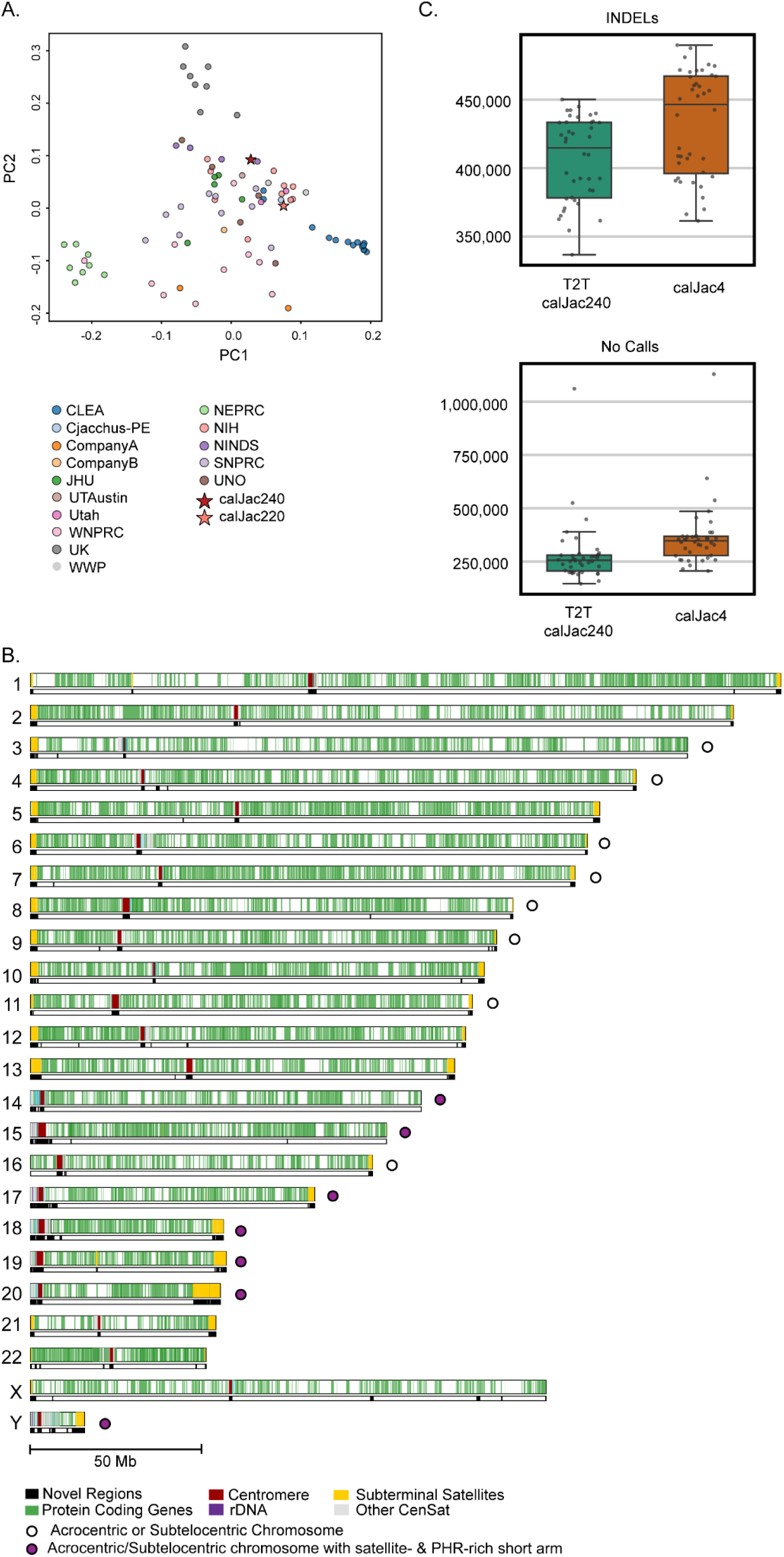
calJac240 and the T2T assembly. (A) PCA of marmoset samples colored by cohort/source institution. PC1 and PC2 capture the primary axes of genetic variation, with calJac240 (dark red star) and calJac220 (pink star) marked. (B) Ideogram of the calJac240 T2T assembly showing the genome-wide distribution of new regions compared to calJac4 (in black), protein-coding genes (green), alpha satellites (red), rDNA (purple), subterminal satellites (yellow), and other centromeric satellites (gray) across all 22 autosomes and the X and Y sex chromosomes. Open circles denote acrocentric or subtelocentric chromosomes; filled purple circles denote acrocentric/subtelocentric chromosomes with satellite- and PHR-rich short arms. (C) Box plots comparing the number of INDELs (top) and no-call sites (bottom) per sample when short-read data from a marmoset cohort were aligned to the T2T calJac240 assembly (green) versus the calJac4 reference (orange). Each point represents an individual sample.

**Table 1.** Assembly Statistics.

| Sample information |  |  | Assembly stats (Hap1/Hap2) |  |  |  |  |  |
| --- | --- | --- | --- | --- | --- | --- | --- | --- |
|  | Tissue | Sex | Total bases (Gbp) | contig N50 (Mbp) | Number of T2T contigs | QV (Illumina) | QV (hybrid) | QV (HiFi) |
| 240 | Fibroblasts | M | 2.76/2.93 | 141.93/141.41 | 15/16 | 58.53/58.08 | 67.32/67.11 | 66.08/66.02 |
| 220 | Fibroblasts | F | 2.94/2.93 | 142.5/142.1 | 17/16 | 57.80/58.34 | 65.29/66.89 | 64.29/65.69 |

Across the four haplotypes, we have one T2T and three near-T2T assemblies. 70% of chromosomes were assembled T2T, gapless (except for rDNA arrays) with telomeres on both ends across all the assemblies (Table **S2**). Flagger^28^ analysis points to an average of 0.9-1.5 Mb of non-rDNA-related assembly issues per haplotype (Table **S3**). QV measured with Illumina-based k-mers and hybrid k-mers (using both Illumina and HiFi reads) using Merqury^29^ was used to assess assembly accuracy further. QV scores of 57.80-58.53 were obtained from Illumina-based k-mer comparisons, though Illumina coverage biases constrain these estimates. Higher QV values of 65.29-67.32 were achieved through the hybrid validation method, and DeepVariant-based polishing contributed an approximately 1.5-point QV increase per haplotype^30,31^ (Table **S4**). Taken together, these results indicate that more than 99% of each genome was assembled completely and accurately, comparable to the T2T-CHM13 human reference and the T2T ape genomes, all of which exhibit potential issues in 0.3-1.2% of the genome^18,19^.

The assemblies resolved previously inaccessible sequence predominantly within acrocentric short arms and centromeric regions (30.24 Mb), segmental duplications (22.41 Mb), and other satellite arrays (36.13 Mb). More than 400 genes (8.41 Mb) were encompassed within these new sequences. Gene distribution across chromosomes follows expected density patterns, with repeat-rich regions now fully resolved (Fig. **1B**).

In both sequence accuracy and contiguity, these marmoset diploid assemblies represent a substantial improvement over all prior marmoset genome assemblies. Upon aligning short-read whole-genome sequencing data from a cohort of marmosets to both the previous reference assembly (*GCF_009663435.1*) and the primary T2T calJac240 assembly, we find considerable gains in reference completeness (Table **S5**). The number of INDELs called per sample is significantly lower when reads are aligned to the T2T assembly compared to calJac4 (Fig. **1C**, **top**). This reduction indicates that calJac240 harbors fewer gaps and incomplete regions that artificially inflate INDEL counts. Furthermore, the number of no-call sites per sample is significantly reduced with the T2T assembly, with approximately 84,000 fewer sites per animal where the genotype could not be determined (Fig. **1C**, **bottom**). Together, these results demonstrate that the T2T calJac240 assembly provides a more complete and continuous reference that reduces variant calling artifacts, improves genotyping success, and enables more reliable population-level analyses across the marmoset cohort.

### Resources

#### Alignments

To provide a framework for comparative genomics, we constructed two complementary alignment resources. First, we generated a multispecies alignment using Progressive Cactus^32^, incorporating the T2T-CHM13 human reference^18^, T2T great ape assemblies^19^, two T2T macaque assemblies^20,21^, our primary marmoset haplotype (calJac240), and dog and mouse as outgroups. This phylogenetically informed alignment enables direct comparison of the marmoset genome with other primate genomes while accounting for evolutionary relationships. To further facilitate translational research, we provide a chain file between the human T2T-CHM13 and calJac240 assemblies, enabling researchers using marmoset as a model organism to directly translate findings between human and marmoset genomes (Fig **2A**, Fig. **S2**). Second, we built a marmoset-specific pangenome using the four haplotypes generated in this study, along with one additional long-read haplotype-resolved assembly (*GCA_030222145.1* and *GCA_030222185.1*^33^, hereafter referred to as CJ1700) with minigraph-Cactus^34^. The resulting pangenome graph, spanning 2.99 Gb, captures structural variation within the species. Of the 58,381 SVs identified, 1,689 overlap protein-coding genes, including 71 cases where deletions fully encompass exons, representing potential pseudogenization events (Table **S6**). This pangenome is a step toward moving beyond single linear reference genomes for marmoset genetic research, providing a more comprehensive representation of genomic diversity within the species^35,36^.

**Figure 2.**
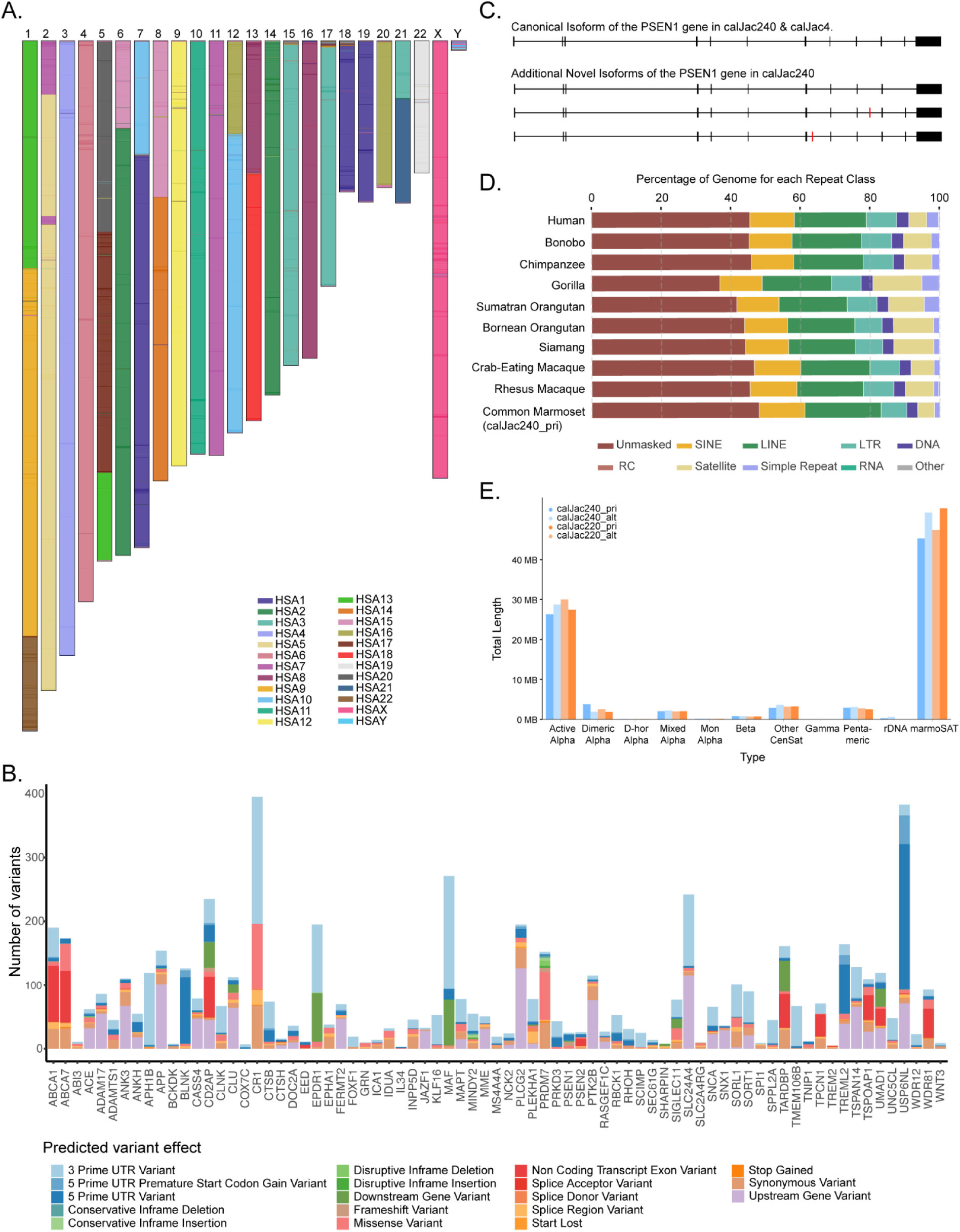
Chromosomal architecture and annotation of calJac240 assembly. (A) Karyotype of the calJac240 marmoset T2T assembly painted with human chromosome synteny, with marmoset chromosomes (1-22, X, Y) colored by their corresponding human chromosome (HSA) homology. (B) Predicted effects at the gene level for 75 AD-associated gene loci. Intronic variants have been excluded for ease of visualization. (C) Gene structure of the *PSEN1* locus, illustrating the canonical isoform present in both calJac240 and calJac4 (top) and three additional novel isoforms identified in calJac240 (bottom three tracks). Black boxes indicate exons, and red marks indicate novel exons. (D) Repeat element composition as a percentage of the genome across ten primate species with T2T genomes, including Human, Bonobo, Chimpanzee, Gorilla, Sumatran Orangutan, Bornean Orangutan, Siamang, Crab-eating Macaque, Rhesus Macaque, and Common Marmoset (calJac240). Repeat classes shown include SINE, LINE, LTR, DNA transposons, RC (rolling circle), satellite, simple repeat, RNA, and other elements, with the unmasked regions also indicated. (E) Total length of different satellite repeat types across the four marmoset haplotypes.

#### Gene annotation

To identify both protein-coding and noncoding RNA (ncRNA) genes for each marmoset haplotype, we applied the Comparative Annotation Toolkit 2 (CAT2) gene-annotation pipeline (improving on CAT^37^). Additional annotations are also available through NCBI RefSeq^38^. The number of protein-coding genes identified was comparable to that of other primates (n = 21,121). Based on GENCODE v46^39^, the CAT2 gene set indicates that 97.8% of corresponding human orthologs are now represented, with more than 90% assembled at full length, a 0.6% increase from the previous gene build. Long-read transcriptomic data (IsoSeq) from 10 different marmoset tissues (Table **S7**) were also incorporated to predict novel protein-coding genes within this gene set^33^. The majority of protein-coding genes show tissue expression (80% expressed in at least one tissue), with over 6,000 genes supported by IsoSeq data across all examined tissues, while approximately 4,000 genes were not supported by transcripts in any of the 10 tissues (Fig. **S3**). Around 9.3% of annotated transcripts have novel splice forms supported by IsoSeq data from at least two different tissues. These include novel transcript isoforms of the *PSEN1* gene (Fig **2C**), which is studied in marmoset models of Alzheimer’s disease^40^. We also identified several protein-coding genes in the marmoset T2T genome that contained new transcript models relative to the human genome annotation, either a gene novel to marmosets, a transcript significantly modified in marmosets, or a duplication specific to marmosets. This includes 566 genes with IsoSeq support from at least 2 tissues.

Among these are biomedically relevant genes such as novel MHC genes, genes related to spermatogenesis, and genes involved in tissue organization. As with the T2T Primates project, we also provide a curated consensus protein-coding gene annotation set by integrating results from both the CAT2 and NCBI pipelines^19^. All alignment and annotation resources are made available through a track hub for the UCSC Genome Browser^41^.

#### Variant burden in Alzheimer’s disease-associated loci

The common marmoset is becoming an established non-human primate model of human diseases, particularly disorders affecting the central nervous system^40,42^. Standing genetic variation in the marmoset may contribute to variability in disease-associated traits, allowing the marmoset to serve as a genetic model of disease risk and progression. To explore this possibility in Alzheimer’s disease, we shortlisted 81 genetic loci associated with early and/or late-onset Alzheimer’s disease and related pathology^43^, including *SNCA* in Parkinson’s disease^44^, and *TARDBP* in amyotrophic lateral sclerosis and frontotemporal dementia^45^. Of the 81 human loci, 76 loci with conserved candidate genes were identified in the marmoset genome. We used SnpEff^46^ to annotate variant effects and impacts in a population of 230 marmosets from the University of Pittsburgh colony.

Standing variation was observed in 75 of the tested loci, with variation absent only in *HS3ST5*. 93.3% of the observed variants were intronic, predicted to be modifiers, and unlikely to affect protein-coding sequences. Among the remaining variants, 3’UTR (1.99%) and upstream gene (1.35%) variants are the most common. Start-loss and splice-acceptor variants were present but rare (both <0.005%). Excluding intronic variants, the greatest number of variants were seen in *CR1* and *USP6NL,* and the fewest in *IL34* and *TREM2* (Fig. **2B**). Variants with modifier impacts are less likely to affect coding sequences; thus, we estimated impacts across genes ranging from low to high (Fig. **S4**, Table **S8**). *CR1*, *PRDM7*, and *ABCA7* had the highest cumulative predicted impact associated with identified variants, whereas *IL34* and *TMEM106B* had the fewest, with no impactful variants observed in *COXC7*. Multiple genes (e.g., *USP6NL*) with high variant frequencies had low cumulative impact because most variants were non-coding and likely play regulatory roles. The complement receptor *CR1*, on the other hand, exhibited high variant frequency with numerous potential coding impacts. These findings illustrate that standing variation in laboratory marmoset colonies may influence Alzheimer’s-related traits, potentially at the same loci as observed in humans.

#### Repeat annotation

We surveyed repetitive elements across all the assembled haplotypes (Table **S9**). Repetitive sequences, including transposable elements, satellite DNA, and variable number tandem repeats (VNTRs), constitute around 51.24% of the marmoset genome. The marmoset repeat landscape exhibits a composition broadly similar to other primates, with the major repeat classes, LINEs, SINEs, LTRs, DNA transposons, satellites, and simple repeats, present in proportions comparable to other species and consistent across all four haplotype assemblies (Fig. **2D**). Our T2T assembly provides an additional 66.37 Mb of repetitive content. This represents a notable improvement over previously reported satellite content, reflecting the enhanced resolution in these challenging genomic regions (Fig **2E**).

### Lineage-specific SDs and gene families

Segmental duplications (SDs) are large blocks of highly similar sequences (>90% identity, >1 kb in length) that arise through genomic rearrangements and represent hotspots for evolutionary innovation and structural variation. Analysis of SD content across the calJac240, calJac220 (this study), and CJ1700 assemblies showed that while marmosets possess fewer SDs than great apes, they harbor more duplicated sequences than other non-ape primates^19,47^. Specifically, the marmoset genomes contain 147.3-165.0 Mb of duplicated sequence per haplotype, representing 17.0-34.8 Mb more than the average of other non-ape primate species, including those with T2T genomes (crab-eating and rhesus macaques) (Fig. 3**A**). This duplicated content comprises 102.2 Mb/haplotype of intrachromosomal duplications and 80.9 Mb/haplotype of interchromosomal duplications.

**Figure 3.**
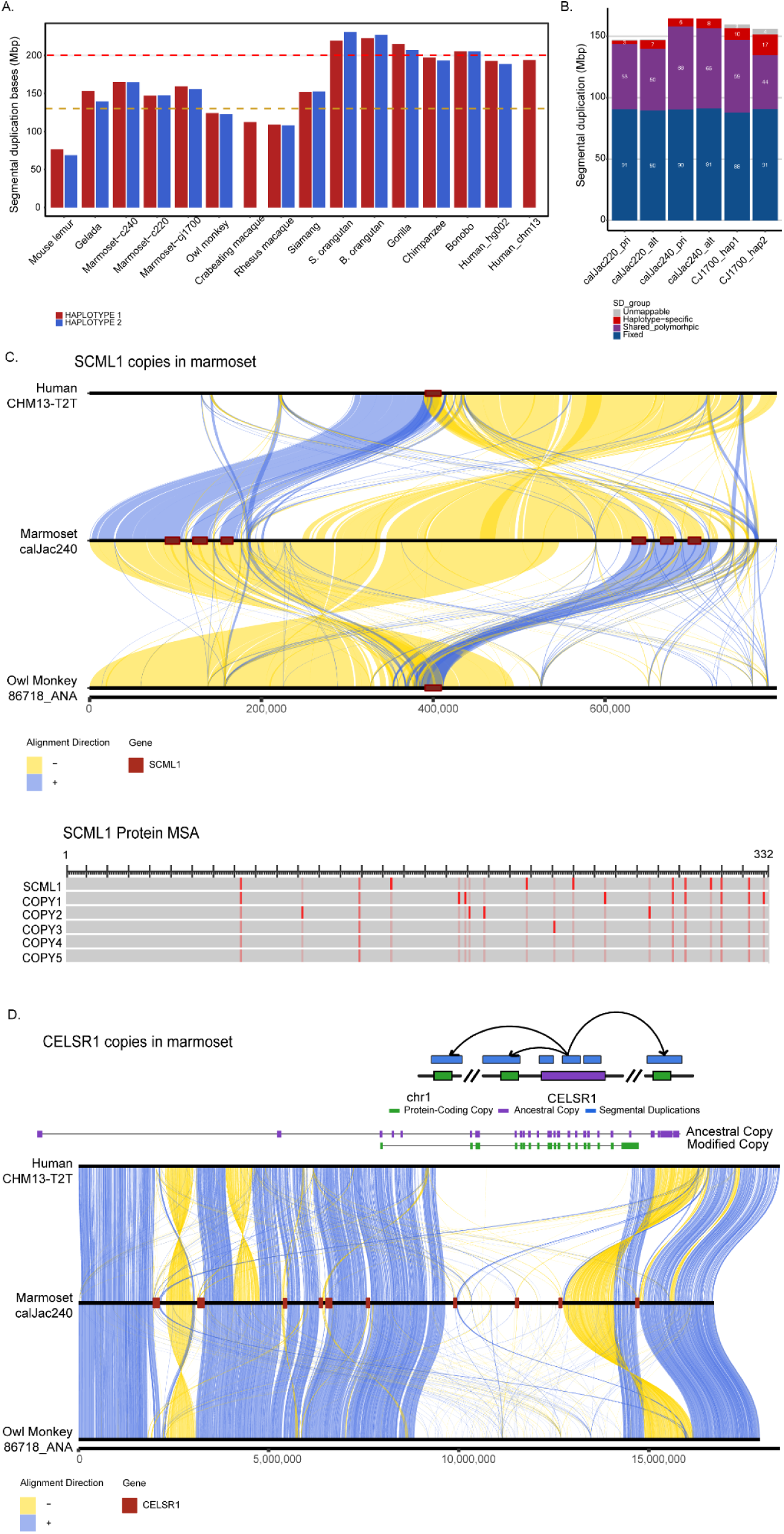
Segmental & Gene duplication landscape in marmoset genomes. (A) Total segmental duplication content (Mbp) across primate species for each haplotype (haplotype 1, red; haplotype 2, blue). Marmosets show intermediate SD content between non-ape primates and apes. (B) Classification of segmental duplications across six marmoset haplotypes by population frequency-fixed duplications present in all haplotypes (blue), shared polymorphic duplications present in more than one but not all haplotypes (purple), haplotype-specific duplications found in a single haplotype (red), and unmappable regions (gray). (C) Duplication and synteny architecture of the *SCML1* gene family on chromosome X. Top: three-way synteny alignment between human (CHM13-T2T), marmoset (calJac240), and owl monkey (86718_ANA) across an approximately 800 kb region. Alignment ribbons indicate forward-strand alignments (blue) and inversions (yellow). *SCML1* gene copies are marked in red on the marmoset chromosome, showing six copies. Bottom: multiple sequence alignment (MSA) of the predicted *SCML1* protein for the ancestral copy and five additional duplicated copies, with red marks indicating amino acid substitutions relative to the ancestral copy. (D) Duplication architecture of the *CELSR1* gene family expansion on chromosome 1. Top: schematic showing the ancestral *CELSR1* locus (purple) flanked by modified protein-coding gene copies (green), with segmental duplication blocks (blue) carrying the modified copies. Cartoon arrows indicate inferred duplication along the chromosome. Middle: gene structure comparison between the ancestral copy (purple) and a representative modified copy (green), illustrating differences in exon-intron architecture. Bottom: three-way synteny alignment between human (CHM13-T2T, top), marmoset (calJac240, middle), and owl monkey (86718_ANA, bottom). Alignment ribbons indicate forward-strand alignments (blue) and inversions (yellow). The *CELSR1* gene loci are marked in red on the marmoset bar.

To characterize SD variability within the species, we analyzed the distribution of duplicated sequences across the six haplotypes (Fig. 3**B**). This analysis revealed that about ∼8.4 Mb/haplotype (3-17 Mb) are haplotype-specific duplications, ∼56.6 Mb/haplotype (44-68 Mb) are polymorphic duplications present in some but not all haplotypes, and ∼90.1 Mb/haplotype (88-91 Mb) of fixed duplications present in all haplotypes. Notably, these duplicated regions encompass 2,754 genes genome-wide, with 921 genes intersecting polymorphic SDs, indicating that segmental duplications contribute substantially to genic variation within the marmoset population.

To identify marmoset-specific duplications, we compared marmoset SDs to those in owl monkey (*Aotus*), a closely related New World monkey species that serves as an outgroup (divergence ∼16.4 MYA). This analysis identified 91-105 Mb of SD sequences specific to the marmoset lineage that are not duplicated in owl monkeys. However, we note that the assembly of owl monkeys is less complete (Fig. **S5**). Functional enrichment analysis indicates that the types of genes expanded through marmoset-specific versus ancestral duplications differ. Genes overlapping marmoset-derived duplications (1,093 genes) were highly enriched for immune system functions (*p*<0.019 but not significant after multiple test correction). In contrast, genes in duplications shared with owl monkeys showed enrichment for metabolic processes (*p*<0.029).

We studied some of these duplicated gene families that have undergone expansion in the marmoset lineage. One example is the *SCML1* (Sex comb on midleg-like 1), a Polycomb group protein involved in transcriptional repression and chromatin remodeling (Fig. 3**C**). In human and owl monkey (outgroup), a single copy of *SCML1* resides on the X chromosome. In contrast, in the marmoset, we identified six copies of *SCML1* distributed across an approximately 2.2 Mb region of chromosome X in calJac240_pri (visualized), 6 in calJac220_pri, and 7 in calJac220_alt. These copies are arranged in a tandem-like cluster embedded within segmentally duplicated blocks. This locus has undergone extensive rearrangement, involving interleaved inversions and duplications, relative to both the human and owl monkey genomes in the common marmoset. Multiple sequence alignment of the predicted protein sequences showed that the duplicated copies have accumulated a few amino acid substitutions relative to the ancestral *SCML1* protein.

Another notable example of lineage-specific gene family expansion is the *CELSR1* gene, which encodes a cadherin EGF LAG seven-pass G-type receptor implicated in brain development (Fig. 3**D**). A single orthologous copy of *CELSR1* resides on chromosome 1. We identified 10 additional modified copies that are duplicated across an approximately 12.7 Mb region on the same chromosome, all distributed within segmentally duplicated blocks in calJac240_pri. This duplication is also confirmed in the other three assembled haplotypes. Most copies consist of segmental duplication blocks carrying protein-coding gene copies that display a modified transcript structure compared to the ancestral locus. Of the ten total loci, nine are annotated as protein-coding genes with intact open reading frames supported by IsoSeq transcripts across at least two tissue types, while one is a pseudogene. The retention of open reading frames in the majority of these modified copies, rather than their degradation into pseudogenes, suggests that these modified *CELSR1* paralogs may be functional and potentially under positive selection. Synteny analysis across human, marmoset, and owl monkey uncovered extensive structural rearrangements in this region in the marmoset, with numerous inversions and duplications relative to both species.

### Marmoset MHC Loci

The Major Histocompatibility Complex (MHC) region encodes highly polymorphic cell-surface proteins that are crucial for antigen presentation and adaptive immunity. In humans, the MHC region is one of the most disease-related and gene-dense regions of the genome^48^. Despite its biomedical importance, especially in a model organism, the highly duplicated and polymorphic nature of the MHC region has made it challenging to assemble and annotate in marmosets. We assembled and annotated the ∼5 Mb MHC region across all four marmoset haplotypes. We employed multiple annotation approaches and integrated them with prior manual annotations from earlier marmoset assemblies to generate consensus annotations for MHC Class I and Class II genes (Table **S10**).

The marmoset MHC Class II region exhibits both conservation and lineage-specific variation compared to other primates. Like humans and great apes, marmoset MHC II includes the canonical gene families-DRA, DRB, DQA, DQB, DOA, DOB, DMA, DMB, DPA, and DPB. However, marmosets show an expansion of the DQA1 and DQB1 loci that is not observed in other primates. The DRB locus, known to exhibit CNV in humans and apes^19,49^, has multiple copies in marmosets, however there is no CNV in these 4 haplotypes. With the sequencing of additional marmoset genomes, the DRB locus may also show CNVs (Fig. **4B**).

**Figure 4.**
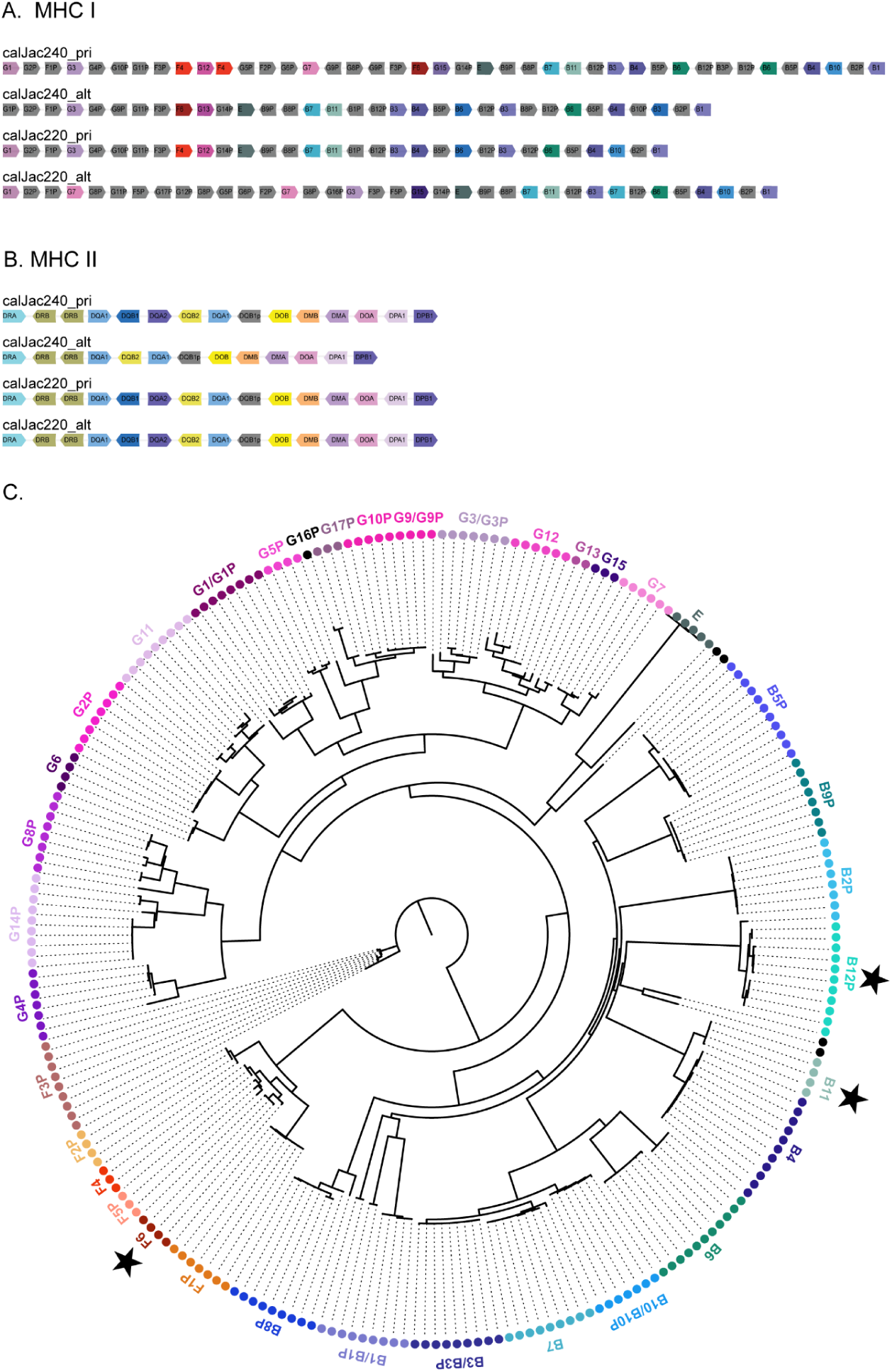
Haplotype-resolved structure of the marmoset MHC. (A) Gene organization of the MHC class I region across four haplotypes. Arrows represent individual genes, color-coded by gene family, with pseudogenes denoted by a “P” suffix. Structural variation is evident between haplotypes. (B) Gene organization of the MHC class II region across the same four haplotypes. Gene order and content are largely conserved, though some haplotype-specific differences are observed. (C) Phylogenetic tree of marmoset MHC class I genes. Tips are color-coded by gene/gene family, with labels indicating gene identity. Black stars denote the novel genes annotated.

The marmoset MHC Class I region exhibits greater structural variation than MHC Class II, with several gene duplications and lineage-specific expansions. Based on previous studies, it is known that marmosets possess MHC I loci orthologous to human HLA-B/C, HLA-G, HLA-F, and HLA-E, designated *Caja*-B, *Caja*-G, *Caja*-F, and *Caja*-E, respectively^50–52^. Copy number variation is particularly prominent in the *Caja*-G and *Caja*-B gene families (Fig. **4A**). Across the four haplotypes analyzed, *Caja*-G shows expansion and variation, with multiple distinct copies identified. Phylogenetic analysis of *Caja*-G sequences shows that there are two major clades, suggesting ancient duplication events followed by independent evolution. The *Caja*-F family is also very diverse, with multiple functional copies distributed across haplotypes. The *Caja*-B locus exhibits similarly high copy number variation. Like *Caja*-G, phylogenetic reconstruction identifies at least two major *Caja*-B clades (Fig. **4C**).

Beyond the established MHC I gene families, our annotations identified several previously unknown MHC Class I genes in the marmoset assemblies. These sequences exhibit structural features characteristic of MHC I genes but diverge significantly from previously catalogued MHC I loci in *C. jacchus*. Whether these represent functional genes, recent duplications that have undergone neofunctionalization, or degrading pseudogenes requires further investigation through expression analysis and functional studies. However, their presence across multiple haplotypes suggests they are not assembly artifacts but rather bona fide genomic features of the marmoset MHC (Fig. **4C**).

### Acrocentric Chromosomes

Marmoset genomes possess 15 acrocentric and subtelocentric chromosomes, as confirmed by both karyotyping and our assemblies. Seven of these contain a short arm that is gene-poor, enriched in satellites, harbor pseudo-homologous regions (PHRs), and can either carry rDNA arrays or remain vacant: 14, 15, 17, 18, 19, 20, and Y (Fig. **5A**, Table **S11**). In addition, the marmoset is the first genome in which we observe that the shared homology extends all the way into the centromeres of each of these chromosomes. Our T2T assembly has complete short arms of these chromosomes, correctly resolving the structure and organization of rDNA array-bearing regions and the satellite sequences that surround them. As in humans, orangutans and gorillas, marmoset rRNA gene arrays are oriented from telomeres to centromeres. The satellite content flanking marmoset rDNA arrays is distinct from that observed in hominids, lacking the DJ region and HSat arrays characteristic of hominid acrocentric short arms. We performed fluorescence in situ hybridization (FISH) analysis on mitotic chromosome spreads using rDNA-specific probes together with chromosome-specific identification markers to detect and measure the fluorescence intensity of each rDNA array and assign it to the corresponding chromosome. rDNA arrays were present on chromosomes 15, 17, 18, 19, and Y in calJac240, and on chromosomes 14, 15, 17, 18, and 19 in calJac220. Using the total rDNA copy number estimate obtained from short-read sequencing data and the fractional fluorescent intensity measurements per array, we calculated the number of rRNA gene units across both haplotypes^53^ (Fig. **5B**, Fig. **S6**).

**Figure 5.**
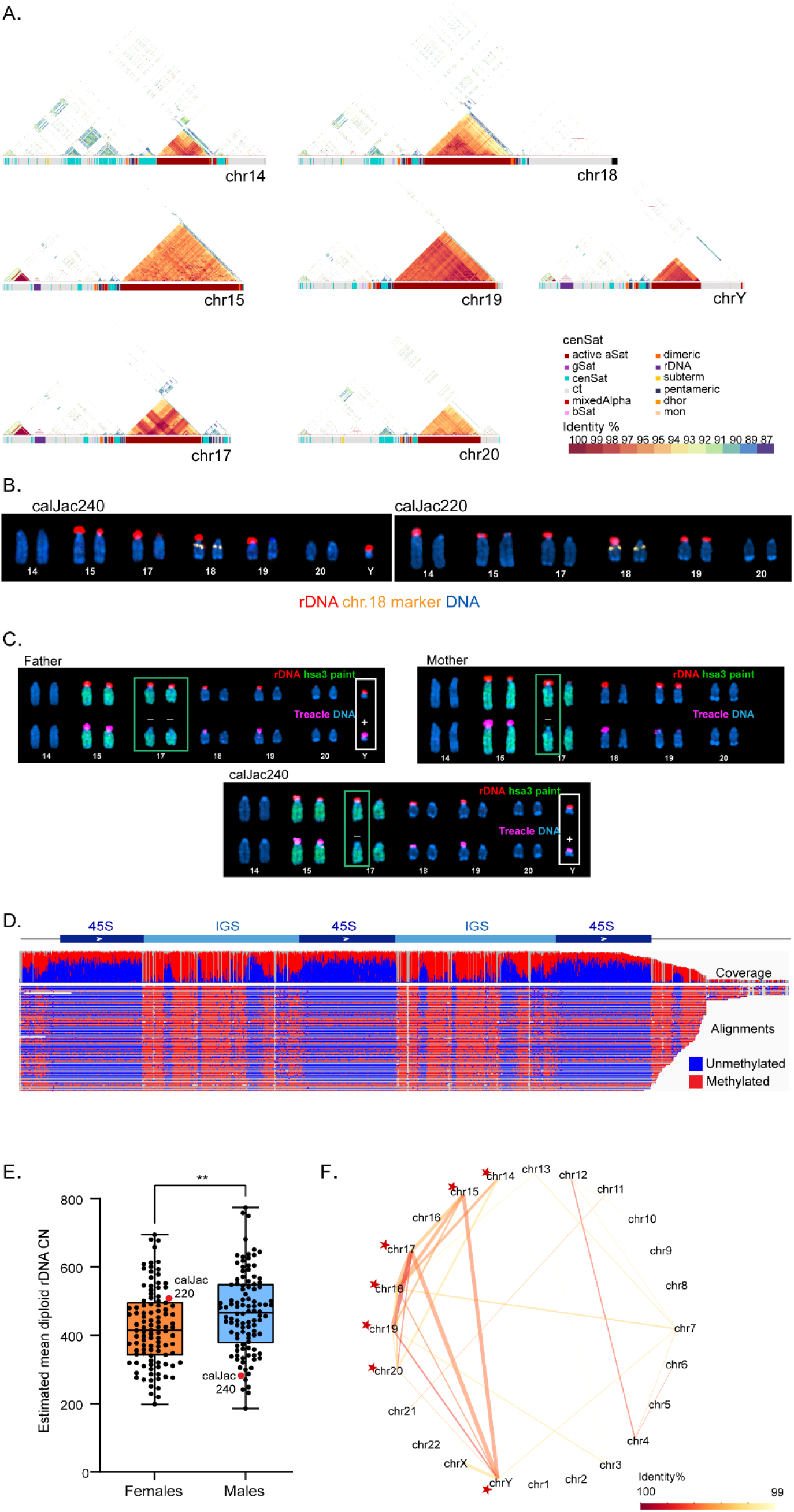
Acrocentric chromosomes in the marmoset. (A) Structure of acrocentric chromosome short arms in the marmoset T2T assembly. Ideograms show the short arm of chromosomes 14, 15, 17, 18, 19, 20, and Y with satellite annotations color-coded by class. Triangular self-identity heatmaps display sequence identity within each short-arm region, with identity percentages indicated by the color scale (87-100%). (B) Acrocentric karyograms from a representative metaphase chromosome spread from calJac240 and calJac220 labeled by FISH with rDNA probe (red), chromosome 18 marker (yellow), and DNA (blue). (C) Immuno-FISH analysis across a marmoset trio (father, mother, and calJac240). The top rows of chromosomes show FISH labeling with rDNA probe (red) and hsa3 whole chromosome paint (green). The bottom rows show the corresponding chromosomes labeled with the Treacle antibody (magenta). DNA was counter-stained with DAPI. Green boxes highlight chromosome 17, white boxes highlight the Y chromosome. Minus signs (-) indicate Treacle-negative (transcriptionally silent) rDNA arrays on chromosome 17, and plus signs (+) indicate Treacle-positive (transcriptionally active) arrays on chromosome Y transmitted to offspring. (D) Methylation profile of rDNA units on chromosome Y showing the 45S and intergenic spacer (IGS) regions. Coverage track (top) and individual read alignments (bottom) are colored by methylation status (blue, unmethylated; red, methylated). (E) Distribution of estimated mean diploid rDNA copy number across marmoset individuals, separated by sex. Females (orange, left) and males (blue, right) show significantly different copy numbers (**, p < 0.01). Individual data points for calJac220 and calJac240 are indicated. (F) Network diagram of PHRs across marmoset chromosomes in the style of Solar et al. ^59^. Nodes represent chromosomes, with acrocentrics marked by red stars. Edges connect chromosomes sharing homologous sequences, with line color indicating identity percentage (scale at right, 99-100%) and line thickness reflecting the amount of shared sequence.

Both calJac240 and calJac220 displayed individual-specific patterns in rDNA copy-number distribution and total copy number, indicating that rRNA gene array profiles are unique to individuals. A notable feature of the marmoset genome is the heterogeneity in the presence or absence of rDNA on acrocentric chromosomes, both between chromosomes and between haplotypes within the same chromosome. The short arms of these chromosomes can either bear rDNA arrays or remain vacant, instead containing other heterochromatic and satellite sequences. Some haplotypes or individuals have rDNA on specific chromosomes, while others lack it. For example, chromosome 14 bears an rDNA array in calJac220 with approximately 141 copies on one haplotype, yet showed no detectable rDNA in calJac240 (Fig. **5A**, chr14 in calJac240). In calJac240, only one haplotype of chromosome 18 and only one of chromosome 19 bear rDNA, indicating differences in rDNA array distribution within haplotypes of a specific chromosome in an individual (Fig. **5B** left). For calJac240, cell lines derived from the father, mother, and sister were also available and were used to assess the transmission patterns of acrocentric p-arms cytogenetically. Several haplotypes could be traced from parents to offspring. For instance, calJac240 inherited a small rDNA array on chromosome 17 from the mother, an rDNA-less haplotype of chromosome 19 from the father, and inherited a chromosome Y rDNA array of comparable size to that in the father. Additionally, each parent carried one copy of an rDNA-less chromosome 18, and calJac240 inherited one of these copies. In contrast, the sister inherited both rDNA-less copies of chromosome 18, demonstrating how each individual acquires a distinctive distribution of rDNA gene arrays, and thus a unique total rDNA copy number (Fig **5C**, Fig. **S7**)

In the rRNA gene unit, the promoter and the coding region generally exist in two distinct epigenetic states: unmethylated and methylated, while the intergenic spacer (IGS) remains methylated in both states^54^. The promoter and the coding region of active genes are bound by RNA Polymerase I transcription factors such as UBF and Treacle. This can serve as a marker for rRNA gene activity and be assessed by immuno-FISH^53^. Colocalization of rDNA FISH probe signal and the Treacle antibody signal indicates an active rDNA array, whereas rDNA FISH signal without Treacle indicates a transcriptionally silent rDNA array. Immunostaining for Treacle confirmed the transcriptional activity of some identified rDNA arrays (Fig. **5C**), whereas several arrays lacked Treacle, indicating they are transcriptionally silent. These findings are in line with previously reported silent rDNA arrays in humans and apes^25,53^.

Analysis of the trio (father, mother, and calJac240) enabled tracing of transmission of rDNA activity status across a generation. On chromosome 17, calJac240 inherited transcriptionally silent rDNA arrays from both parents, a large array from its father and small one from its mother, as confirmed by the absence of Treacle signal on both haplotypes despite the presence of rDNA copies (Fig. **5C**). These silent arrays were also inactive in the parents, suggesting that the epigenetic silencing state of rDNA arrays can be inherited across generations. The high sequence similarity among rDNA copies across the short arms of somatic acrocentric chromosomes and haplotypes rendered the assignment of short reads to DNA impractical, precluding analysis of their methylation status. However, immuno-FISH assays using Treacle as a marker of active NORs provided a direct, chromosome-specific readout of transcriptional activity, allowing detection of silent rDNA arrays. This provides evidence of the potential inheritance of a silent rDNA state in a non-human primate, paralleling recent observations in human trios^55,56^.

Notably, the Y chromosome in calJac240 carries an rDNA array of approximately 38 copies. Y-linked rDNA arrays have been observed in several other primate species, including orangutans and siamang^25^, as well as in non-primate mammals such as dogs^57^ and the lesser mouse-deer (*Tragulus javanicus*)^58^. Long reads from the Y-linked rDNA array spanning neighboring regions could be assigned unambiguously, enabling methylation analysis of the rDNA array on this chromosome. Based on the low methylation status of the coding regions of Y-linked rRNA genes and the positive Treacle signal by immunostaining, the rDNA array on the Y chromosome is transcriptionally active in calJac240 and the father (Fig. **5C**, **5D**). The Y-linked rDNA array could create a sex difference in rRNA gene copy number, though the functional implications of this remain unclear. Comparison of total diploid rDNA copy number across 211 marmoset samples, including our two sequenced individuals, revealed sex-specific differences. Males showed significantly higher mean diploid rDNA copy numbers compared to females (p < 0.01), likely reflecting the contribution of the Y chromosome rRNA gene copies. Both individuals sequenced in this study fell within the normal range of variation, with calJac240 (male, ∼282 total copies) slightly below the male median and calJac220 (female, ∼508 total copies) slightly higher than the female median (Fig. **5E**).

Recent analyses of human and great ape T2T genomes have shown that there are pseudo-homologous regions (PHRs) shared between the short arms of acrocentric chromosomes. These are regions sharing >99% sequence identity that undergo active recombination despite residing on heterologous chromosomes^59^. To investigate whether similar PHRs exist in marmosets, we analyzed sequence similarity across all acrocentric short arms using a minimum threshold of 99% identity over 10 kb spans. We see specific patterns of sequence sharing between acrocentric pairs, with the strongest PHR relationships observed between chromosomes 17 and 19 (Fig. **5F**). The pattern of PHRs in marmosets suggests that chromosomes 14, 15, 17, 18, 19, 20, and Y can exchange genetic information. The two marmoset genomes in this study lack rDNA on chromosome 20. Still, given the heterogeneity in rDNA arrays among haplotypes documented here and the presence of the PHR, we speculate that chromosome 20 may contain rDNA in some individuals.

We also identified an unexpected partially degraded rDNA array in the pericentromeric region of metacentric chromosome 6, present in all four assembled haplotypes. Unlike the canonical rDNA arrays, this array contains rDNA units disrupted by Alu and LINE element insertions within the rDNA gene unit (Fig. **S8**). The presence of retrotransposon insertions within rDNA units is inconsistent with active ribosomal RNA transcription, implying that this array is functionally inactive^60^. A similar configuration has been observed in giraffe^61^, though comparable observations have not, to our knowledge, been reported in T2T-assembled primate genomes. The chromosome 6 array may represent an ancestral translocation or duplication event that subsequently degraded. Such arrays may not be detectable by conventional FISH, highlighting the utility of T2T assembly for identifying degenerate rDNA loci.

### Centromeres & Satellites

#### Centromeric Satellites

Common marmoset centromeres are composed of alpha satellites organized into relatively compact arrays, similar in size to apes but much smaller than in Old World monkeys (OWM)^19,20^ (Fig. **S9**, Note **S1**). The average size of an active array is ∼1.2 Mb, about 2 times less than that of humans^22^ and about 8 times less than that of macaques^20^. Like apes and unlike macaques, marmoset centromeres contain far fewer inversions within active arrays^19,20,22,62^. A comprehensive NWM alpha-satellite classification and HMMER-based annotation tool was built using methods previously described for OWM^20^, and we annotated the centromeric satellites, including the active alpha-satellite arrays across the assemblies (Note **S1**). We identified CDRs in these active alpha-satellite arrays, suggesting these are candidate sites for kinetochore assembly. Consistent with previous data for NWM^23,24^, marmoset active alpha satellites are dimeric and lack HORs, with one notable exception we discovered on the X chromosome (Note **S1**). The active dimeric repeats are formed by two major ∼170 bp monomer types, S3 and S4 (NWM-SF1). The older generations of these dimers were found to form small, sometimes symmetrical inactive arrays on centromere flanks (SF2, SF3, and SF4, respectively), and they represent the remnants of the centromeres of ancestors of marmosets and other NWM (Fig. **6A**, Note **S1**). The two older pericentromeric alpha satellite families (NWM-SF 5 and 6), reconstructed from only a few short relic arrays, likely represent the centromeres of entirely extinct taxa that split from NWM shortly after divergence from the common OWM-ape lineage. Upon comparing NWM-specific dead layers with ape/OWM shared layers, and those shared only between OWM and apes, we find that the last common ancestor of all three branches had APE-SF11 centromeres (Note **S1**).

**Figure 6.**
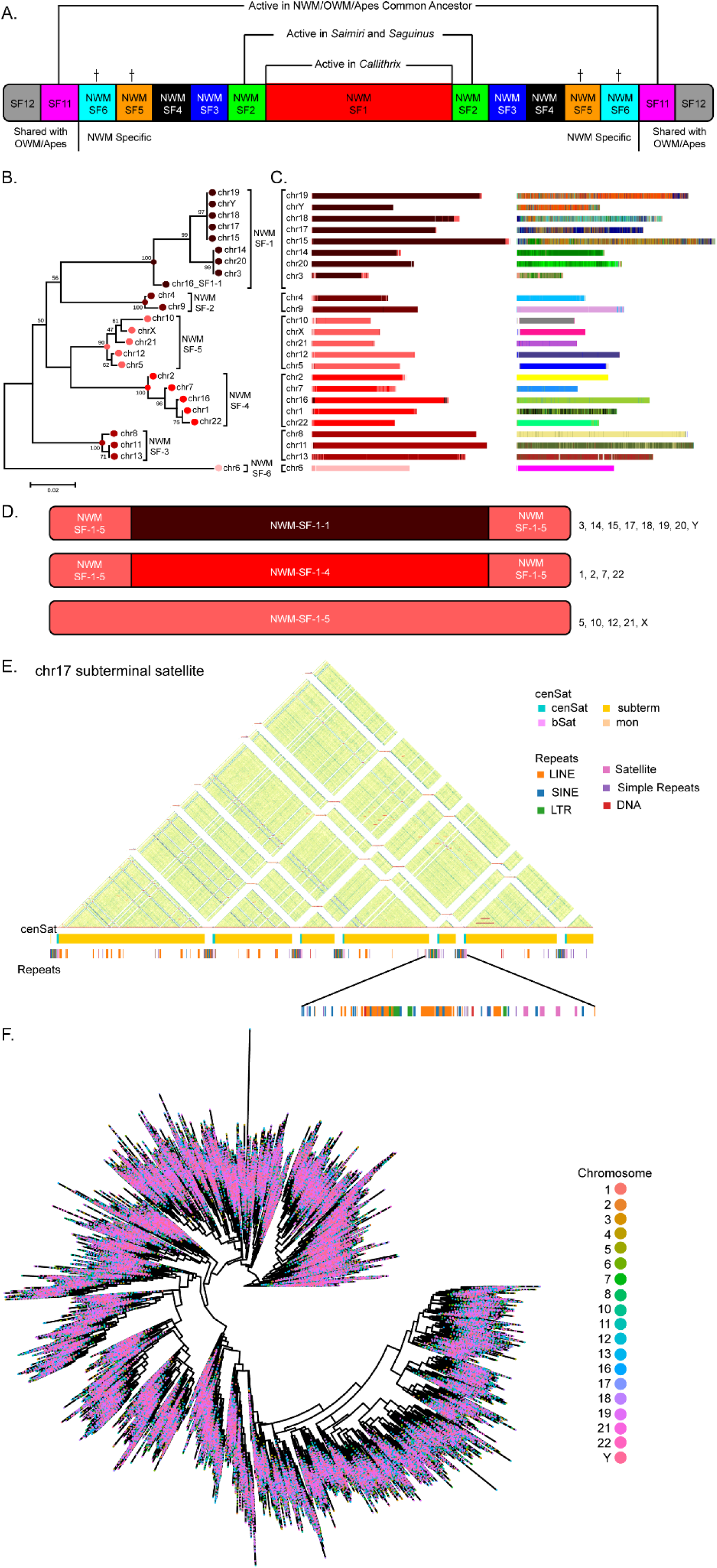
Characterization of centromeric and subterminal satellite DNA across marmoset haplotypes. (A) Reconstructed generalized marmoset centromere (not to scale) with its active (red) and inactive satellite layers and their relationship to the NWM species as established in this paper. The oldest NWM-specific layers (NWM-SF 5 and 6, marked by crosses) correspond to entirely extinct species. In fact, they may not represent two independent families, and their relative order could not be established reliably due to their rarity (Note **S1**). The unmarked ones may yet be found as active centromeres when T2T assemblies of species from NWM branches other than Cebidae will become available. SF11 (shared with ape/OWM lineage) was ancestral to all NWM-specific layers (Note **S1**). (B) Minimum evolution phylogenetic tree of alpha satellite consensus S3S4 dimers derived from 500 random monomers extracted from active arrays of each marmoset centromeric satellite array. For chromosome 16, 151 SF1-1 monomers from two flanking inactive arrays were also included. Branches corresponding to subSFs within NWM-SF1 are marked. (C) LEFT: The centromeric satellites of calJac240 assembly annotated with the NWM-HMMER-subSF tool. Multiple cases of alpha satellite interlayers, where active arrays of one subSF are flanked by small arrays of another subSF, indicate that a recent change in centromeric satellite identity can be observed. Chromosome 16, an acrocentric chromosome not covered by SF1-1, retains remnants of the old SF1-1 centromere (black) on both sides of the active array. RIGHT: The same complement of the centromeric satellite arrays as in C, but annotated for chromosome-specific dimhaps. Note that outside of SF1-1, all centromeres are chromosome-specific. (D) Competition of subSFs within active SF1-1 in marmoset centromeres is shown using SF1-5 (pink) as an example. Originally, it was the most numerous family and covered 17 centromeres, however only 5 (1, 10, 12, 21, and X) have retained SF1-5 active regions to this day. In acrocentrics and chromosome 3, it has been replaced by SF1-1 (dark brown), and in chromosomes 1, 2, 7, and 22, by SF1-4 (red). (E) Self-identity heatmap of a representative subtelomeric satellite region on chromosome 17, showing highly organized higher-order repeat arrays interspersed with non-satellite regions. Tracks below indicate the distribution of centromeric satellites (cenSat) and various repeat elements (LINE, SINE, LTR). (F) Unrooted Neighbor-Joining phylogenetic tree using 140,316 full-length subterminal satellite (MarmoSat) monomers derived from calJac240 T2T assembly, showing a lack of chromosomal or array-specific homogenization.

The active NWM-SF1 dimers can be further subdivided into six major groups (subSFs) distributed across different chromosomes. The most prevalent SF1-1 dimer (S3-1, S4-1) is present on chromosomes 3, 14, 15, 17, 18, 19, 20, Y (all acrocentric/subtelocentric, and all containing satellites & PHRs in their short arm, except 3). In contrast, the rarest SF1-6 dimer (S3-6, S4-6) is unique to chromosome 6 (Fig. **6B**, **6C**).

With further analysis, we also discovered the history of competition between subSFs, with SF1-5 losing most of its original occupancy to SF1-1 (found in 7 acrocentric chromosomes) and SF1-4 (found in chromosomes 1, 2, and 22) (Fig. **6D**, Note **S1**). Contrary to expectations^24^, high-copy dimers exhibit specificity for individual centromeres, enabling the identification of centromere-specific dimer haplotypes (dimhaps) and the development of chromosome-specific k-mer signatures for most centromeres. We also studied marmoset centromere evolution in detail (Fig. **6C** (right)), using an additional chromosome-specific module of the NWM annotation tool that we built. NTRprism kmer periodicity analysis^22^ of all active arrays in the T2T assembly confirmed a predominant dimeric repeat structure with some tetrameric repeats. The X chromosome uniquely contains higher-order repeats, with the most common being a 28-mer. Thus, the chromosomal specificity of centromeric satellites in marmosets is not based on uniquely structured HORs (like in apes), but relies almost solely on dimhaps (like in OWM). A comparative study of calJac240 and calJac220 assemblies showed considerable intraspecies diversity, with distinct centromeric haplotypes across several chromosomes (Note **S1**).

Interestingly, the subSF1-1 was originally shared as the active centromere among only acrocentric chromosomes (3, 14-20, and Y). Among the confirmed stably rDNA array-bearing acrocentrics (15, 17, 18, 19, and Y), chromosomes 19, Y, 17, and 15 harbor the most closely related dimer sequences (Fig. **6B**, **6C**), with chromosomes 18, 14, and 20 also falling within the SF1-1 clade. Chromosome 20, which retains an active SF1-1 centromere with dimers closely related to those of chromosome 14, lacks an rDNA array in the four haplotypes assembled here. But, this may represent unsampled polymorphism, consistent with the variable rDNA array presence documented on chromosome 14. Chromosome 16 is an acrocentric chromosome by morphology, with short arms that are not satellite-rich or PHR-containing.

The active SF1-1 centromere on this chromosome has been replaced, and it now carries an SF1-4 active array. However, SF1-1 arrays persist on both flanks as inactive remnants of its former centromere (designated chr16_SF1-1 in Fig. **6B**). Without additional genomic information from closely related species, we cannot infer how chromosome 16 arrived at its current state. It may have either once participated in the network of chromosomes sharing alpha satellite, rDNA, and PHRs and was later excluded, or alternatively, it may represent an ancestral source of the SF1-1 sequence that donated it to other chromosomes without ever having carried rDNA and PHRs itself.

#### Subterminal satellite arrays

Marmosets possess satellite arrays positioned near almost all chromosome ends, with a few exceptions (Fig. 1B), confirming observations from previous studies^63^. These subterminal structures occupy 1.5-1.8% of each haplotype assembly, up from the previous estimate of 1.09%^63^. Using NTRprism analysis, we were able to confirm that these arrays are composed of a characteristic 170/171 bp monomer unit that is conserved across all arrays (Fig. **6E**). The subterminal satellite DNA (MarmoSAT) and alpha satellite repeat units are similar in size (∼171 bp). Despite the same motif size, MarmoSAT and alpha satellites do not share any nucleotide characteristics or conserved secondary structure that suggests a common origin.

Structurally, marmoset subterminal arrays parallel those recently described in great apes and gibbons, featuring interspersed non-satellite regions enriched for transposable elements. However, the sequence composition differs both in monomer size and content. To ascertain the homogenization dynamics of the subterminal satellite arrays related to their chromosomal origin, we mined all monomers from calJac240. An NJ phylogenetic tree with 140,316 full-length copies of the subterminal satellite revealed no differential homogenization of specific variants within subtelomeric arrays (Fig. **6F**). Our results suggest the presence of homologous and ectopic recombination through persistent subterminal associations.

### Sex Chromosomes

The sex chromosomes have historically been among the most incomplete regions of primate genome assemblies due to their high repeat content, ampliconic regions, and heterochromatic sequences^25^. Our T2T assembly added approximately 4.1 Mb of new sequence to the X chromosome and approximately 6.0 Mb to the Y chromosome from calJac4, which is approximately 40% of the chromosome length (15.8 Mb). These sequences are enriched for repetitive content on the Y chromosome; satellite sequences account for 3.6 Mb, and segmental duplications for 2.7 Mb of newly resolved bases (Fig. **7A**, **7B**). The most Y chromosome sequence assembled before this was in mCalJac1 (*GCA_011100555.2*)^12^, which reported 13.84 Mb of Y-linked sequence, but the main Y chromosome scaffold (*NC_071465.1*) in that study is 4.9 Mb long.

The assembly of the Y chromosome reveals a complex arrangement of distinct sequence classes. Ancestral regions, which retain detectable homology with the X chromosome, are interspersed with large ampliconic blocks containing multi-copy gene families. Analysis of the Y chromosome highlights palindromic structures, high-identity duplications, and subterminal satellites (Fig. **7B**).

The marmoset Y chromosome retains some ampliconic gene families shared with humans, including *CDY*, *DAZ*, *TSPY*, and *RBMY*, confirming that these amplifications predate the divergence of New World and Old World primates. In contrast, *BPY2* and *PRY* are absent in marmosets, emerging only in African apes. At the same time, *HSFY* and *VCY* are absent in marmosets but present in Old World primates, indicating that these gene families arose after the New World/Old World primate split^25^. We were able to identify three ampliconic gene families on the marmoset Y chromosome-*DDX3Y*, *EIF1AY*, and *SHOX*, that exist as single-copy genes in the human Y chromosome, confirming previous findings^12^. These expansions seem to be specific to the marmoset lineage and represent independently derived amplifications not shared with great apes. *DDX3Y* encodes an RNA helicase essential for spermatogenesis and is present as a single-copy gene on the male-specific region of the Y chromosome in humans. In the marmoset, we identified a total of nine copies. Upon aligning all copies to the orthologous gene model, we find that most duplicates are highly similar (Fig. **7C**). Iso-Seq long-read transcriptome data indicate that at least six of the eight duplicated copies are transcriptionally active across multiple tissues, while one shows limited expression and one lacks detectable Iso-Seq support, suggesting it may be a pseudogene. The functional significance of this expansion, whether it reflects selection for increased dosage of *DDX3Y* in marmoset spermatogenesis or has acquired novel regulatory or functional roles, warrants further investigation. Similarly, *EIF1AY* is a single-copy gene on the male-specific region of the human Y chromosome that encodes a protein related to translation initiation. In the marmoset, we identified five extra copies with fully intact ORFs, though none appear transcriptionally active based on Iso-Seq data. *SHOX* is a homeobox gene that controls the formation of many body structures. It is present in a single copy in the pseudoautosomal region in humans and great apes. Marmosets harbor three additional copies; however, these have pseudogenized, as they lack an intact ORF and transcriptional support.

**Figure 6.**
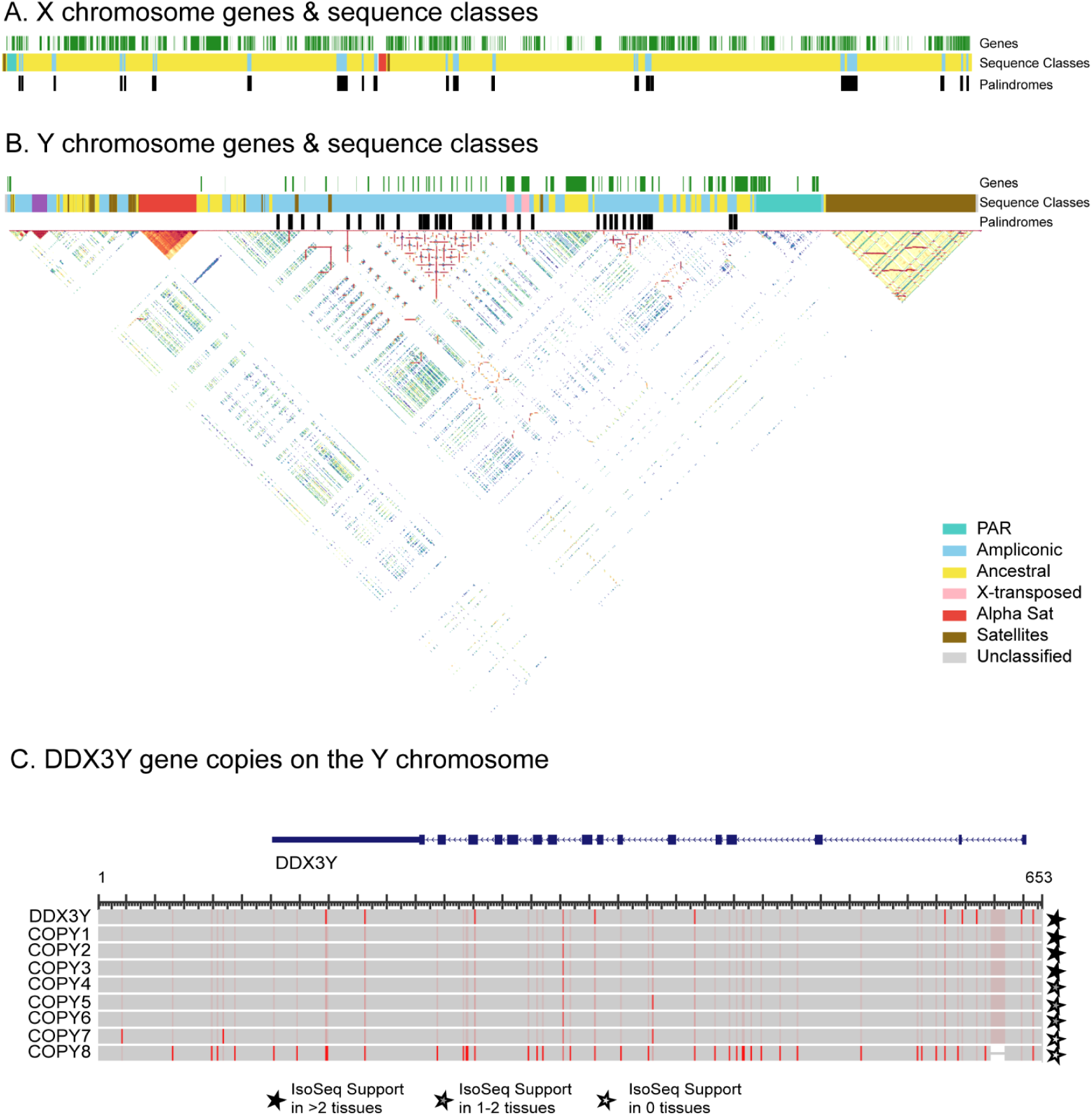
T2T assembly of the marmoset sex chromosomes. (A) Structure of the X chromosome, showing gene annotations, sequence class assignments, and palindromic sequences in the style of Makova et al^25^. Sequence classes are colored as described in the legend below. (B) Structure of the Y chromosome with the same annotation tracks as in (A), overlaid with a triangular self-identity heatmap. Sequence classes include pseudoautosomal regions (PAR, green), ampliconic sequences (blue), ancestral regions (yellow), X-transposed (only on Y) (pink), alpha satellite arrays (orange), other satellite sequences (brown), and unclassified regions (grey). Large palindromic structures and ampliconic duplications are visible as high-identity blocks along the diagonal and off-diagonal. (C) Amplification of *DDX3Y* on the Y chromosome. The ancestral copy (*DDX3Y*, top) is shown with its full gene model (exons as filled blocks). The proteins of the ANC copy, along with the additional copies, are aligned below, with red colouring indicating sequence variants relative to the ancestral copy. Filled, half-filled, and open stars indicate Iso-Seq transcriptional support in more than two tissues, one to two tissues, or no tissues, respectively. Copies 1-6 show broad transcriptional support, while copies 7 and 8 show reduced or no evidence of expression.

## Discussion

The telomere-to-telomere marmoset genome assemblies presented here represent a substantial advance in genomic resources for a key biomedical model organism. These assemblies provide the first complete view of a New World monkey genome. Together with a multispecies primate alignment, a marmoset pangenome graph, and gene annotations incorporating long-read transcriptomic data, these resources establish a comprehensive foundation for leveraging the marmoset in translational research and comparative genomics.

The identification of novel transcript isoforms, including new splice forms of neurodevelopmentally relevant genes, demonstrates the translational relevance of improved transcript annotation in this species, as incomplete references not only obscure gene content but mask isoform diversity relevant to disease modeling. The adoption of the calJac240 T2T assembly as the NCBI RefSeq and UCSC Genome Browser reference will ensure that future marmoset studies benefit from these improvements.

The MHC Class I and Class II variation in marmosets likely reflects long-term balancing selection that maintains immune diversity in response to pathogen pressures, similar to patterns documented in humans and great apes^64^. The high degree of copy-number variation observed between haplotypes within individuals and between individuals, based on preliminary analysis, suggests that MHC diversity in marmosets is maintained through gene duplication and structural rearrangement. We confirm the presence of MHC I genes across multiple haplotypes that are divergent from all previously known common marmoset MHC class I loci. This warrants functional investigation to determine whether they represent neofunctionalized duplications, degrading pseudogenes, or adaptations to meet the demands of serving as ligands for polymorphic receptors of natural killer cells.

The marmoset genome has undergone dynamic remodeling via segmental duplication, with implications for lineage-specific adaptation across a broad spectrum of biological functions. Many gene family amplifications occur within regions of extensive structural rearrangements characterized by inversions and duplications, as in other primate genomes. The enrichment of marmoset-specific segmental duplications for immune-related functions, in contrast to the enrichment of ancestrally shared SDs for metabolic processes, suggests that lineage-specific duplication events have preferentially targeted immune gene families in the marmoset lineage. This pattern, along with the marmoset-specific expansions of the *Caja*-G/*Caja*-B loci, may reflect adaptation to distinct pathogen pressures or the particular ecological niches occupied by marmosets. The duplication of gene families implicated in reproduction and development further points to selective pressures unique to *Callitrichids*, who are characterized by distinctive traits, including obligate dizygotic twinning, chimerism^65,66^, and reduced body size^67^. These duplications may have facilitated the evolution of the hormonal and developmental pathways underlying such specializations. Beyond immune and reproductive functions, the marmoset genome harbors expansions across diverse gene families, including those involved in cytoskeletal organization, cell cycle regulation, nuclear transport, skeletal development, and DNA damage response, among others. Several of these expanded families have given rise to novel transcript structures, suggesting that duplication has been accompanied by regulatory and functional diversification rather than copy number increase alone. The retention of intact open reading frames across multiple copies in several expanded gene families suggests that at least a subset of these duplications may be functionally relevant rather than neutral. These observations underscore the role of duplications and inversions in driving genic innovation in the marmoset lineage and shaping adaptation across immunity, reproduction, and a wide range of cellular processes.

We also provide a comprehensive classification and annotation of New World monkey centromeric satellite arrays. Centromeric regions in the marmoset exhibit a mosaic of features-their compact arrays and absence of large-scale inversions within active regions resemble those of ape centromeres, while their dimeric active-repeat organization and limited chromosomal specificity (in terms of structural diversity) are shared with Old World monkeys. The stratified architecture aligns with the predictions of the expanding centromere/layered expansion model^22,68^. The rapid centromere divergence observed between closely related *Callitrichid* species (Note **S1**) and the considerable intraspecies diversity between our two studied individuals highlight the exceptional evolutionary turnover of centromeric sequences in this lineage.

The resolution of common marmoset acrocentric short arms shows a degree of variability in rDNA array-bearing chromosomes that has not been appreciated in primates so far-the gain and loss of rDNA arrays on individual chromosomes, as opposed to the well-documented variation in rDNA copy number between homologs. This is consistent with evidence that primate genomes maintain substantial excess rDNA capacity. In orangutans, multiple arrays can be transcriptionally silent^53^, and in humans, about half of rRNA genes are silent as inferred from methylation patterns^54^. Since rRNA genes are typically present in excess, some acrocentric chromosomes can lack rDNA arrays, and the individual can still have a sufficient overall copy number.

The chromosomal positions harboring rDNA show conservation across primates-the short arms of acrocentric chromosomes, even as the number of rDNA array-bearing chromosomes is not, up to nine rDNA-bearing pairs in orangutans^19^ and at least six pairs in marmosets. This conservation of rDNA within acrocentric short arms holds for apes and New World monkeys, both of which harbor rDNA arrays on multiple acrocentric pairs. Old World monkeys differ substantially, at least in currently well-assembled species, in that they maintain rDNA on only a single chromosome pair. The shared multi-chromosome rDNA arrangement in New World monkeys and apes may represent the ancestral state, whereas the Old World monkey configuration may reflect a lineage-specific one. Given the different inter-individual configurations documented here, the ancestral rDNA array bearing complement may have been larger, with independent losses accumulating in each lineage.

Our trio analysis provides evidence that the epigenetic state of rDNA arrays can be transmitted across generations, along with size. However, we are unable to confirm this through methylation analyses because we do not yet have assemblies for the parental samples. We were also able to confirm, through cytogenetics and methylation data, the existence of an active Y-linked array and, through cytogenetics, its inheritance. The functional consequences of such an array remain unclear, but the implications may extend beyond rRNA dosage. The marmoset’s short generation time and routine twinning make it a particularly tractable system for studying rDNA epigenetics, including the inheritance and functional consequences of the Y-linked array.

We also find that all chromosomes sharing a PHR also share similar centromeric DNA sequences. This is consistent with the pattern observed across primate lineages, in which the centromeres of rDNA array-bearing acrocentric chromosomes belong to the same alpha satellite superfamily. In marmosets, all potentially rDNA array-bearing acrocentrics share SF1-1, with several harboring near-identical dimers. In humans and gorillas, all rDNA array-bearing chromosomes have APE-SF2 centromeres; in chimpanzees, all have undergone coordinated replacement to APE-SF1. Such consistency across independently evolving lineages would hardly be expected under neutral centromere evolution.

We propose that this pattern reflects that shared centromeric sequences are maintained by selection because they facilitate the exchange of heterologous short arms. During interphase, acrocentric short arms co-localize within the nucleolus, bringing rDNA arrays on heterologous chromosomes into physical proximity and creating conditions favorable for ectopic recombination. This could account for the observed sequence sharing between heterologous acrocentric short arms and may provide a mechanism for both the gain and loss of rDNA arrays on individual chromosomes. PHRs are not merely a consequence of shared rDNA sequence but may also actively facilitate the inter-chromosomal exchanges that drive differences in rDNA array presence and copy number. When rDNA array polymorphism reduces copy number below a functional threshold, recombination between acrocentrics with similar centromeric sequences could redistribute rDNA arrays and restore adequate copy number. The gene-poor, repeat-rich architecture of acrocentric short arms minimizes the dosage consequences of such exchanges, while shared repetitive sequences, including satellites and SST1 repeat arrays, provide substrates for recombination.

Several lines of evidence support this model. Hybrid centromeric arrays combining sequences from different acrocentrics have been documented^69^, indicating that inter-chromosomal exchange does occur, and shared centromeric identity may increase the probability of such events. In humans, the strongest PHR relationships occur between chromosomes 13 and 21, which share nearly identical centromeric sequences. Another strong PHR relationship is between chromosomes 14 and 22, which also share a high degree of centromeric identity^59^. Our results in marmosets are consistent with this. All acrocentric chromosomes originally shared SF1-1 as their active centromere, and all confirmed rDNA array-bearing acrocentrics retain it.

This framework may also account for the delayed Y centromere phenomenon. In species where the Y lacks an rDNA array (humans, chimpanzees, Old World monkeys), its centromere retains ancient alpha satellite families absent from autosomal centromeres, consistent with exclusion from the interchromosomal exchanges that homogenize centromeric sequences^19,20,25^ (Note **S1**). In species where the Y bears an rDNA array (orangutans, gibbons, marmosets), its centromere belongs to the same satellite family as that of other rDNA array-bearing acrocentric autosomes. In these species, the Y chromosome lacks a pseudoautosomal region (PAR) on the same arm as the rDNA array, whereas in humans and bonobos, where there is no rDNA array on the Y, PARs are present on both arms. However, this does not hold for chimpanzees which do not have a PAR on both arms. The gorilla Y chromosome, which carries a current-generation centromere but lacks an rDNA array, may represent either a recent rDNA array loss or a case in which the “delayed Y” trend was broken purely by stochastic factors.

The marmoset subterminal satellite arrays architecturally parallel those of great apes and gibbons but differ in sequence composition, indicating independent evolutionary origins. Satellite DNA repeats on the same array or chromosome usually show a higher level of sequence similarity to each other when compared to non-homologous chromosomes. Our results, however, suggest the presence of persistent homologous and ectopic recombination through the subterminal satellite DNA arrays (MarmoSAT), resulting in the lack of array-or chromosomal-specific homogenization, resembling the pattern observed in the gorilla subterminal heterochromatic caps^70^. We hypothesize that the meeting of subterminal regions from non-homologous chromosomes facilitates ectopic pairing and DNA exchange in the first meiotic prophase, generating variability and genomic dispersion of subterminal arrays^70,71^. Therefore, an increase in subterminal satellite blocks likely leads to higher ectopic recombination rates, thereby creating an incubator for genomic diversity within subtelomeric regions^70^. The recent origin in the common ancestor of *Callitrichini* and *Callimico goeldi* (∼8 Mya)^63^ and the expansion of MarmoSAT subterminal arrays suggest that this repeat may act as an underlying engine of chromosomal differentiation.

Despite the significant findings presented here, some limitations should be noted. These assemblies are derived from only two individuals. Broader population-level sequencing and assembly will be needed to fully characterize genomic diversity within the species. Many of the findings reported here will require experimental validation to determine their functional significance. Additionally, the quality of comparative analyses is inherently constrained by the completeness of available assemblies from related species. As sequencing technologies and assembly methods continue to improve and more individuals and species are sequenced, the resources established here will serve as a foundation for more comprehensive studies of marmoset biology and primate genome evolution.

## Author Contributions

Conceptualization: P.H., A.T., S.J.S.R., A.S., G.W.C., K.H.M., E.E.E., D.F.C., J.L.G., I.A., B.P.

Resource and Data Generation: A.H., S.B., G.H.G, J.G., S.H., S.K.H, B.M., K.M.M, J.P., W.E.S., C.S., I.V., A.T., S.J.S.R., A.S., G.W.C.

Analysis: P.H., T.P., H.L., K.R., M.F.R., F.R., Jo.M., D.Y., L.G.L., A.H., S.K., M.B., H.D.H., M.M., B.M., Ju.M., K.P., S.P., C.R., V.S., C.S., L.W., K.D.M., E.E.E., D.F.C., J.L.G., I.A., B.P.

Initial Draft: P.H., B.P.

Writing/Editing: P.H., T.P., H.L., K.R., M.F.R., F.R., Jo.M., D.Y., L.G.L., A.H., S.K., M.B., H.D.H., M.M., B.M., Ju.M., K.M.M., K.D.M., G.W.C., K.H.M., E.E.E., D.F.C., J.L.G., I.A., B.P.

Figures: P.H., T.P., H.L., K.R., M.F.R., F.R., Jo.M., D.Y., L.G.L., S.K., M.B., G.W.C., E.E.E., D.F.C., J.L.G., I.A., B.P.

Supervision: K.D.M., A.T., S.J.S.R., A.S., G.W.C., K.H.M., E.E.E., D.F.C., J.L.G., I.A., B.P.

Funding: A.T., S.J.S.R., A.S., G.W.C., K.D.M., E.E.E., B.P.

Project coordination: B.P.

All authors read and approved the final manuscript.

## Acknowledgements

We thank Monika Cechova (Masaryk University, Brno, Czechia) for her valuable input and advice on generating the assemblies.

P.H. was supported, in part, by the Jack Baskin and Peggy Downes-Baskin Fellowship. T.P. and J.L.G. were supported, in part, by the Stowers Institute for Medical Research. F.R. was supported by the HSE basic research program. E.E.E. is an investigator of the Howard Hughes Medical Institute. This article is subject to HHMI’s Open Access to Publications policy. HHMI lab heads have previously granted a nonexclusive CC BY 4.0 license to the public and a sublicensable license to HHMI in their research articles. Pursuant to those licenses, the author-accepted manuscript of this article can be made freely available under a CC BY 4.0 license immediately upon publication.

Grants: R01HG014490, U01HG010961, U24HG010262, OT2OD026682, and U24HG011853 (B.P., P.H., and UCSC personnel); R50CA305001 (T.P.); R01AG058851 (S.B., and A.T.); U19AG074866 (S.B., S.H., A.T., S.J.S.R., A.S., and G.W.C.); R01HG002385 (E.E.E.); R01CA266339 (J.L.G.); R35GM151945 (K.D.M.); 5T32HG012344-04 (H.D.H.).

Deep Sequencing of the Omni-C libraries was performed at the UCSF CAT, supported by UCSF PBBR, RRP IMIA, and NIH 1S10OD028511-01 grants.

## Declarations of Interest

E.E.E. is a scientific advisory board (SAB) member of Variant Bio, Inc. All other authors declare no conflicts of interest.

## Resource Availability

### Lead contact

Requests for further information and resources should be directed to and will be fulfilled by the lead contacts, Benedict Paten (bpaten{at}ucsc.edu) or Prajna Hebbar (pnhebbar{at}ucsc.edu).

### Materials Availability

All data used in the assemblies were generated from fibroblasts obtained from marmosets in the University of Pittsburgh colony.

### Data and Code Availability

Sequencing data generated for this project is available on SRA under the bioproject accession *PRJNA1228037*. Assemblies are available from GenBank under the accessions *GCA_049354715.1*, *GCA_049354665.1*, *GCA_049354655.1*, and *GCA_049354675.1*. Any new code written for the analyses and figure generation for this paper is available through https://github.com/ph09/t2t_marmoset/. The temporary hub (until tracks are available directly through the UCSC Genome Browser) is hosted here: https://public.gi.ucsc.edu/~pnhebbar/marmoset/assemblyHub/hub.txt. Any additional information required can be provided by the lead contacts upon request.

### Declaration of generative AI & AI-assisted technologies in the writing process

During the preparation of this work, the authors used LLMs to assist with specific programming tasks and, on rare occasions, to improve the language of the manuscript. After using these services, the authors reviewed and edited the content as needed and take full responsibility for the content of the published article.

