## Supplement for "A Complete Genome for the Common Marmoset"

### Supplementary Figures

##### Fig. S1. PCA from JAX

Principal component analysis (PCA) of marmoset samples colored by cohort/source institution. PC1 and PC2 capture the primary axes of genetic variation, with calJac240 (dark red star) and calJac220 (pink star) marked.

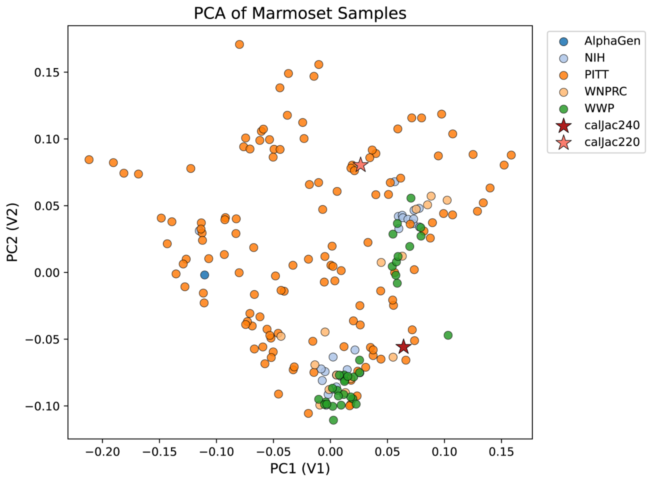


##### Fig. S2. Synteny between HSA and CJA genomes

Genome-wide synteny map comparing human (top) and marmoset (bottom) genomes at a chromosome-level. Colored ribbons connect orthologous regions, illustrating extensive chromosomal rearrangements between the two.

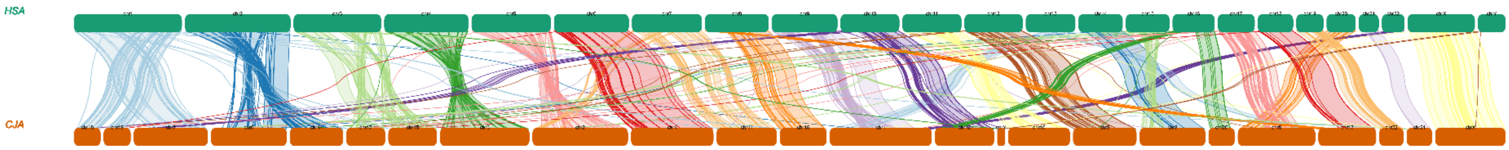


##### Fig. S3. IsoSeq support for Protein Coding Genes

Distribution of protein-coding genes by the number of tissues with Iso-Seq long-read transcriptional support. The majority of genes are supported across all 10 tissues (~9,500), while ~4,500 genes lack support in any tissue.

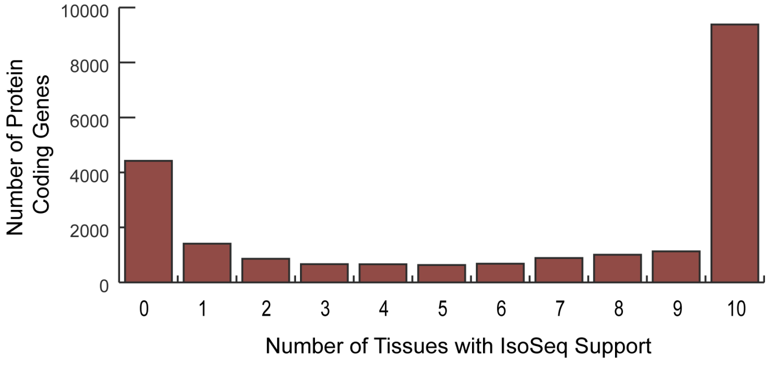

**Fig. S4. Variants in Alzheimer's Disease-Associated Gene Loci**

Predicted variant impacts at the gene level for 75 AD-associated gene loci. Variants with modifier impacts have been excluded for ease of visualization.

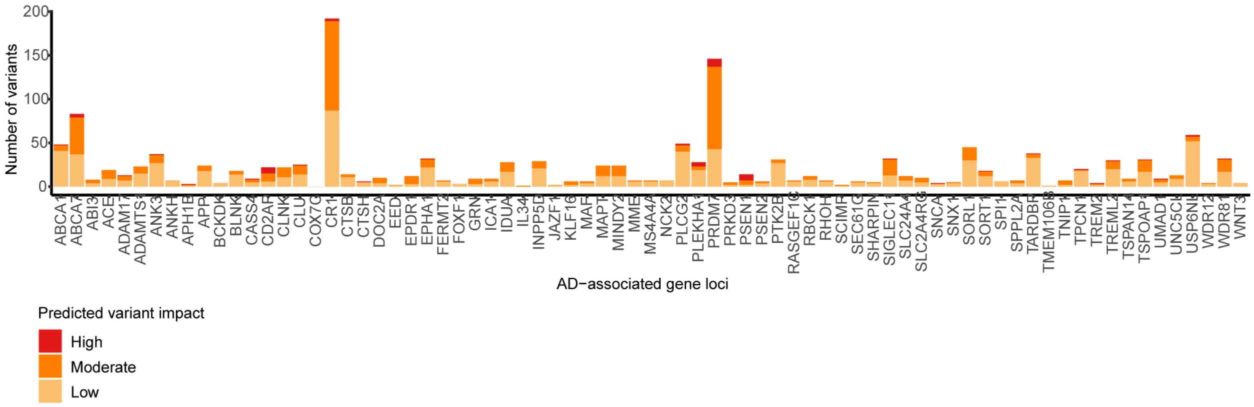


**Fig. S5. Putative marmoset-derived segmental duplication with respect to owl monkey.**

Duplication sequences shared with owl monkey and specific to marmoset are indicated by purple and red, respectively. The part of unaligned sequence shorter than 1000kbp, are indicated by grey

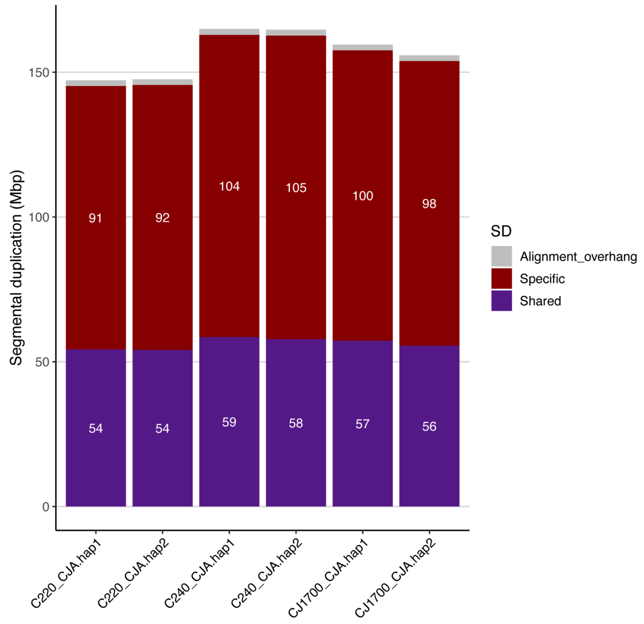


**Fig. S6. FISH-based rDNA copy number estimates per array in the CalJac 240 and CalJac 220 using fluorescent intensity measurements.**

The rDNA FISH signals were measured as fractions of the total fluorescent intensity in the chromosome spread and converted to rDNA copy numbers. Box-and-whisker plots displaying the full distribution of individual measurements. The boxes represent the interquartile range, with the edges indicating the upper and lower quartiles. The line inside the boxes indicates the median. Whiskers show the range from minimum to maximum values. All individual data points are shown.

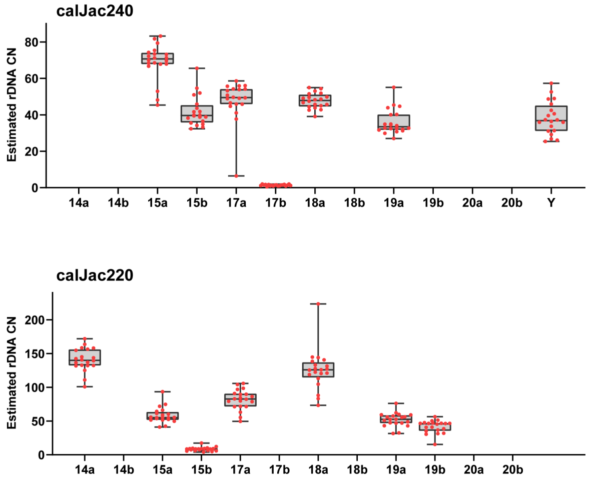


**Fig. S7. rDNA array transmission of selected haplotypes in the calJac 240 family.**
Karyogram panels show acrocentric chromosomes from the father, mother, calJac240, and sister, with rDNA probe signals in red and a chromosome 18-specific marker in yellow. DNA was counterstained with DAPI (blue). Highlighted boxes trace haplotype transmissions: calJac240 inherited a small rDNA array on chromosome 17 from the mother (red boxes), an rDNA-less haplotype on chromosome 19 from the father (blue box), and a Y chromosome rDNA array comparable in size to the father's (white box). Each parent carried one rDNA-less copy of chromosome 18 (yellow boxes), of which calJac240 inherited one; in contrast, the sister inherited both rDNA-less copies, resulting in her chromosome 18 pair entirely lacking rDNA.

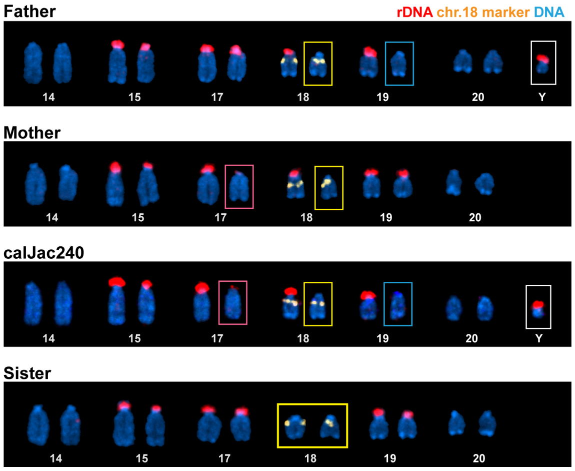


**Fig. S8. Genome Browser Visualization of rDNA Loci on Chr17 and Chr6**
UCSC Genome Browser tracks showing repetitive element annotations at rDNA-containing loci on chromosome 17 (top) and chromosome 6 (bottom) in the calJac240_pri assembly.

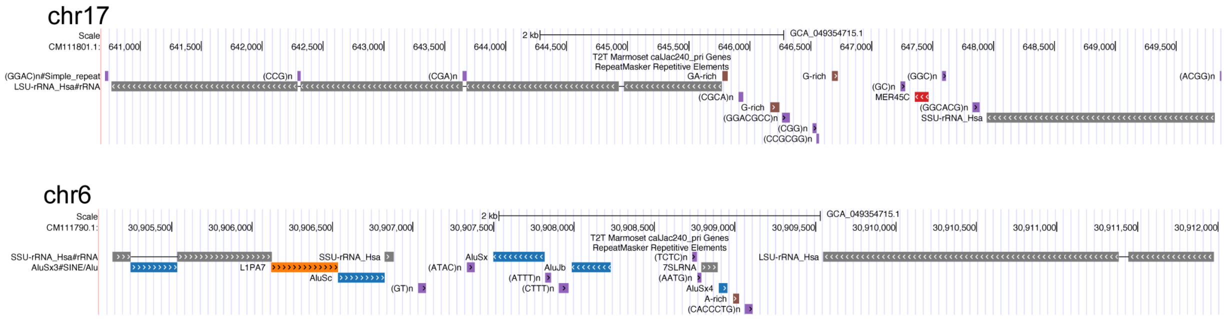


**Fig. S9. Active Alpha Satellite Array Size Across Chromosomes**
Active Alpha Satellite array sizes (in MB) across all chromosomes for the two assemblies (calJac240 and calJac220), shown separately for primary (pri) and alternate (alt) haplotypes.

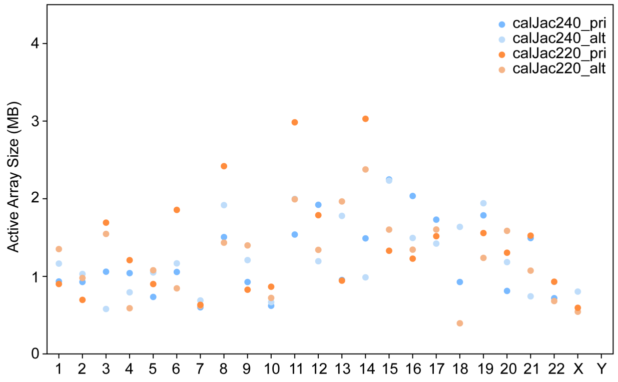

### Methods

#### 1. Sequencing

##### 1.1 Sample Selection & Cell Propagation

We randomly selected two animals (climb IDs 220 and 240) from the University of Pittsburgh colony. Fibroblasts were selected for sequencing to minimize potential chimerism that can occur in hematopoietic tissues due to in utero genetic exchange between siblings.

##### 1.2 PacBio HiFi Sequencing at UW

HMW DNA samples were sheared to a mode length of 20-25 kbp using Megaruptor 3 Hydropores (Diagenode, E07010003) with settings 28/31 and used to generate PacBio HiFi libraries via the SMRTbell Prep Kit 3.0 (PacBio, 102-182-700) using barcoded adapters (PacBio, 102-009-200). At all steps, quantification was performed with Qubit dsDNA HS (Thermo Fisher, Q32854) measured on DS-11 FX (Denovix) and size distribution checked using FEMTO Pulse (Agilent, M5330AA & FP-1002-0275.) Size selection was performed with Pippin HT using a high-pass cutoff of 17 kbp (Sage Science, HTP0001 & HPE7510.) Libraries were sequenced on the Revio platform on 1.5 SMRT Cells 25M each with Revio Chemistry V1 (PacBio, 102-817-900) with Adaptive Loading and 30-hour movies to a coverage target of 60X. Data postprocessing was done onboard and through SMRT Link v13.0.

##### 1.3 ONT Sequencing at UCSC

Frozen fibroblast cell pellets were received from collaborators at the Jackson Laboratory for Genomic Medicine. Ultra High Molecular Weight (UHMW) DNA was extracted from approximately 20 million frozen cells per sample using the NEB HMW DNA Extraction Kit for Tissues (NEB #T3060), following the ONT recommended gDNA Extraction Protocol (v114_revL_27Nov2022). Library preparation was performed using the ONT Ultra-Long DNA Sequencing Kit (SQK-ULK114). The libraries were eluted in 810 µL EB to accommodate four library loads across three flow cells per sample, resulting in a total of 12 libraries per sample. All samples were run on the PromethION 48 sequencer (PRO-SEQ048) with R10.4.1 Flow Cells (FLO-PRO114M). Flow cells were washed using the ONT wash kit (EXP-WSH004) and reloaded with fresh libraries every 24 hours, for a total runtime of 96 hours. The experiment was repeated a second time with additional flow cells to achieve the target of 30X genomic coverage with reads greater than 100 kb in length. Raw Oxford Nanopore signal data were basecalled using Dorado v0.5.3 (Oxford Nanopore Technologies; https://github.com/nanoporetech/dorado) with the super-accuracy model (dna_r10.4.1_e8.2_400bps_sup@v4.3.0) on R10.4.1 flow cells using E8.2 chemistry at 400 bps. Basecalling was performed with simultaneous CpG methylation detection, calling 5-methylcytosine (5mCG) and 5-hydroxymethylcytosine (5hmCG) using the modified base model dna_r10.4.1_e8.2_400bps_sup@v4.3.0_5mCG_5hmCG@v1. Output was written as unaligned BAM (uBAM) files containing both sequence reads and per-read base modification probabilities.

##### 1.4 OmniC Sequencing at UCSC

The Omni-C libraries were prepared using the Dovetail TM Omni-C TM Kit (Dovetail Genomics, Scotts Valley, CA) according to the manufacturer's protocol with slight modifications. Briefly, chromatin was fixed in place in the nucleus. Fixed chromatin was digested *in situ* with DNase I. After digestion, the cells were lysed with sodium dodecyl sulfate (SDS), and chromatin fragments were bound to chromatin capture beads. Chromatin ends were repaired and ligated to a biotinylated bridge adapter followed by proximity ligation of adapter containing ends. After proximity ligation, crosslinks were reversed and the DNA was purified from proteins. Purified DNA was treated to remove biotin that was not internal to ligated fragments. NGS libraries were generated using an NEB Ultra II DNA Library Prep kit (NEB, Ipswich, MA) with an Illumina compatible y-adaptor. Biotin-containing fragments were then captured using streptavidin beads. The post capture product was split into two replicates prior to PCR enrichment to preserve library complexity with each replicate receiving unique dual indices. Sequencing was performed at the Center for Advanced Technologies at the University of California, San Francisco (UCSF CAT) on an Illumina NovaSeq X platform (Illumina, CA) to generate approximately 100 million 2 x 150 bp read pairs per GB genome size.

##### 1.5 Illumina Sequencing at JAX

Whole-genome sequencing (WGS) libraries were prepared by the Genome Technologies group at The Jackson Laboratory using the KAPA HyperPrep kit, according to the manufacturer’s protocols. The protocol entails shearing the DNA using the E220 Focused-ultrasonicator (Covaris), size selection targeting 400 bp, ligating Illumina specific barcoded adapters, and PCR amplification. Libraries were sequenced on an Illumina NovaSeq X Plus, yielding approximately 325 million paired-end reads of 150 bp per individual library. Median genome-wide coverage ranged from 28X to 32X. Percent duplicate reads ranged from 9% to 12%, and base quality scores had a median PHRED score > 35.

##### 1.6 PCA of Sequenced Individuals

To better understand the representativeness of the individuals used to build the assemblies, we compared them to 83 high-coverage, unrelated individuals from captive colonies of the US and Asia. We mapped the short-read sequencing data onto calJac4 using bwa-mem (version 0.6). Duplicate reads were marked using Picard (version 3.2.0). We then called SNPs using GATK (version 4.6.0). HaplotypeCaller was run on each sample independently, then unified using GenotypeGVCFs. To reduce false positives, common quality filters were applied to both indels and SNPs. Using the unified callset of 2 T2T individuals plus 83 MCC animals from captive colonies, we ran PCA using plink (version 1.90b6.20) using only bi-allelic SNPs with minor allele frequency greater than 1%.

#### 2. Assembly

##### 2.1 Assembly

The complete, haplotype-resolved assemblies were generated using a combination of Verkko[1,2] and manual curation. Verkko v2.2.1 was run with the parameters --screen human and -hic1 *R1*fastq.gz --hic2 *R2*fastq.gz. Haplotype-consistent contigs and scaffolds were automatically extracted from the labeled Verkko graph, with unresolved gap sizes estimated directly from the graph structure. Assembly gaps were filled using ONT long reads mapped to the assembly graph with GraphAligner[3]. To address unresolved gaps, we identified reads spanning the flanking nodes and used them to infer traversal paths through the graph. Gap-filling sequences were then incorporated into the assembly graph using Verkko’s `insert_aln_gaps.py` script separately with adjusted parameters. Graph structures that could not be resolved by any of the above approaches were retained as gaps in the final assembly. NCBI Foreign Contamination Screen (FCS) was run on the assemblies to identify any potential contaminant related sequences in the assembly. Completeness was assessed by identifying telomeric sequences at contig ends, counting remaining gaps, and assigning chromosome names to contigs using an alignment to the previous reference. These were the final assemblies that were subjected to polishing and curation.

#### 2.2 Polishing

We aligned 60x-coverage PacBio HiFi DCv1.2 reads for each sample to the diploid assembly using minimap2 with the parameters `-a -x map-hifi --cs --eqx -L -Y -I8g`. In order to correct read phasing in long stretches of homozygosity in these alignments, the PHARAOH pipeline (https://genome.cshlp.org/content/early/2025/05/15/gr.280149.124) was run with default parameters, using as input 30x coverage ONT UL >100kb reads aligned to each haplotype of the input assembly with minimap2 and parameters `-a -x map-ont --cs --eqx -L -Y`. DeepVariant v1.6.1 [4] was run with `modelType=PACBIO` on the PHARAOH-corrected HiFi read alignments. Homozygous-PASS vcf edits were applied to each assembly (29,822 total edits for Maisie, and 25,352 total edits for Baguette.

#### 2.3 Haplotype assignment, chromosome orientation, and numbering

Chromosomes were ordered by length, and names were assigned based on that order (longest as chromosome 1, etc.), except for the sex chromosomes. Orientation was assigned based on the location of the centromere (p-arms at the beginning). For each chromosome pair, the more complete chromosome (telomeres found on both ends, fewer gaps), more accurate (higher QV), and one with a more complete rDNA array was assigned to the primary haplotype. In calJac220, each haplotype has one X chromosome; in calJac240, the primary haplotype contains both sex chromosomes.

#### 2.4 Assessment of the genome assembly

Alignments were generated by mapping the HiFi and ONT reads onto each assembly. Minimap2 (v2.28)[5] was used for the alignment with standard parameters. Flagger (v1.1.0) (https://github.com/mobinasri/flagger) was run using the `flagger_end_to_end_with_mapping.wdl` file deposited in the repository. NucFreq was run using HiFi reads as a part of the assembly_eval pipeline (https://github.com/EichlerLab/assembly_eval). Collapses are defined as locations where the second most common base had a depth of coverage greater than 5, and Duplicated/HiFi-deplete are defined as regions of the assembly with 0 or decreased read coverage by HiFi reads. QV was estimated using Meryl v1.4.1 and Merqury using a hybrid database of 31-mers collected from Illumina and HiFi reads. Hybrid databases were made as in the T2T primates project described in https://github.com/arangrhie/T2T-Polish/tree/master/merqury**.**

#### 2.5 Quantifying improvements in alignment & variant calling

To quantify the improvements in alignment and variant calling with the new T2T assembly, we mapped high-coverage short-read data for 24 families (40 individuals) to both calJac4 and T2T. Illumina paired-end reads from 24 families (40 individuals) were aligned to both the cj1700_1.1 and calJac240_pri reference marmoset genome assemblies using `bwa`[6]. Duplicate reads were marked using `Picard Markduplicates` (Picard Version: 3.2.0). Alignments were filtered with `samtools`[7] to keep only reads with an alignment score (AS) of 50 or higher. Joint-genotyping was performed following GATK best practices for GATK 4.6.0.0 with a genotype quality filter of GQ < 20 and standard hard filters as follows for `SNPs: QD<2.0, FS>60.0, MQ<40.0`,`MQRankSum<-12.5`, `ReadPosRankSum<-8.0, SOR>3.0` and `short INDELs: FS > 200.0, SOR > 10.0,ReadPosRankSum < -20.0`. Repeat regions were filtered using bed files from RepeatMasker. In order to make comparisons between the two genome builds, only variants from homologous regions between cj1700_1.1 and calJac240_pri were analyzed.

Seven PacBio long read libraries were analyzed to create a “gold standard” SV dataset (SRA Project numbers PRJNA1008611, PRJNA566173, PRJDB8242, PRJNA877605, PRJNA1228037). These libraries were a mixture of CLR and CCS/HiFi data. Long reads were aligned to both the T2T assembly and CalJac4 using pbmm2. For CLR data, SVs were called using the PBSV pipeline, and CCS/HiFi data were analyzed with sawfish. Complex SVs and breakends were filtered from the resulting callsets, and the two callsets were merged to produce a reference map of SV genotypes. These SVs were then genotyped across the same 40 MCC animals used in short-read comparisons above, using paragraph.

### 3. Alignment

#### 3.1 Progressive Cactus Alignment

We used Progressive Cactus[8] to construct the multiple genome alignments - a 12-way primary alignment of Human (T2T-CHM13), Bonobo (GCA_029289425.2), Chimpanzee (GCA_028858775.2), Gorilla (GCA_029281585.2), Sumatran Orangutan (GCA_028885655.2), Bornean Orangutan (GCA_028885625.2), Siamang (GCA_028878055.3), Rhesus Macaque (GCA_049350105.1), Crab-eating Macaque (GCA_037993035.1), Common Marmoset (GCA_049354715.1), Dog (GCA_011100685.1), and Mouse (GCA_964188535.1). We used MashTree v1.4.620 with default arguments to compute guide trees for the alignment. The resulting guide tree for the 12-way primary alignment was-

```
(('GCA_011100685.1':0.08093,('GCA_049354715.1':0.0491300000000001,(('GCA_049350105.1':0.00214000000000003,'GCA_037993035.1':0.0025000000000000022):0.02910999999999997,('GCA_028878055.2':0.0196300000000001,(('GCA_028885655.2':0.00178000000000038,'GCA_028885625.2':0.00170000000000007):0.0147399999999975,('GCA_029281585.2':0.0086100000000007,(hs1:0.0065400000000018,('GCA_029289425.2':0.0020000000000000018,'GCA_028858775.2':0.0022999999999999965):0.0042999999999998):0.0015999999999999903):0.00745000000000012):0.0029799999999999827):0.0086400000000009):0.0171199999999996):0.03497):0.05437,'GCA_964188535.1':0.05437);
```

and the Cactus (v2.9.8) commands used to construct the alignments were

```
cactus ./js ./aln.seqfile ./t2t-apes-dog-mouse.hal --batchSystem slurm --caching=false --maxLocalJobs 12000 --consCores 64 --slurmTime 1000:0:0 --configFile ./config-slurm.xml --logFile t2t-apes-dog-mouse.hal.log --doubleMem true --maxMemory 1.4Ti
```

##### 3.2 minigraph-Cactus Pangenome & graph-based SV calls

We constructed a pangenome graph using Minigraph-Cactus[9] from diploid calJac240, calJac220, and CJ1700 genomes (6 haplotypes total) and is referenced on the primary T2T calJac240 assembly. The graphs were constructed all at once, rather than being split by reference chromosome, in order to better account for inter-chromosomal alignments. The graphs were constructed on a Slurm cluster using Cactus v2.8.1 and the following commands.

```
cactus-pangenome ./js-pg ./marmosets_mc.seqfile --outDir
marmosets_mc_pangenome --outName marmosets_mc --reference
calJac240_pri calJac240_alt --noSplit --gbz clip full --gfa clip
full --xg clip full --odgi --vcf --giraffe clip --haplo clip
--vcfReference calJac240_pri calJac240_alt --logFile
marmosets_mc.log --batchSystem slurm --coordinationDir /data/tmp
--batchLogsDir ./batch-logs --consMemory 1500Gi --indexMemory
1500Gi --mgMemory 500Gi --mgCores 72 --mapCores 8 --consCores
128 --indexCores 72 --giraffe clip
```

Minigraph-Cactus produces VCF output alongside the graph representations. We used

`bcftools norm -f`

`vcfbub -l 0 -a 100000`

then

`vcfwave -I 1000`

Then Truvari v4.2.2 to merge similar SVs using

`truvari collapse -r 500 -p 0.95 -P 0.95 -s 50 -S 100000`.

Note that multiallelic sites were split with

`bcftools norm -m -any | bcftools sort`

before Truvari and remerged with

`bcftools norm -m +any | bcftools sort`

after Truvari. Note that these VCFs exclude sites with variants >100 kbp.

#### 4. Gene Annotation & Novel Gene Analysis

Genome annotation was performed using CAT. First, whole-genome alignments were generated using Cactus (12-way primary alignment described above). CAT then used the whole-genome alignments to project the UCSC GENCODEv35 CAT/Liftoff v2 annotation set from T2T-CHM13v2 to the primates. CAT was run with transMap, transMap-pairwise AUGUSTUS, Liftoff, AUGUSTUS-PB, and miniprot modes. transMap lifts over gene annotations from the reference onto all the genomes in the cactus alignment & transMap-pairwise does the same using pairwise minimap2 alignments. Liftoff lifts over gene annotations from a reference onto a minimap2 alignment between the reference transcripts and target genome. The miniprot mode uses protein homology information to improve gene annotations. CAT was given Iso-Seq FLNC data to provide extrinsic hints to the Augustus PB (PacBio) module of CAT, which performs ab initio prediction of coding isoforms. CAT then combined these ab initio prediction sets with the various human gene projection sets to produce the final gene sets and UCSC assembly hubs used in this project.

##### Novel gene annotation and curation of the integrated protein-coding gene annotation set

The annotations generated by CAT were first compared with those generated by the NCBI RefSeq pipeline. To resolve the differences between the annotations generated by the two pipelines, we provide a unique and useful gene annotation resource in the form of a consensus gene annotation between the two pipelines. To generate this, a reliable orthologous gene set was first generated. The orthologous genes were identified using the transMap method of CAT, which uses Cactus alignments to map orthologs. The cases in which genes were mapped to a completely different neighborhood than in humans were flagged and resolved using mappings from liftoff mode. Then, the loci of all protein-coding genes in this set were compared with the orthologous loci assigned by the NCBI RefSeq pipeline. For genes mapped to two completely different loci, transcripts from both were mapped to humans. Depending on the percentage identity of the generated protein to human (>50%), the transcripts were either discarded or assigned as orthologs/novel paralogs. The novel gene loci annotated by either pipeline were collected and filtered at three levels: protein length>200 AA, human protein identity>50%, and Iso-Seq transcript support. These were then merged into the consensus gene set.

#### 5. Repeat Annotation

To identify repeats in the genomes, a RepeatMasker (v4.1.2-p1) run was completed on each genome with a combined library of Dfam 3.6 and Repbase (v20181026) repeat sequences using sensitive settings (-s), the RepeatMasker-compatible NCBI BLAST search engine RMBlast (-e ncbi), and the species tag common marmoset. This was then combined with satellites identified in the cenSat pipeline described in CenSat Annotation.

#### 6. CenSat Annotation

##### Centromeric satellites annotation

Alpha satellites superfamily annotations were merged and sorted into summary bins (activeSF, HOR, mon/hor, diverged HOR dhor, and mixed) based on Superfamily type and size using the script SF_marmoset_summary.sh. Active alphaSatellite arrays were identified.

Ribosomal arrays were annotated using HMMs based on two of Callithrix jacchus rDNA genes; 5.8S (NCBI Reference Sequence XR_008479118.1) and 18S (NCBI Reference Sequence XR_008479122.1). These annotations were then merged to create a complete summary annotation. Scaffolding gaps (sequences of Ns) were annotated with `gap_Annotation.wdl` using Seqtk gap ( https://github.com/lh3/seqtk) to provide complete annotation coverage of rDNA arrays where gaps exist.

Classical satellites (HSATII and HSATII) were annotated using workflow `identify-hSat2and3.wdl,` which runs a Perl script also described in[10], which uses a database of human-specific kmers for annotation. Annotations were then merged to create summary bins over regions where strand switching breaks the annotation. HSat annotations were augmented with simple repeat annotations from RepeatMasker (pentameric repeat ATGGA).

De novo centromeric satellite regions were identified manually using RepeatModeler annotations, dinucleotide content, and AniAnns output[11]. Dinucleotide content was calculated using seqrequester microsatellite (https://github.com/marbl/seqrequester), which was then visualized in the UCSC genome browser to identify regions with dinucleotide content patterns that differ from those of the surrounding

DNA, indicating satellite DNA. Appropriate models from RepeatModeler and RepeatMasker output were then identified and incorporated into the final cenSat annotation dataset. This includes a centromere-associated pentameric repeat n(GGCAA).

All other centromere-associated satellites identified in primates[12] (HSat1A, HSat1B, bSats, gSats, sub-telomeric Pcht, SST1, SATR, TAF11, 5SRNA, CER, and ACRO) were annotated using RepeatMasker[13]. The final CenSat annotation set was created using the workflow `CenSatAnnnotations_Marmoset.wdl,` which compiles and curates satellite annotations described above. This includes resolving overlaps, filtering out small satellite arrays (<2kb), and merging incomplete annotations. Strand information, where available, was recorded in separate files ending with "SatelliteStrandv1.0.bed." Centromere transition (CT) regions were identified by merging satellite annotations within 2 MB and intersecting them with the location of the active alpha satellite array.

##### Alpha Satellite annotation & NWM SuperFamily identification

Annotation of alpha satellite (AS) suprachromosomal families (SFs) and strand inversion analysis were performed using the custom AS SF tracks and AS strand tracks in the UCSC Genome Browser as described[10,14]. SF-tracks for ape/OWM-lineage SFs (APE-SFs) were obtained using the HumAS-HAMMER-SF tool without a custom detection threshold (score/length=0.7) normally used for human and great ape sequences[10,15]. SF tracks for NWM-specific SF analysis were obtained using a new tool specially developed for this paper which was built using the protocols described in previous studies[10,14]. Initially we used our unpublished NWM-AS classification tool trained on Build1.1 of *Callitrix jacchus* assembly which appeared on NCBI server on 19.10.2010 (see Note **S1**) which we validated and re-trained using the current marmoset-T2T assembly. Briefly, the alpha satellite monomers were annotated, extracted, aligned and minimum evolution phylogenetic trees were built in MEGA5[16] or PHYLIP (PHYLIP (Phylogeny Inference Package) version 3.6 (Distributed by the author, Department of Genome Sciences, University of Washington, Seattle, 2005). Each branch was exported and installed as an additional HMM profile in HumAS-HMMER-SF (https://github.com/enigene/HumAS-HMMER) classifier using the detection threshold as described[10]. Next, the marmoset assembly was re-processed by the new tool version, examined, and the profiles renamed and color-coded according to their genome organization. Several cycles of the so-called color separation procedure[10,17] where the uncovered or erroneously covered monomers were picked up, their phylogenetic placement verified using phylogenetic trees, and their HMM installed as a variant monomer of a certain class or subclass into the HMMER tool[18]. After the tool demonstrated a coverage of satisfactory completeness and resolution, it was used to process the assemblies of *Callitrix jacchus* described in this paper. Five UCSC Browser tracks were built for this paper: (1) HumAS-SF track showing the classification of monomers into human-lineage SFs where most NWM-specific monomer classes were identified as SF10 (Ba) or SF11 (Ja) monomers. These tracks were generated with a low custom detection threshold (score/length=0.7) or without a threshold to provide for a complete coverage at the expense of more noise in chromosome arms. (2) NWM-SF track where SF monomer classes starting with SF7 and older were identified the same way as in HumAS-SF, and the younger, NWM-specific classes, were identified as belonging to 6 NWM-specific repeats. Three of them formed small inactive layers located at array periphery: p-monomer (NWM-SF6), 12mer dHOR (NWM-SF5) and 4 generations of S3S4 dimers (NWM-SF 1-4). S3S4 dimer formed the presumed large and centrally located active arrays. (3) The CJ-subSF track which visualizes 6 subSFs within the active NWM-SF1 and shows that there are 6 groups of chromosomes which share within a group particularly close dimer variants in the active arrays. (4) CJ-dimhap track which visualises the chromosome-specific (or smaller group-specific) dimer variants. The NWM-SF annotation tool would work well in any representative of the Cebidae NWM family correctly identifying the dead pericentromeric layers, but could need updating for active layers in some branches. It would work partially to identify S3S4 dimers (without reliable subdivision into SFs 1-4) and would correctly identify SFs 5 and 6 in any NWM species. The subSF and CJ-dimhap tools are either species-specific or genus specific which should be established when the centromeric assemblies of the other representatives of *Callitrix* will become available. (5) The strand track which was the same NWM-SF track, but colored according to the direction of AS in a monomer. Where AS went on the direct strand a monomer was colored blue, and on the reverse strand a monomer was colored red. This track helps to visualize inversions in AS. The details of building the annotation tools, tracks, the results of their examination and evaluation and testing on other available NWM assemblies are provided in Note **S1**.

##### Repeat Length Identification

Satellite arrays were extracted with `bedtools getfasta` and repeat lengths were identified using NTRprism (https://github.com/altemose/NTRprism) with default parameters. NTRprism performs an in-silico restriction digest to identify the most common tandem repeat lengths.

##### Subterminal Satellite Analysis (marmoSAT)

NTRprism analysis of the subterminal regions of the chromosomes indicated the presence of a 170/171 bp monomer unit organized in tandem. BLAST searches revealed that this monomer unit is somewhat conserved across all arrays and was previously described as a subterminal satellite DNA named MarmoSAT[19]. We expanded the characterization of the subterminal satellite blocks of all chromosomes using RepeatMasker to find the matching regions plus 10 kb of flanking sequences on both ends for all arrays. All subterminal arrays were manually curated using visual inspection by generating dot plots with the Dotlet applet with a 15 bp word size and 60% similarity cut-off. We made regressive changes in the consensus sequences used and that enabled us to describe the sequences properly. By manual curation, we were able to identify the beginning and end of 140,316 monomers regarding the consensus generated. All monomeric sequences analyzed were characterized with the same initial and final point and orientation regarding the consensus for the sake of alignment. We aligned all 140,316 subterminal satellite DNA full-length monomeric sequences retrieved from assembled genomes using MAFFT FFT-NS-1. We conducted the phylogenetic analysis by using the Bio Neighbor-Joining method based on the best-fit substitution model (Kimura two-parameter + G, parameter = 5.5047) inferred by Jmodeltest2.

#### 7. Acrocentric Chromosomes

##### 9.1 Designating acrocentric chromosomes

To identify acrocentric chromosomes based on centromeric position, we used the same approach as the T2T Primates project[12], the definitions outlined by the Denver Study Group in 1960[20] and by Levan et al. in 1964[21].

We employed three approaches: a ratio-based approach comparing the lengths of the long (l) and short (s) arms in base pairs (r=l/s), the centromeric index-based approach comparing short arm to the total chromosome length (c) (I=100s*c) and the centromeric index-based approach using the total chromosome length without the centromeric sequence length (c’) (I’=100s*c’). Lengths of the long and short arms exclude the centromeric length. The ratio criteria used was -

$0,7<r<1.58$ for metacentric (M)
$1.58<r<3$ for submetacentric (S)
$r>3$ for acrocentric (A).
Employing this method we categorized chromosomes 3, 4, 6, 7, 8, 9, 11, 14, 15, 16, 17, 18, 19, 20 and Y as either subtelocentric (3, 4, 6, 7, 8, 9, 11, and Y) or acrocentric (14, 15, 16, 17, 18, 19, and 20) in the common marmoset. Of these, chromosomes 14, 15, 17, 18, 19, 20, and Y have gene poor, satellite rich short arms.

Short arms were visualized with moddotplot[22].

#### 9.2 Chromosome spreads and Fluorescent In-Situ Hybridization (FISH)

For the preparation of chromosome spreads, cells were blocked in mitosis by the addition of Karyomax colcemid solution (0.1 µg/ml, Life Technologies) for 6-7h and collected by trypsinization. Cells were incubated in hypotonic 0.4% KCl solution for 12 min and prefixed by addition of methanol:acetic acid (3:1) fixative solution (1% total volume). Pre-fixed cells were spun down and then fixed in Methanol:Acetic acid (3:1). Chromosome spreads were dropped on a glass slide and incubated at 65°C overnight. Before hybridization, slides were treated with 0.1mg/ml RNAse A (Qiagen) in 2xSSC for 45 minutes at 37°C and dehydrated in a 70%, 80%, and 100% ethanol series for 2 minutes each. Slides were denatured in 70% deionized formamide/2X SSC solution pre-heated to 72°C for 1.2 min. Denaturation was stopped by immersing slides in 70%, 80%, and 100% ethanol series chilled to -20°C.

For the 45S rDNA FISH probe, fragments 2927-13332 of the human rDNA reference sequence KY962518.1 corresponding to the conserved coding region of the rRNA gene was cloned into pUC-19 vector and labeled with fluorescence- or biotin-conjugated dUTPs using the nick translation kit (Enzo Life Sciences). Human whole chromosome paints for synteny-based identification of marmoset acrocentric chromosomes were from Applied Spectral Imaging. Labeled rDNA BAC probes RP11-307C12 (GenBank: AL451085.20) and RP11-177M20 (GenBank: AL390122.16) for distinctions of morphologically similar chromosomes 18 and 19 that are both syntenic with human chromosome 1 were from Empire Genomics.

Probes used in combinations were denatured in a hybridization buffer (Empire genomics) by heating to 80°C for 7 minutes before applying to denatured slides. Specimens were hybridized to the probes under a glass coverslip or HybriSlip hybridization cover (GRACE Biolabs) sealed with the rubber cement or Cytobond (SciGene) in a humidified chamber at 37°C for 48-72hours. After hybridization, slides were washed in 50% formamide/2X SSC 3 times for 5 minutes per wash at 45°C, then in 1x SSC solution at 45°C for 5 minutes twice and at room temperature once. For biotin detection, slides were incubated with streptavidin conjugated to Cy5 (Thermo) for 2-3 hours in PBS containing 0.1% Triton X-100 and 5% bovine serum albumin (BSA), and then washed 3 times for 5 minutes with PBS/0.1% Triton X-100. Slides were rinsed in water, air-dried in the dark, and mounted in Vectashield containing DAPI (Vector Laboratories). Z-stack confocal images were acquired on the Nikon TiE microscope equipped with 100x objective NA 1.45, Yokogawa CSU-W1 spinning disk, and Flash 4.0 sCMOS camera (Hamamatsu).

#### 9.3 Estimating rDNA copy number from FISH images

Image processing was performed in FIJI. Acrocentric chromosomes were identified based on size, heterochromatin patterns and chromosome-specific fluorescent markers, and individual rDNA arrays were segmented based on the fluorescent intensity threshold applied to all arrays, adjusted based on the overall brightness of the signal of the spread. The fluorescence intensity of the regions of the same chromosomes that did not contain the rDNA was used to subtract the local background. The background-subtracted integrated intensity was measured for each array, and the fraction of the total fluorescence signal was calculated for each array. The sum of all background-subtracted intensities of all rDNA loci represented the total amount of rDNA per cell. The total rDNA copy number in the genome was estimated from short-read DNA sequencing data (see "Estimating total rDNA copy number from k-mer coverage"). The fraction of the total rDNA fluorescence intensity was used as a proportion of the total rDNA copy number to determine the number of rDNA copies on specific chromosomes in each chromosome spread. Twenty chromosome spreads were quantified for each specimen.

#### 9.4 Estimating rDNA copy number from k-mer coverage

Ribosomal DNA copy numbers were estimated from k-mer frequencies in the Illumina PCR-free short read whole genome sequencing data received from JAX (described above). NCBI entry XR_008479118.1 was used as a reference sequence for marmoset 45S rDNA. The 18S copy number served as a proxy for the greater 45S unit, as each unit contains a single 18S segment. A custom pipeline counted k-mers of size 31 from the 18S consensus in short read Illumina sequencing data and normalized it to counts of 31mers from G/C matched windows elsewhere in the rDNA containing chromosomes. The matched windows were of similar size to the 18S, and ten of these were randomly selected per rDNA-containing chromosome. Any k-mers which also occurred outside the matched windows were removed to ensure that counts were exclusively from the matched windows. k-mer sets were filtered to remove those with whole genome sequencing counts greater than three standard deviations from the mean of the set, or those which were missing entirely. Counts were divided by their genomic multiplicity. Finally, the median count from the 18S k-mers was divided by the median count of the matched windows to yield a copy number approximation. A second normalization step was applied using single copy genes in *Callithrix Jacchus*. Genes were parsed to identify the fifty closest in G/C percentage to the 18S, then twelve of those were manually selected to ensure that they were of reasonable size and from a range of different chromosomes. The copy number for each gene was calculated to a second decimal using CONKORD V8. The average value of all genes was taken, and the true diploid value of 2 was divided by this average. That number was used as a correction factor, so it was multiplied by the copy number approximation from the previous step. A pipeline referred to as CONKORD Version 8 (https://github.com/borcherm/CONKORD) was used for this process.

#### 9.5 Methylation of rDNA arrays

To quantify rDNA methylation per chromosome, ONT reads were aligned back to the assembled diploid genome (minimap v2.26), then filtered to retain the primary alignments overlapping rDNA arrays, not overlapping coordinates deemed unreliable by flagger, with MAPQ60. These reads were binned by chromosome, and then aligned to a reference rDNA unit. Using a BED annotation of the rDNA unit and modkit (v0.3.0), methylation was calculated across the 45S gene or within just the promoter, and copy number was estimated as unit coverage / genome coverage. Only reads >100 kb that are anchored in unique sequence outside the rDNA array and span at least two 45 S units were used.

#### 8. MHC Annotations

Initial identification and annotation of putative classical and nonclassical MHC class I genes and pseudogenes within the two haplotypes of each marmoset telomere-to-telomere (T2T) genomic assembly was performed using nucleotide-level annotations derived from previously published MHC regions based on bacterial artificial chromosome (BAC) clones[23,24]. These published annotations encompass two large segments of the *Callithrix jacchus* MHC class I region: the 854 kb *Caja*-G/F segment and the 1,079 kb *Caja*-B/C segment, which were annotated and reported as separate genomic regions[23,24]. The annotation approach we took was largely similar to that of Yoo et al.[12].

BAC-based annotations were downloaded from NCBI using accession numbers AB600201, AB600202, AB809558, AB809559, and AB809560, following Shiina et al.[23] and Kono et al.[24]. The Caja-E MHC class I locus, which lies between the *Caja*-G/F and *Caja*-B/C regions, was not included in these prior annotations. Therefore, the *Caja*-E gene sequence (NHP09919) was retrieved from the IPD-MHC database[25]. All previously published MHC class I annotations of functional MHC genes included full-length loci comprising both exon and intron sequences as well as corresponding CDS sequences.

Previously annotated *C. jacchus* MHC class I genes and pseudogenes were mapped to chromosome 4 of each T2T haplotype using Minimap2 v2.30 (r1287)[5]. In each alignment, the chromosome 4 sequence was specified as the target, followed by each annotated MHC class I gene or pseudogene as the query. Resulting PAF files were inspected manually, and high-quality target alignments—defined by mapping score, alignment length, and genomic position—were retained for further analysis. All such high-quality targets were considered, as they may reflect copy number variation among closely related genes and/or pseudogenes. Nucleotide sequences corresponding to these high-quality alignments were extracted using custom BioPython scripts[26].

Annotation of putative classical and nonclassical MHC class II genes and pseudogenes in the two haplotypes of each T2T marmoset assembly was carried out using the same general approach. However, because no comprehensive MHC class II annotations are available for *C. jacchus*, MHC II gene and CDS annotations from the human T2T-CHM13v2.0 reference assembly were used as queries. MHC class II regions are generally more conserved among primates than MHC class I regions[12,27], and thus orthologous human annotations were expected to enable reliable identification of MHC class II loci in the marmoset assemblies.

For MHC class I, all newly extracted loci from the assemblies were manually evaluated to determine whether they represented putatively functional genes or pseudogenes. To do so, sequence alignments were constructed for each locus by combining: (i) newly extracted T2T MHC class I sequences, (ii) MHC class I sequences from Shiina et al.[23] and Kono et al.[24], (iii) the IPD-MHC *Caja*-E sequence (NHP09919), and (iv) MHC class I loci annotated from three additional *C. jacchus* genome assemblies (GCA_011100555.2, GCA_009663435.2[28], and one unpublished assembly provided by Dr. Ricardo del Rosario, Broad Institute). The MHC class I regions of these additional assemblies were annotated using the same approach as described for the assemblies (Malukiewicz and Plösch, unpublished data).

An initial multiple sequence alignment containing all previously annotated MHC class I loci and newly identified T2T loci was generated in AliView v2.8.1[29] and aligned using MUSCLE v3.8.31[30]. This alignment was visually inspected to assess sequence similarity and clustering patterns between novel T2T sequences and previously annotated MHC class I loci. Annotations were manually curated to retain a single representative gene annotation per locus for the assemblies. Following assignment of T2T loci to putatively orthologous MHC I locus groups, smaller locus-specific alignments were generated. For putatively functional genes, both genomic and coding sequence (CDS) alignments were curated. CDS sequences for all T2T MHC I annotations were manually extracted by removing introns based on CDS from previously annotated loci and translated into protein alignments within AliView. T2T sequences were examined for premature stop codons or absence of a valid start codon; sequences exhibiting either feature were classified as putative pseudogenes. For MHC class II loci, alignments were generated separately for each putatively orthologous locus due to the greater sequence and structural divergence between alpha- and beta-chain encoding genes. Manual verification and curation of MHC class II loci in the T2T marmoset assemblies then proceeded as described above.

Phylogenetic trees were constructed for all MHC class I loci genes using IQ-TREE with default settings. The best-fitting substitution model was selected automatically, and node support was assessed using ultrafast bootstrap approximation. Resulting trees were manually inspected to verify expected clustering of newly identified T2T loci with known MHC loci. When T2T loci grouped outside their expected clades, alignments were re-examined and loci were reclassified if necessary. Well-supported clades lacking correspondence to previously described MHC loci were interpreted as candidate novel MHC loci.

Finally, manually curated MHC I and II annotations for the marmoset assemblies were compared with results from the automated genome-wide CAT annotation pipeline. Because these approaches rely on different evidence and criteria, minor discrepancies were expected. Supplementary Table MHCGENES provides a detailed comparison of annotations from both methods, along with a consensus classification supported by phylogenetic evidence.

#### 9. Sex Chromosome Analysis

To classify the sex chromosomes into different sequence classes, we used the pipeline as described in Makova et al[31]. Briefly, assemblies were repeat-masked using a combination of RepeatMasker (v4.1.7), WindowMasker (BLAST+ v2.16.0), and Tandem Repeats Finder (TRF v4.09.1), with outputs merged using bedtools (v2.31.1) prior to soft-masking and indexing with samtools (v1.21). Candidate PARs were identified using SEDEF[32] on repeat-masked assemblies, selecting X-Y regions with >99% sequence identity over ≥100 kb. Boundaries were manually refined using lastz [44] dotplots and tandem repeat annotations. Non-PAR regions were classified into satellite, ampliconic, ancestral, and “other” categories. Satellite regions (>0.25 Mb) were identified via RepeatMasker and cenSat annotations. Ampliconic regions combined palindrome calls (PALINDROVER) and intrachromosomal BLASTn (v2.16.0) similarity searches performed on sliding windows across the chromosome sequence (≥50% identity, ≥90 kb), excluding overlaps with PAR and satellite regions. Palindromes were identified on both the X and Y chromosomes from lastz self-alignments (≥98% identity, ≥8 kb, spacer ≤500 kb), filtered against satellite and repeat regions.

#### 10. Segmental duplication

Repetitive elements in the assembled genomes were identified and masked using a combination of three independent approaches. Tandem repeats were annotated with TRF (v4.1.0)[33] using the parameters `trf [asm.fa] 2 7 7 80 10 50 2000 -l 30 -h -ngs`. Interspersed repeats were detected with RepeatMasker (v4.1.6)[34] executed as `RepeatMasker -s -e ncbi -xsmall -species Primates [asm.fa]`. In addition, WindowMasker (v2.2.22)[35] was applied in a two-step procedure consisting of count generation (`windowmasker -mk_counts -mem 16384 -smem 2048 -infmt fasta -sformat obinary -in [asm.fa] -out [asm.count]`) followed by interval masking (`windowmasker -infmt fasta -ustat [asm.count] -dust T -outfmt interval -in [asm.fa] -out [asm.interval]`). The outputs from all three methods were merged to generate a unified set of repeat annotations, which was used to soft-mask the genomic sequences. Segmental duplications were subsequently detected on the repeat-softmasked assemblies using SEDEF (v1.1)[32]. Resulting duplication calls were filtered to retain alignments longer than 1 kb with greater than 90% sequence identity and with less than 70% satellite sequence content.

Mapping of segmental duplications (SDs) across assemblies was performed through a multi-step procedure. First, individual SDs separated by less than 100 kb were merged into continuous SD chains. Second, only SD chains located within reliably mappable regions were retained, defined as those overlapping alignment blocks of at least 100 kb. Third, these chained SDs were projected onto putative homologous SD loci that contained a minimum of one 100 kb unique flanking sequence. To further evaluate positional homologs for sequence novelty, pairwise alignments of SD sequences were generated using minimap2 (v2.26)[5]. SDs were classified as candidates for assembly-specific duplications under the following conditions: (i) absence of a positional homolog among query SDs annotated in the target genome, (ii) substantial divergence in sequence content, with less than 80% of the sequence conserved, and (iii) evidence of copy expansion, defined as a minimum twofold increase in size. We also classified more lenient variation in SDs length at least 20% different, as polymorphic by length. Comparison of duplicated sequences with owl monkey was performed by directly aligning SDs allowing for secondary alignment using the following parameters: `minimap2 -cx asm20 --eqx -Y -N 1000 -p 0.1 [asm.fa]`.

#### 11. Methylation

Reads from calJac220 and calJac240 were pooled within each sample and aligned to their corresponding diploid genome assemblies (primary and alternate haplotypes) using minimap2 (v2.28). Alignments to the primary and alternate assemblies were performed separately. Alignments were then converted to BAM format and indexed using samtools (v1.21). Aligned BAM files had 5mC methylation aggregated across reference CpGs using modkit (v0.4.2), to produce a bedMethyl file. BedMethyl files were trimmed with awk to make four column bedgraphs of the fraction modified. Bedgraphs were converted to bigwigs using bedgraphtobigwig (v2.10), for browser use.

### Supplementary Note 1

#### Alpha satellite (AS) analysis in the T2T genome of the common marmoset (New World monkey).

##### Old view of the NWM centromeres

In a nutshell, the old view of OWM and NWM centromeres was that their organization was pan-chromosomal (i.e., that all centromeres had more or less the same AS repeats) and that there were no HORs[36]. Later, however, some reports demonstrating HORs in NWM were published[37], but it was not entirely clear whether these were in active or dead centromeres.

##### Old annotation tool

*Callithrix jacchus* genome build 1.1 appeared on the NCBI server on 19.10.2010 (release date: 25 October 2010); it had reference assemblies for chromosomes 1 to 22, X and Y, and unplaced (Unk). This assembly was used to construct the initial version of the NWM AS classification. AS monomers were identified using the PERCON tool[15] and S3 and S4 consensus sequences[36]. It turned out that most AS was in the unplaced contigs (Unk chromosome). All the monomers with a length between 160 and 180 bp and rs>=0.35 were extracted and aligned, and the monomer phylogenetic tree was constructed. It had 18 branches, which were extracted, converted to PERCON annotation standards, and used to annotate the *C. jacchus* and other NWM reads for internal lab purposes.

##### Validation/update of the old tool, building NWM-HMMER-SF

The annotation standards of the old tool were converted to HMMs and used with the HMMER platform. This tool was used to annotate marmoset assemblies and was subsequently validated and updated using these annotations. The methods used were identical to those we reported in a similar project to validate and update the old annotation tool for the Old World Monkeys[14,36]. Random selections of monomers and all the monomers from one or two chromosomes were extracted from the initial annotation and viewed on the trees. It was found that the old tool annotated the assemblies adequately; some finer branching within the S3 and S4 classes was noted and incorporated into the annotation, leading to the discovery of 6 subSFs within SF1.

##### Annotation of the T2T marmoset assemblies

Two T2T genome assemblies of Marmoset (calJac220 and calJac240) were analyzed using NWM-SF HMMER-based classifier.

**Table SN1. Consensus order, size, and divergence of NWM-specific AS layers in marmoset** (in one complement with both X and Y chromosomes).

| Layer | Monomer classes | CenSat | Total length, kbp | Color in graphs and tracks | Similarity |
|---|---|---|---|---|---|
| NWM-SF1-1 | S3-1 & S4-1 | active_dimer (S3S4) | 20591 | dark brown | 0.953 |
| NWM-SF1-2 | S3-2 & S4-2 | | | brown | 0.951 |

| | | | | | |
|---|---|---|---|---|---|
| **NWM-SF1-3** | S3-3 & S4-3 | | | dark red | 0.966 |
| **NWM-SF1-4** | S3-4 & S4-4 | | | red | 0.938 |
| **NWM-SF1-5** | S3-5 & S4-5 | | | light red | 0.909 |
| **NWM-SF1-6** | S3-6 & S4-6 | | | light pink | 0.951 |
| **NWM-SF2** | S3b & S4b | dimer (S3bS4b) | 417 | green | 0.796 |
| **NWM-SF3** | S3c & S4c | dimer (S3cS4c) | 243 | blue | 0.786 |
| **NWM-SF4** | S3d & S4d | dimer (S3dS4d) | 34 | black | 0.764 |
| **NWM-SF5** | p | mon (p) | 71 | cyan | 0.819 |
| **NWM-SF6** | abcdefghijk | dhor (11mer) | 67 | orange | 0.691 |

Note: The AS SF layers are numbered according to the consensus order, from the active array (SF1) towards the arms on each side. The relative order of the last 2 layers could not be established, as they occur together in only one instance (on chr16) and are located some distance from the centromere.

#### Note on SF names

Six NWM-specific AS suprachromosomal families (NWM-SFs) were identified in the final annotation. There are also the OWM-SFs (OWM-specific families) identified recently[14] and the SFs of the ape lineage known for a long time and traditionally used w/o any prefix. Of these, SFs7 and older are shared with OWM, and SFs11 and older are shared with NWM. In the context of one primate branch, people often use SF numbers w/o a prefix (as we did in the main text). So, we propose, from now on, using the APE-prefix for the ape-lineage SFs, if needed, to avoid confusion.

#### Note on similarity calculation

Up to 1000 random dimers per layer were extracted. Dimers on the + strand were taken as S3S4; dimers from the - strand were taken as S4S3, reversed-complemented, and renamed to S3S4. Dimers were aligned, and mean divergence was calculated.

We estimated divergence from the multiple sequence alignment using a column-based, sitewise approach with pairwise deletion: in each column, we ignored gaps ("-") and ambiguous bases ("N"), considered only A/C/G/T, and computed the fraction of mismatched pairs among the remaining symbols. We then averaged the column values to obtain the overall divergence. Sequence similarity was defined as 1 minus this divergence.

**Fig. SN1. Annotation of AS layers in marmoset centromere 1 by the NWM-SF tool.** One can see every AS monomer with its SF classification in the full view of the annotation track and appreciate dimeric and 11mer periodicities. Note that if one inverts the blue pieces in the strand track (then all AS would go on reverse strand), the consensus order of layers S3S4_[S3S4]b_[S3S4]c_[S3S4]d_11mer dHOR will be restored. Also note that the gaps between the arrays are mostly composed of SDs, which were probably inserted in the originally continuous centromere and disrupted the AS array.

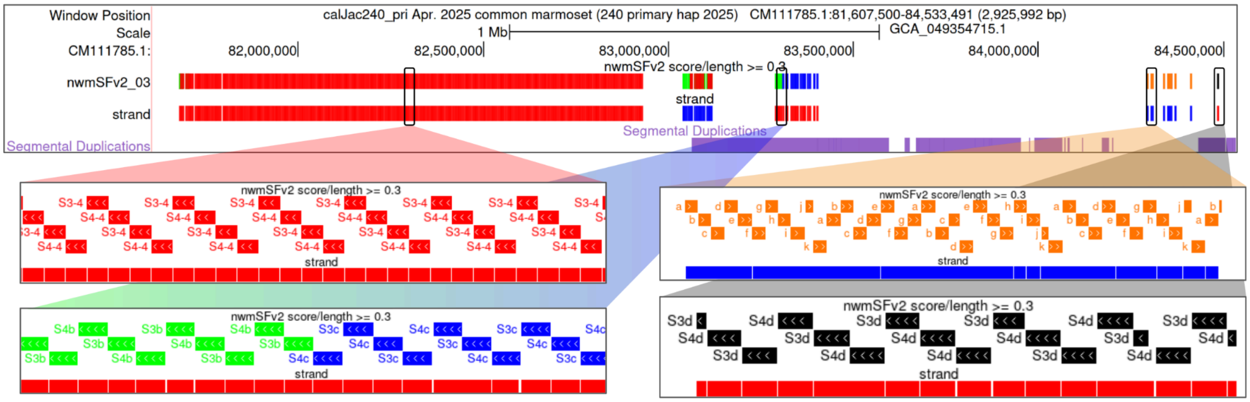


The large active arrays were formed by S3S4 dimers, followed by small inactive flanking S3bS4b (NWM-SF2) in most chromosomes and S3cS4c (NWM-SF3) in many, in that order. In just a few locations (see Table S2), small arrays of S3dS4d(NWM-SF4) were present, usually somewhat removed from the centromere. Genomic size and intra-array similarity decrease progressively from the active centromere outward, as predicted by the expanding centromere/layered expansion model[10,38]. It indicates that the inactive generations of S3S4 dimers (b-c) likely represent the remnants of the centromeres of the marmoset ancestors within the NWM branch (see schematics in Fig. 6A). Localization of SF4 clusters in non-centromeric locations may indicate that the centromere has re-positioned after the split of NWM from the ape/OWM lineage. Note that such small satellite fragments may not necessarily be relics of the proper centromeric arrays in such distant ancestors, but may instead come from the AS-containing SDs. The latter, however, are usually also concentrated around centromeres (except for the acrocentric short arms), so the distant locations may still indicate centromere re-positioning.

#### Interpretation of the p-layer

The consensus order of the 2 smallest layers (NWM-SF 5 and 6) could not be established because they occur only once in adjacent locations. All but one SF6 array is located in the short arms of the rDNA bearing acrocentric chromosomes (Table **SN2**). These arms are very similar to each other, and the arrays' positions and structures are similar across arms, not adjacent to the centromere or any other AS array. So, in effect, it is the same array that may be considered an AS-containing SD together with the surrounding non-AS sequences (Fig. **SN2**). The remaining location on chr16, which is also acrocentric, differs in structure and context and contains the p-mon array adjacent to the 11mer HOR array, which could be taken as weak support for the initially centromeric location of the p-arrays. One could suggest that p-mons went either before or after the 11mer HOR in the evolutionary succession of AS dead layers. However, on a phylogenetic tree of consensus monomers, the p-mon is close to one of the monomers of the 11mer (monomer a), not equidistant to them as would be expected if it were ancestral to the 11mer. Therefore, the likely interpretation of the p-mon layer within the framework of an expanding centromere model would be as an expansion of one of the HOR monomers, which got amplified and formed the next generation of centromeres in a hypothetical NWM ancestor. However, the intra-array similarity in the p-arrays (similar within each one and across all of them) was ~82%, which is much higher than in 11mer and even in all much younger inactive dimer arrays. So, this is the case in which the phylogenetic age (position on a monomer tree) and amplification age (intra-array divergence) of a repeat differ dramatically (see discussion in the Suppl. Note in Makova et al 2024[31]). This calls for a more complex explanation.

Perhaps the p-arrays were indeed generated as an expansion of one of the HOR monomers, but this expansion occurred at a late date when the HOR centromeres were already dead and remained only as small bits and perhaps far from the centromeres. Such expansions in long dead centromeric material are well documented in humans, especially in the long arms of acrocentric chromosomes. They are revealed as low-copy HOR expansions of APE-SFs with 4-6 repeats, specific to 2 or several acrocentric chromosomes. Such expansions usually occur within some of the copies of AS-containing SDs, numerous in the short arms[10,15]. Thus, the p-layer may never have formed the active regions in any NWM ancestor and should not be considered an independent layer. We have marked that by a special note in the legend of Fig. 6a.

To gain further insight into the origins and fate of the p-arrays, we have extracted them and compared them. Genomic organisation of the arrays is shown in Fig. **SN2.** One can appreciate a similar organisation of the loci in all acrocentrics except for chr16. Detailed comparison indicated precisely matched pairs: cens 17 and 19 (type 1, a complete locus with a large L1 element between p-subarrays); cens 15 and Y (type 2, a near-complete locus with shortened right subarray); cens 14 and 18 (type 3, a deleted locus with only the right subarray). Cen20 array is very close to type 3 with minor differences, and cen16 locus is completely different, although a similar adjacent unidentified composite repeat (made of various Alus and U6 in a regular manner) can be seen on the RepMask track. As these relationships did not parallel those between active dimers in the centromeres, we constructed a phylogenetic tree of the p-arrays, which confirmed the above-stated pattern (Fig. **SN3**).

The rDNA-centromere coordination hypothesis we propose in this paper (see the main text Discussion) suggests that similar or even near-identical repeats in the centromeres of acrocentric chromosomes may lead to occasional exchanges of entire short arms between non-homologous chromosomes (with recombination breakpoints somewhere in the active arrays). One could speculate that the frequency of such exchanges would be proportional to the degree of similarity between centromeres. In such a case, it is expected that phylogenetic relationships of various sequences along the length of the short arms will be the same as in centromeres (or at least in the p-arm parts of the centromeric arrays). Our analysis of the p-array phylogeny indicates that this is not the case. As the p-array is likely only a small part of a larger PHR (pseudo-homologous region), these regions may also serve as recombination points, leading to the exchange of only parts of the short arms (which still contain rDNA loci) and partially obscuring the expected relationships. Therefore, it would seem that the general architecture of acrocentric short arms (repeat-rich and gene-poor) has evolved to provide multiple possibilities for non-homologous exchange that involves the rDNA cluster.

**Fig. SN2. The p-loci in acrocentric chromosomes.**

**cenY (808) reversed**

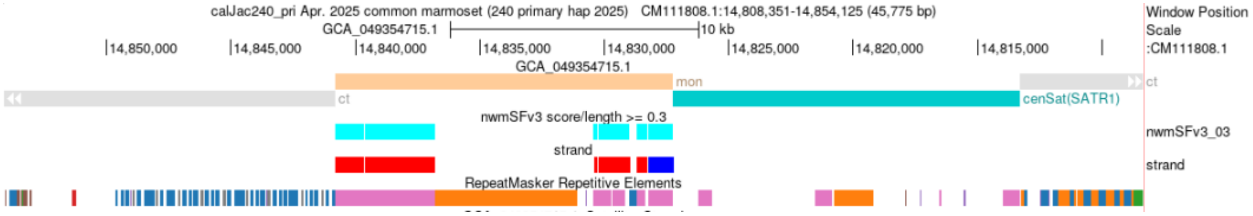


**Cen15**

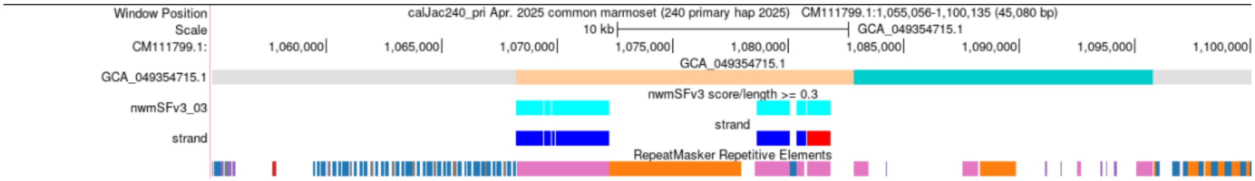


**Cen17**

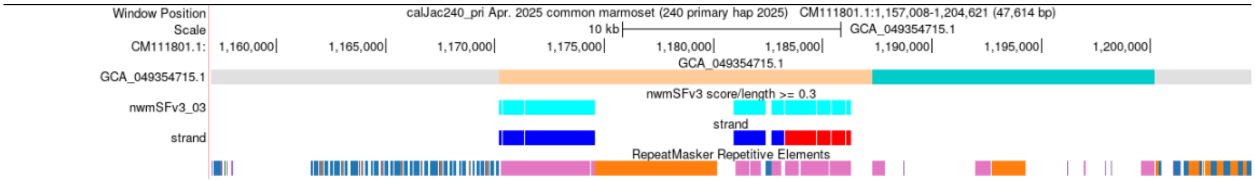


**Cen19**

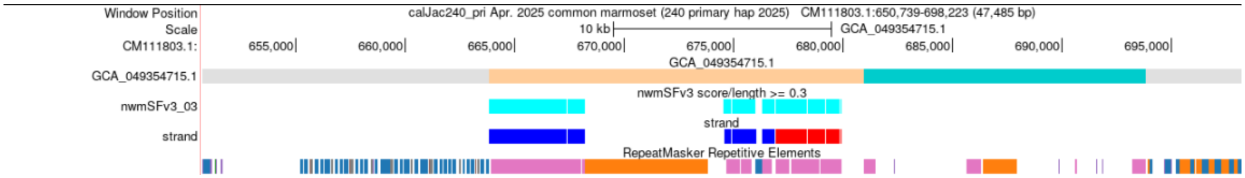


**Cen14**

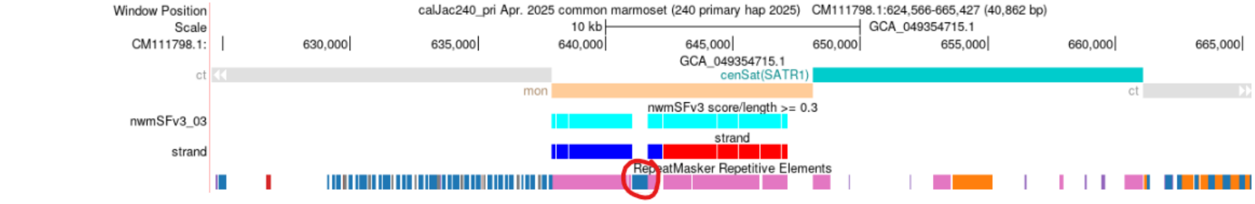


**Cen18**

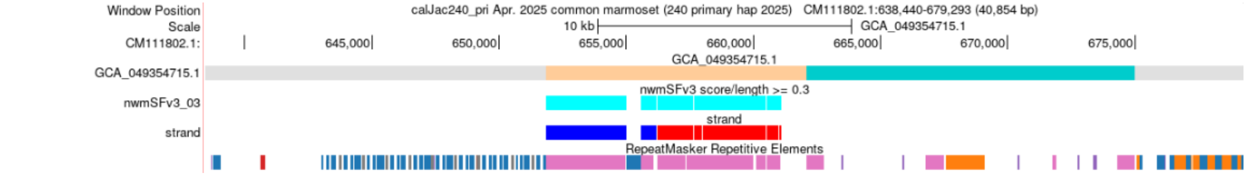


**Cen20**

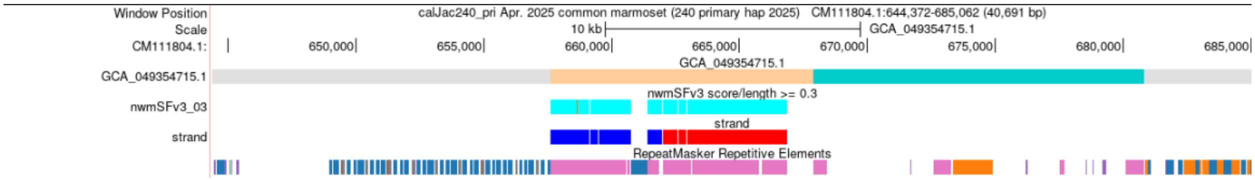


**Cen16**

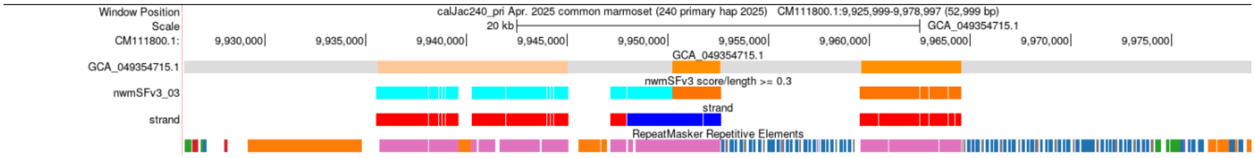


**Fig. SN3. Phylogenetic relationships of the p-arrays in the acrocentric chromosomes do not exactly parallel dimhap relationships in the centromeres.** A. Minimum evolution phylogenetic tree of the p-arrays. The 15/Y and 17/19 pairs are exactly identical over the indicated span (over 6 kb) and the 14/18 pair has one mismatch. B. Minimum evolution phylogenetic tree of chromosome-specific SF1-1 dimhaps. One can see that near-identical p-arrays of 14/18 pair show up in 2 chromosomes which have significantly different (6 differences between ~340 bp dimer consensus sequences) centromeric repeats. This may be indicative of a recent recombination which happened somewhere in the short arm PHRs. Note that according to independent analysis in Fig. 5F, chrs 14 and 18 share a robust PHR (>99% identity) region.

A.

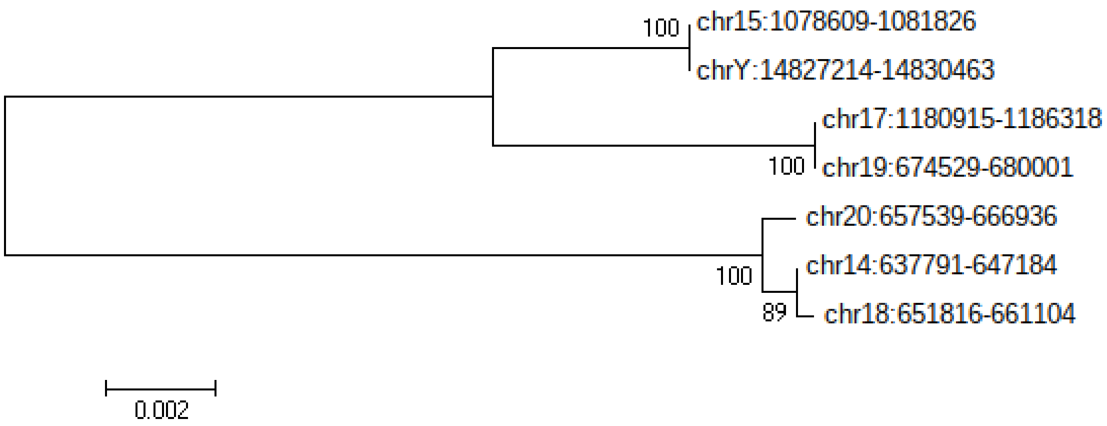


B.

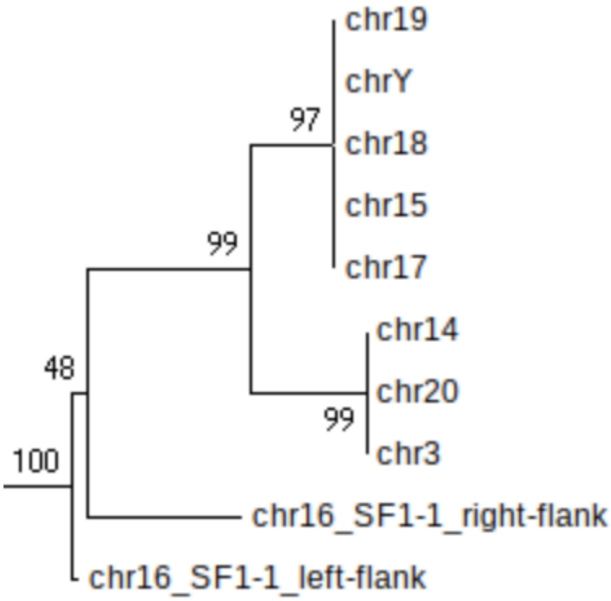


#### Test for HOR presence in the active regions

To test whether any HORs are present in the marmoset active arrays, we used NTRprism, which identifies periodically spaced kmers in repeat arrays and thus detects various periodicities[10]. The results are shown in Fig SN4. We confirmed the general HOR absence by running NTRprism on all AS in the primary haplotype, which identified a dimeric repeat with some tetrameric repeats. The only exception we identified was on the X chromosome (CM111807.1), where we found evidence of higher-order repeats, the most common of which is a 28-mer.

**Fig. SN4. NTRPrism check for HORs**

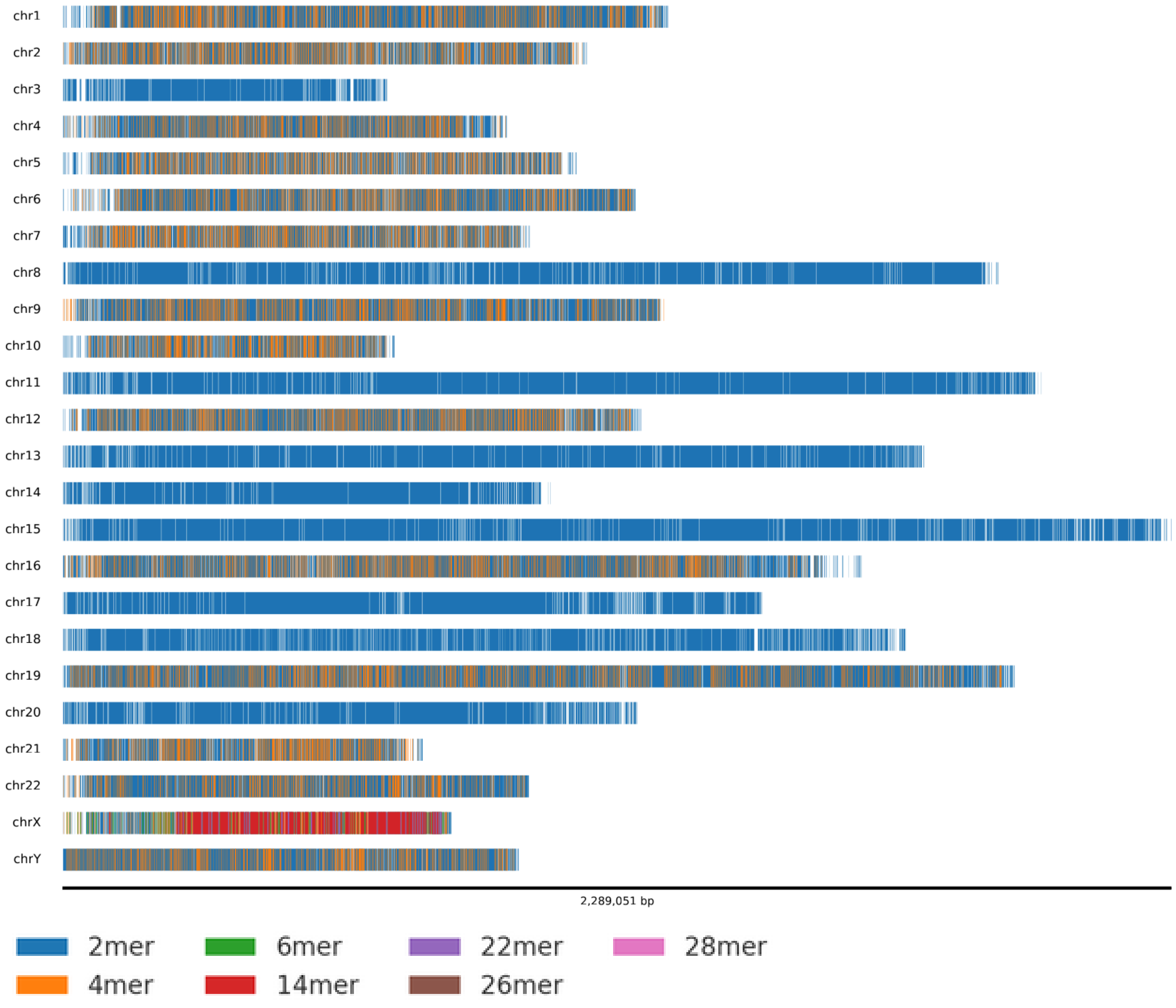


#### SubSFs in active SF1

It was believed in the old literature that centromeres in both the OWM and NWM were made by very similar sequences (pan-chromosomal organisation)[36]. We have recently shown that this was not exactly the case in macaques (OWM)[14], where a limited degree of chromosomal specificity was documented. To test this in NWM, we have extracted the S3 and S4 monomers separately from active arrays of all marmoset chromosomes, aligned and clustered them according to the consensus differences they share (Fig **SN5**). This revealed 6 distinct types, each located within a specific chromosomal group, as summarized in Tables **SN1** and **SN2**. The same groups were revealed by processing S3 and S4 monomers. Thus, we have identified 6 subSFs within SF1, each characterized by a specific dimer sequence and located on specific chromosomes (Table **SN2**).

**Fig. SN5. Alignment plot of S3 monomer subtypes.** Shade helps to distinguish subtypes. 300 S3 monomers were randomly picked from all marmoset chromosomes. Each type was converted to HMM and used in the subSF annotation tool.

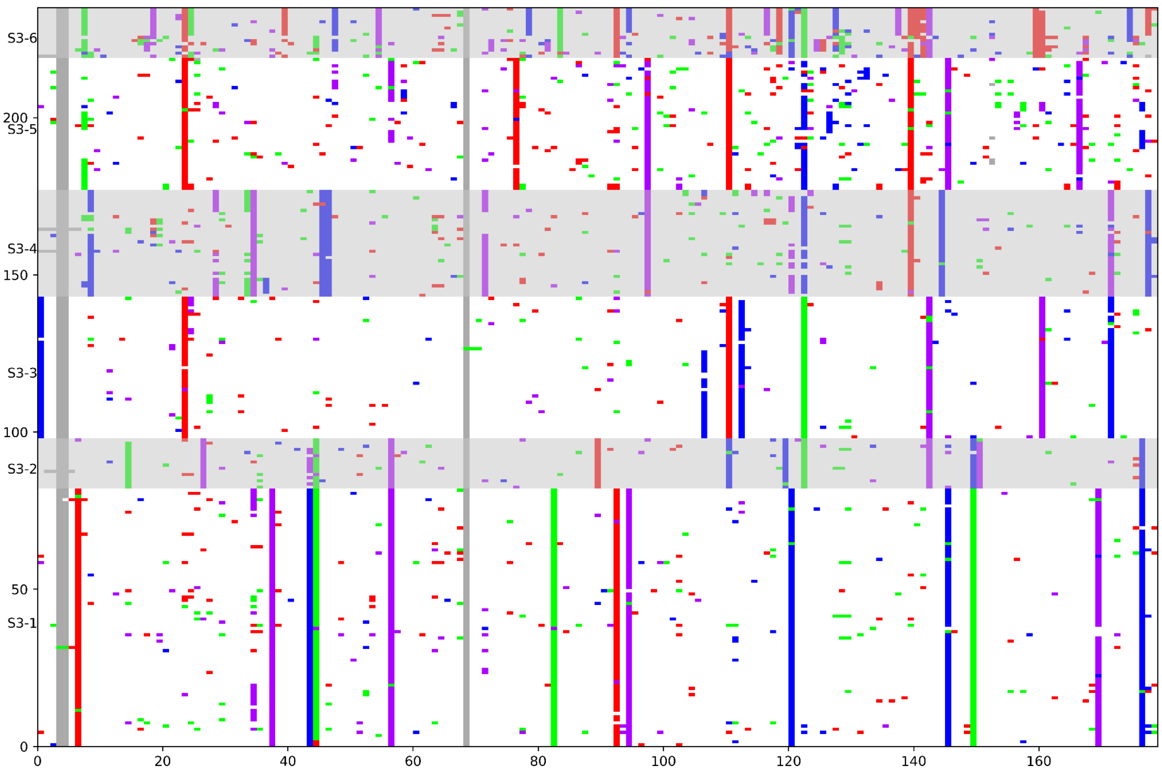


Each monomer type was converted to HMM, and 12 new monomer types were installed into the NWM-SF annotation tool instead of the 2 HMMs for S3 and S4 monomers, thus creating the CJ-subSF annotation tool. It was used to annotate the assembly. The results are shown in Table **SN2** and Fig. **6C** (left) in the main text. One can see that each centromere contains mostly the dimers of only one kind; minor amounts of other dimers are present in most. Manual inspection of annotation tracks revealed that, in all cases, the large central domain (active array) was formed by the same subSF, but could often be flanked by short arrays of another subSF and, further distally, by even smaller arrays formed by mixtures of monomers belonging to different subSFs. This indicated (as previously observed in apes) that the centromeres of one subSF could be inactivated and replaced by the insertion and expansion of another subSF, which moved the halves of the old centromere sideways, followed by subsequent shrinking of the flanking old arrays. Such a sequence of events is stipulated by an expanding centromere/layered expansion model[10,38]. In particular, cases in which several active SFs or subSFs in the same genome compete and replace one another were called centromeric interlayers[31]. The subSF-homogeneous large active arrays identified by our CJ-subSF annotation tool always contained CDRs, as observed in the methylation Browser track. CDRs are the dips in overmethylated active arrays, which correspond to kinetochore position[10].

**Table SN2. The statistics of the marmoset AS monomers annotated by the final version of the NWM_HMMER-SF/subSF tool.**

| A | B | C | D | E | F | G | H | I | J | K | L | M | N | O | P | Q | R | S | T | U | V | W | X | Y | Z | AA | AB | AC | AD | AE | AF | AG | AH | AI | AJ | AK |
|---|---|---|---|---|---|---|---|---|---|---|---|---|---|---|---|---|---|---|---|---|---|---|---|---|---|---|---|---|---|---|---|---|---|---|---|---|
| contig | chrom | S3-1 | S4-1 | S3-2 | S4-2 | S3-1 | S4-1 | S3-4 | S4-4 | S3-5 | S4-5 | S3-6 | S4-6 | S3b | S4b | S3c | S4c | S3d | S4d | p | a | b | c | d | e | f | g | h | i | j | k | Aa | Ja | Pa | Ta | Qa |
| CM111799.1 | 15 | 6512 | 6513 | 12 | 0 | 1 | 8 | 4 | 3 | 93 | 291 | 220 | 0 | 92 | 112 | 82 | 83 | 0 | 0 | 41 | 0 | 0 | 0 | 0 | 0 | 0 | 0 | 0 | 0 | 0 | 0 | 0 | 2 | 0 | 0 | 1 |
| CM111803.1 | 19 | 5657 | 5657 | 9 | 0 | 0 | 5 | 0 | 0 | 55 | 202 | 165 | 0 | 88 | 106 | 86 | 87 | 2 | 2 | 56 | 0 | 0 | 0 | 0 | 0 | 0 | 0 | 0 | 0 | 0 | 0 | 0 | 2 | 0 | 0 | 0 |
| CM111802.1 | 18 | 4794 | 4786 | 13 | 0 | 0 | 8 | 17 | 11 | 218 | 412 | 220 | 1 | 91 | 109 | 86 | 87 | 0 | 1 | 51 | 0 | 0 | 0 | 0 | 0 | 0 | 0 | 0 | 0 | 0 | 0 | 0 | 0 | 0 | 0 | 0 |
| CM111801.1 | 17 | 4149 | 4152 | 9 | 0 | 0 | 5 | 0 | 0 | 47 | 192 | 162 | 0 | 84 | 98 | 86 | 87 | 1 | 2 | 58 | 0 | 0 | 0 | 0 | 0 | 0 | 0 | 0 | 0 | 0 | 0 | 0 | 0 | 0 | 0 | 0 |
| CM111804.1 | 20 | 3443 | 3416 | 2 | 0 | 5 | 7 | 0 | 0 | 10 | 83 | 71 | 0 | 95 | 105 | 77 | 80 | 0 | 0 | 51 | 0 | 0 | 0 | 0 | 0 | 0 | 0 | 0 | 0 | 0 | 0 | 0 | 0 | 0 | 0 | 0 |
| CM111798.1 | 14 | 2891 | 2883 | 7 | 0 | 40 | 11 | 2 | 2 | 61 | 273 | 176 | 1 | 95 | 127 | 210 | 212 | 0 | 0 | 51 | 0 | 0 | 0 | 0 | 0 | 0 | 0 | 0 | 0 | 0 | 0 | 0 | 0 | 0 | 0 | 0 |
| CM111808.1 | Y | 2730 | 2732 | 6 | 0 | 0 | 4 | 0 | 0 | 15 | 179 | 172 | 0 | 71 | 80 | 86 | 87 | 4 | 6 | 42 | 0 | 0 | 0 | 0 | 0 | 0 | 0 | 0 | 0 | 0 | 0 | 0 | 0 | 0 | 0 | 0 |
| CM111787.1 | 3 | 1635 | 1526 | 17 | 0 | 59 | 9 | 0 | 0 | 147 | 436 | 158 | 0 | 92 | 156 | 1 | 1 | 1 | 1 | 0 | 0 | 0 | 0 | 0 | 1 | 0 | 0 | 0 | 0 | 0 | 0 | 39 | 14 | 0 | 0 | 0 |
| CM111793.1 | 9 | 12 | 12 | 3528 | 3527 | 6 | 3 | 0 | 0 | 2 | 48 | 52 | 10 | 22 | 31 | 20 | 18 | 0 | 0 | 0 | 0 | 0 | 0 | 0 | 0 | 0 | 0 | 0 | 0 | 0 | 0 | 0 | 1 | 7 | 6 | 0 |
| CM111788.1 | 4 | 172 | 19 | 2265 | 2252 | 62 | 12 | 0 | 0 | 4 | 298 | 93 | 0 | 46 | 77 | 63 | 57 | 13 | 11 | 0 | 0 | 0 | 0 | 0 | 0 | 0 | 0 | 0 | 0 | 0 | 0 | 0 | 0 | 0 | 0 | 1 |
| CM111795.1 | 11 | 0 | 3 | 55 | 42 | 5816 | 5813 | 0 | 0 | 5 | 63 | 50 | 1 | 121 | 143 | 31 | 31 | 0 | 0 | 0 | 0 | 0 | 0 | 0 | 0 | 0 | 0 | 0 | 0 | 0 | 0 | 0 | 0 | 0 | 0 | 0 |
| CM111792.1 | 8 | 0 | 1 | 3 | 0 | 5549 | 5548 | 0 | 0 | 3 | 34 | 23 | 0 | 311 | 329 | 0 | 2 | 2 | 2 | 0 | 0 | 0 | 0 | 0 | 0 | 0 | 0 | 0 | 0 | 0 | 0 | 0 | 1 | 0 | 0 | 0 |
| CM111797.1 | 13 | 207 | 3 | 5 | 0 | 4983 | 4442 | 0 | 0 | 8 | 758 | 16 | 0 | 59 | 70 | 75 | 77 | 0 | 1 | 0 | 0 | 0 | 0 | 0 | 0 | 0 | 0 | 0 | 0 | 0 | 0 | 0 | 0 | 0 | 0 | 0 |
| CM111800.1 | 16 | 168 | 155 | 4 | 0 | 6 | 7 | 4361 | 4360 | 3 | 131 | 96 | 0 | 77 | 93 | 0 | 0 | 0 | 0 | 69 | 7 | 7 | 7 | 6 | 3 | 2 | 2 | 2 | 3 | 2 | 2 | 0 | 0 | 0 | 0 | 0 |
| CM111785.1 | 1 | 38 | 13 | 11 | 0 | 46 | 12 | 3338 | 3339 | 99 | 285 | 125 | 1 | 110 | 142 | 163 | 164 | 9 | 9 | 0 | 16 | 14 | 15 | 15 | 15 | 15 | 15 | 15 | 14 | 13 | 12 | 0 | 1 | 0 | 0 | 0 |
| CM111786.1 | 2 | 0 | 3 | 12 | 0 | 11 | 11 | 3008 | 3009 | 42 | 191 | 135 | 0 | 28 | 83 | 1 | 1 | 0 | 0 | 0 | 0 | 0 | 0 | 0 | 0 | 0 | 0 | 0 | 0 | 0 | 0 | 29 | 2 | 0 | 0 | 0 |
| CM111806.1 | 22 | 0 | 0 | 3 | 0 | 2 | 1 | 2706 | 2713 | 111 | 105 | 1 | 0 | 8 | 9 | 0 | 0 | 0 | 0 | 0 | 0 | 0 | 0 | 0 | 0 | 0 | 0 | 0 | 0 | 0 | 0 | 0 | 0 | 0 | 0 | 0 |
| CM111791.1 | 7 | 0 | 3 | 6 | 0 | 21 | 3 | 2184 | 2171 | 534 | 682 | 132 | 0 | 47 | 97 | 0 | 0 | 0 | 1 | 0 | 0 | 0 | 0 | 0 | 0 | 0 | 0 | 0 | 0 | 0 | 0 | 32 | 1 | 0 | 0 | 0 |
| CM111796.1 | 12 | 1 | 1 | 7 | 0 | 1 | 3 | 0 | 40 | 3367 | 3550 | 224 | 2 | 4 | 28 | 0 | 0 | 1 | 0 | 0 | 0 | 0 | 0 | 0 | 0 | 0 | 0 | 0 | 0 | 0 | 0 | 1 | 0 | 0 | 0 | 0 |
| CM111789.1 | 5 | 1 | 2 | 10 | 0 | 77 | 7 | 0 | 0 | 2871 | 2993 | 43 | 0 | 38 | 58 | 2 | 0 | 1 | 0 | 0 | 0 | 0 | 0 | 0 | 0 | 0 | 0 | 0 | 0 | 0 | 0 | 0 | 0 | 12 | 9 | 13 |
| CM111807.1 | X | 2 | 0 | 6 | 0 | 81 | 18 | 0 | 3 | 2157 | 2291 | 84 | 1 | 44 | 66 | 0 | 0 | 99 | 112 | 0 | 1 | 0 | 0 | 0 | 0 | 0 | 0 | 0 | 0 | 0 | 1 | 0 | 0 | 0 | 0 | 0 |
| CM111805.1 | 21 | 0 | 1 | 9 | 0 | 72 | 4 | 0 | 1 | 2006 | 2133 | 80 | 0 | 64 | 119 | 0 | 0 | 0 | 0 | 0 | 0 | 0 | 0 | 0 | 0 | 0 | 0 | 0 | 0 | 0 | 0 | 13 | 1 | 0 | 0 | 0 |
| CM111794.1 | 10 | 21 | 3 | 4 | 0 | 38 | 5 | 1 | 0 | 1835 | 2002 | 114 | 1 | 34 | 118 | 0 | 1 | 2 | 0 | 0 | 21 | 23 | 19 | 18 | 18 | 9 | 9 | 9 | 11 | 11 | 11 | 0 | 1 | 0 | 0 | 0 |
| CM111790.1 | 6 | 1 | 4 | 6 | 0 | 4 | 11 | 0 | 1 | 5 | 150 | 3341 | 3173 | 41 | 106 | 1 | 0 | 0 | 0 | 0 | 0 | 0 | 0 | 0 | 0 | 0 | 0 | 0 | 1 | 0 | 0 | 2 | 3 | 0 | 0 | 0 |

Note: columns C to N show subSFs of SF1, O to AF show OWM-specific monomers of SFs 2-6 as defined in Table S1, and columns AG to AK show the monomers of ancient centromeric layers common to apes, OWM, and NWM[10,39].

#### SubSF competition in active arrays

Manual examination of the tracks revealed that interlayers were common in the marmoset genome (Table SN3) and revealed the following patterns:

(1) Previously, the most common active array in marmoset was SF1-5 (16 cens); however, in all but 5 cases, it was replaced by other subSFs: SF1-1 (8 cases), SF1-4 (3 cases).

(2) Cen16 was originally SF1-1, but it was replaced by SF1-4.

(3) Also, in one case, SF1-3 was replaced by SF1-2.

Note that all but one cases where SF1-1 has won occurred in acrocentric chromosomes, which are potentially rRNA-bearing and have a very similar architecture of the short arms, which possibly makes them prone to non-homologous exchange of the short arms, as it was proposed in humans[40,41]. It seems that, in marmoset, the history of events at the centromere may also be shared, or that centromeres (or parts of them) could similarly be exchanged between non-homologous acrocentrics. Thus, multiple cases of SF1-5 replacement by SF1-1 may not be independent events but a multiplication of just one or two original replacements. Additionally, one may speculate that SF1-5, which formed most of the original S3S4 centromeres, appeared to be somehow weaker than the other SubSFs, which could eventually replace it completely. Finally, various mixes that usually occupy the very tips of the active arrays are likely to be earlier subSFs that may be resolved in the future by adding yet more HMMs, which would provide a closer fit and more accurate recognition. For instance, in chr13, these mixed flanks of the active array are especially large (~100kb) on both sides. The S4 monomer is always identified as S4-5, and S3 monomers are identified variably as S3-1 or S3-3.

**Table SN3. Active subSF interlayers in marmoset centromeres (subSF competition).**

| chrom | left flank | core | right flank |
|---|---|---|---|
| chr1 | SF1-5 | SF1-4 | - |
| chr2 | SF1-5 | SF1-4 | - |
| chr3 | - | SF1-1 | SF1-5 |

| | | | |
|---|---|---|---|
| chr4 | - | SF1-2 | - |
| chr5 | - | SF1-5 | - |
| chr6 | - | SF1-6 | - |
| chr7 | - | SF1-4 | - |
| chr8 | - | SF1-3 | - |
| chr9 | - | SF1-2 | - |
| chr10 | - | SF1-5 | - |
| chr11 | SF1-2 | SF1-3 | SF1-2 |
| chr12 | - | SF1-5 | - |
| chr13 | - | SF1-3 | - |
| chr14 | - | SF1-1 | SF1-5 |
| chr15 | SF1-5 | SF1-1 | SF1-5 |
| chr16 | SF1-1 | SF1-4 | SF1-1 |
| chr17 | SF1-5 | SF1-1 | SF1-5 |
| chr18 | SF1-5 | SF1-1 | SF1-5 |
| chr19 | SF1-5 | SF1-1 | SF1-5 |
| chr20 | - | SF1-1 | SF1-5 |
| chr21 | - | SF1-5 | - |
| chr22 | SF1-5 | SF1-4 | - |
| chrX | - | SF1-4 | - |
| chrY | - | SF1-1 | SF1-5 |

#### Chromosomal specificity of S3S4 dimers within subSFs.

500 random dimer sequences extracted from active arrays of each chromosome were then used to derive chromosome-specific consensus sequences. Active arrays were defined as the active subSF arrays as annotated in the CenSat annotation track. Additionally, all 150 SF1-1 monomers from both cen16 flanking inactive arrays (separately) were added to this dataset. The resulting phylogenetic tree of consensus sequences is shown in Fig. **SN6** (the SF1-1 part) and in full in Fig. **6B** in the main text.

6 major branches corresponding to 6 subSFs can be seen. Most chromosomes have somewhat different consensus sequences, indicating a significant degree of specificity with some notable exceptions. Cens 11 and 13 have identical consensus dimers, and in SF1-1, just 2 dimers cover all centromeres, with dimer 1-1A forming centromeres in chrs 15, 17-19, and Y (all rDNA-bearing acrocentrics) and dimer 1-1B in chrs 14, 20, and 3 **(**Fig. **SN6)**. Of the latter, chromosomes 14 and 20 are acrocentric, and chromosome 14 is shown to occasionally bear rDNA. Inactive dimer from cen16 appears as a distant outlier. Note that chromosomes 16 and 3 are subtelocentrics with the shortest short arms, so they could be easily included in the broader acrocentric group. Thus, it would appear that all acrocentrics originally had, or currently have, SF1-1 centromeres. Moreover, the interlayer history indicates that they all (except chr16) had SF1-5

centromeres, which were then replaced by SF1-1. Overall, it seems plausible that the dead array in cen16 represents the ancestral sequence of SF1-1, which has spread from there to all acrocentrics, including chr3, and has developed 2 distinct variants along the way. One of these (1-1A) is firmly associated with rDNA, and the other is loosely associated.

To gain further insight into the chromosomal specificity of the active dimers, we extracted 500 active dimers from each chromosome, aligned them, and converted them into HMMs, which were used to annotate the assembly. The results indicated a high degree of chromosomal specificity (except for subSF1-1) as shown in Fig. **6C (right)** in the main text. Annotation statistics are shown in Table **SN4**.

**Table SN4. Chromosome-specific DimHap counts in marmoset active centromeres.**

| | dh_1 | dh_2 | dh_3 | dh_4 | dh_5 | dh_6 | dh_7 | dh_8 | dh_9 | dh_10 | dh_11 | dh_12 | dh_13 | dh_14 | dh_15 | dh_16 | dh_17 | dh_18 | dh_19 | dh_20 | dh_21 | dh_22 | dh_X | dh_Y |
|---|---|---|---|---|---|---|---|---|---|---|---|---|---|---|---|---|---|---|---|---|---|---|---|---|
| chr1 | 0.48 | 0.00 | 0.00 | 0.00 | 0.00 | 0.00 | 0.00 | 0.00 | 0.00 | 0.00 | 0.00 | 0.00 | 0.00 | 0.00 | 0.00 | 0.44 | 0.00 | 0.00 | 0.00 | 0.00 | 0.00 | 0.07 | 0.00 | 0.00 |
| chr2 | 0.00 | 1.00 | 0.00 | 0.00 | 0.00 | 0.00 | 0.00 | 0.00 | 0.00 | 0.00 | 0.00 | 0.00 | 0.00 | 0.00 | 0.00 | 0.00 | 0.00 | 0.00 | 0.00 | 0.00 | 0.00 | 0.00 | 0.00 | 0.00 |
| chr3 | 0.00 | 0.00 | 0.55 | 0.00 | 0.00 | 0.00 | 0.00 | 0.00 | 0.00 | 0.00 | 0.00 | 0.00 | 0.00 | 0.40 | 0.00 | 0.00 | 0.00 | 0.01 | 0.00 | 0.04 | 0.00 | 0.00 | 0.00 | 0.00 |
| chr4 | 0.00 | 0.00 | 0.00 | 0.97 | 0.00 | 0.00 | 0.00 | 0.00 | 0.03 | 0.00 | 0.00 | 0.00 | 0.00 | 0.00 | 0.00 | 0.00 | 0.00 | 0.00 | 0.00 | 0.00 | 0.00 | 0.00 | 0.00 | 0.00 |
| chr5 | 0.00 | 0.00 | 0.00 | 0.00 | 0.99 | 0.00 | 0.00 | 0.00 | 0.00 | 0.01 | 0.00 | 0.00 | 0.00 | 0.00 | 0.00 | 0.00 | 0.00 | 0.00 | 0.00 | 0.00 | 0.00 | 0.00 | 0.00 | 0.00 |
| chr6 | 0.00 | 0.00 | 0.00 | 0.00 | 0.00 | 1.00 | 0.00 | 0.00 | 0.00 | 0.00 | 0.00 | 0.00 | 0.00 | 0.00 | 0.00 | 0.00 | 0.00 | 0.00 | 0.00 | 0.00 | 0.00 | 0.00 | 0.00 | 0.00 |
| chr7 | 0.00 | 0.00 | 0.00 | 0.00 | 0.00 | 0.00 | 0.96 | 0.00 | 0.00 | 0.00 | 0.00 | 0.00 | 0.00 | 0.00 | 0.00 | 0.04 | 0.00 | 0.00 | 0.00 | 0.00 | 0.00 | 0.00 | 0.00 | 0.00 |
| chr8 | 0.00 | 0.00 | 0.00 | 0.00 | 0.00 | 0.00 | 0.00 | 0.97 | 0.00 | 0.00 | 0.03 | 0.00 | 0.00 | 0.00 | 0.00 | 0.00 | 0.00 | 0.00 | 0.00 | 0.00 | 0.00 | 0.00 | 0.00 | 0.00 |
| chr9 | 0.00 | 0.00 | 0.00 | 0.02 | 0.00 | 0.00 | 0.00 | 0.00 | 0.98 | 0.00 | 0.00 | 0.00 | 0.00 | 0.00 | 0.00 | 0.00 | 0.00 | 0.00 | 0.00 | 0.00 | 0.00 | 0.00 | 0.00 | 0.00 |
| chr10 | 0.00 | 0.00 | 0.00 | 0.00 | 0.00 | 0.00 | 0.00 | 0.00 | 0.00 | 0.99 | 0.00 | 0.00 | 0.00 | 0.00 | 0.00 | 0.00 | 0.00 | 0.00 | 0.00 | 0.00 | 0.00 | 0.00 | 0.01 | 0.00 |
| chr11 | 0.00 | 0.00 | 0.00 | 0.00 | 0.00 | 0.00 | 0.00 | 0.11 | 0.00 | 0.00 | 0.87 | 0.00 | 0.02 | 0.00 | 0.00 | 0.00 | 0.00 | 0.00 | 0.00 | 0.00 | 0.00 | 0.00 | 0.00 | 0.00 |
| chr12 | 0.00 | 0.00 | 0.00 | 0.00 | 0.01 | 0.00 | 0.00 | 0.00 | 0.00 | 0.00 | 0.00 | 0.98 | 0.00 | 0.00 | 0.00 | 0.00 | 0.00 | 0.00 | 0.00 | 0.00 | 0.00 | 0.00 | 0.00 | 0.00 |
| chr13 | 0.00 | 0.00 | 0.00 | 0.00 | 0.00 | 0.00 | 0.00 | 0.00 | 0.00 | 0.00 | 0.19 | 0.00 | 0.81 | 0.00 | 0.00 | 0.00 | 0.00 | 0.00 | 0.00 | 0.00 | 0.00 | 0.00 | 0.00 | 0.00 |
| chr14 | 0.00 | 0.00 | 0.02 | 0.00 | 0.00 | 0.00 | 0.00 | 0.00 | 0.00 | 0.00 | 0.00 | 0.00 | 0.00 | 0.89 | 0.00 | 0.00 | 0.00 | 0.00 | 0.00 | 0.09 | 0.00 | 0.00 | 0.00 | 0.00 |
| chr15 | 0.00 | 0.00 | 0.01 | 0.00 | 0.00 | 0.00 | 0.00 | 0.00 | 0.00 | 0.00 | 0.00 | 0.00 | 0.00 | 0.06 | 0.56 | 0.00 | 0.16 | 0.12 | 0.08 | 0.01 | 0.00 | 0.00 | 0.00 | 0.00 |
| chr16 | 0.00 | 0.00 | 0.00 | 0.00 | 0.00 | 0.00 | 0.02 | 0.00 | 0.00 | 0.00 | 0.00 | 0.00 | 0.00 | 0.00 | 0.00 | 0.97 | 0.00 | 0.00 | 0.00 | 0.00 | 0.00 | 0.01 | 0.00 | 0.00 |
| chr17 | 0.00 | 0.00 | 0.01 | 0.00 | 0.00 | 0.00 | 0.00 | 0.00 | 0.00 | 0.00 | 0.00 | 0.00 | 0.00 | 0.00 | 0.26 | 0.00 | 0.60 | 0.07 | 0.06 | 0.00 | 0.00 | 0.00 | 0.00 | 0.00 |
| chr18 | 0.00 | 0.00 | 0.04 | 0.00 | 0.00 | 0.00 | 0.00 | 0.00 | 0.00 | 0.00 | 0.00 | 0.00 | 0.00 | 0.06 | 0.15 | 0.00 | 0.13 | 0.52 | 0.08 | 0.02 | 0.00 | 0.00 | 0.00 | 0.00 |
| chr19 | 0.00 | 0.00 | 0.01 | 0.00 | 0.00 | 0.00 | 0.00 | 0.00 | 0.00 | 0.00 | 0.00 | 0.00 | 0.00 | 0.00 | 0.23 | 0.00 | 0.11 | 0.09 | 0.56 | 0.00 | 0.00 | 0.00 | 0.00 | 0.00 |
| chr20 | 0.00 | 0.00 | 0.03 | 0.00 | 0.00 | 0.00 | 0.00 | 0.00 | 0.00 | 0.00 | 0.00 | 0.00 | 0.00 | 0.46 | 0.00 | 0.00 | 0.00 | 0.02 | 0.00 | 0.48 | 0.00 | 0.00 | 0.00 | 0.00 |
| chr21 | 0.00 | 0.00 | 0.00 | 0.00 | 0.01 | 0.00 | 0.00 | 0.00 | 0.00 | 0.00 | 0.00 | 0.00 | 0.00 | 0.00 | 0.00 | 0.00 | 0.00 | 0.00 | 0.00 | 0.00 | 0.98 | 0.00 | 0.01 | 0.00 |
| chr22 | 0.00 | 0.00 | 0.00 | 0.00 | 0.00 | 0.00 | 0.00 | 0.00 | 0.00 | 0.00 | 0.00 | 0.00 | 0.00 | 0.00 | 0.00 | 0.06 | 0.00 | 0.00 | 0.00 | 0.00 | 0.00 | 0.93 | 0.00 | 0.00 |
| chrX | 0.00 | 0.00 | 0.00 | 0.00 | 0.00 | 0.00 | 0.00 | 0.00 | 0.00 | 0.00 | 0.00 | 0.00 | 0.00 | 0.00 | 0.00 | 0.00 | 0.00 | 0.00 | 0.00 | 0.00 | 0.01 | 0.00 | 0.99 | 0.00 |
| chrY | 0.00 | 0.00 | 0.01 | 0.00 | 0.00 | 0.00 | 0.00 | 0.00 | 0.00 | 0.00 | 0.00 | 0.00 | 0.00 | 0.01 | 0.21 | 0.00 | 0.07 | 0.11 | 0.56 | 0.01 | 0.00 | 0.00 | 0.00 | 0.01 |

It can be seen that 21 centromeres exhibit a significant degree of chromosomal specificity (more than 50% of dimers belong to their "own" type). The rest form several groups, of which the 2 largest ones cover SF1-1 in acrocentric rDNA-bearing chromosomes (as discussed in detail in the main text). The homogeneity of these groups may be supported by selection and may represent a special case. The other groups are pairs: 11/13 and 1/16; they may represent occasional recent cases of interchromosomal sequence exchange/amplification[38] that did not have enough time to diverge sufficiently. Such cases are also known in humans (e.g., 1/5/19 triplet).

**Fig. SN6. The phylogeny of SF1-1 chromosome-specific dimers in marmoset.** The SF1-1 sequences from 2 different flanks on chr16 are shown separately to demonstrate their similar phylogenetic positions. The whole tree is shown in Fig. **6B**.

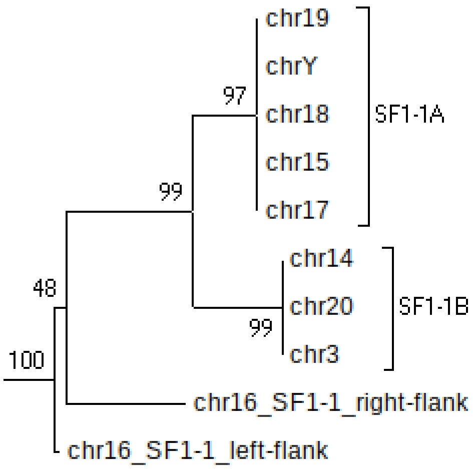


#### Kmer analysis of chromosomal specificity and homolog differences

Sequences of active arrays from every centromere across 4 chromosome complements were analysed using ClusterSourMash (https://github.com/fedorrik/ClusterSourMash), which compares the kmer spectra and copy numbers of the input sequences. The output was clustered to produce a UPGMA dendrogram (Fig. SN7). The dendrogram faithfully reflected the subSF structure (differently colored branches). In all cases, the same chromosomes were grouped together, except for acrocentric chromosomes in the SF1-1 group. Mixing in this branch could reflect the history of non-homologous exchanges between rDNA-bearing acrocentrics (see Discussion in the main text).

**Fig. SN7. ClusterSourMash analysis of active (S3S4) arrays in marmoset centromeres.**

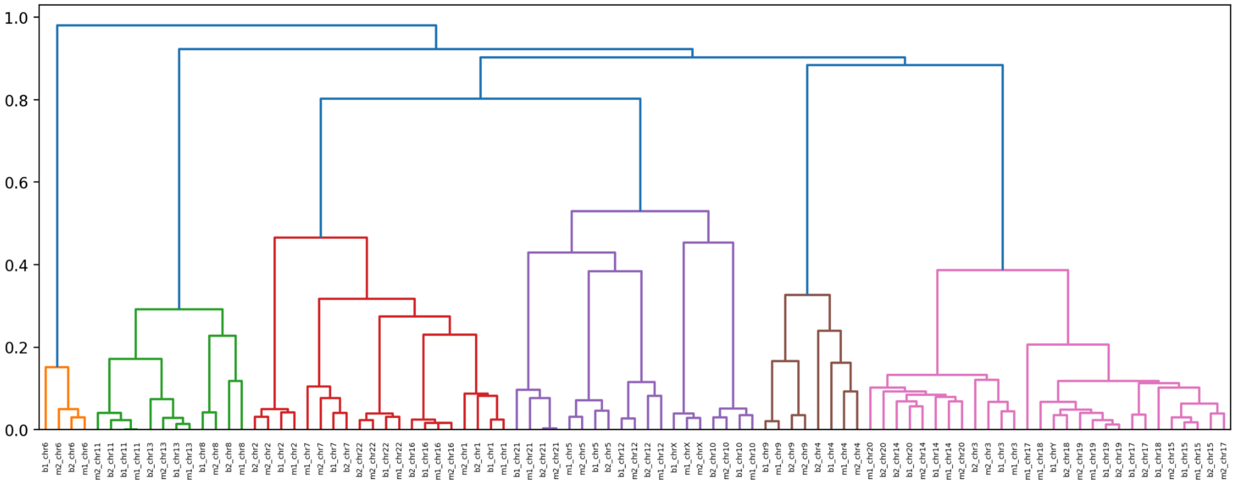

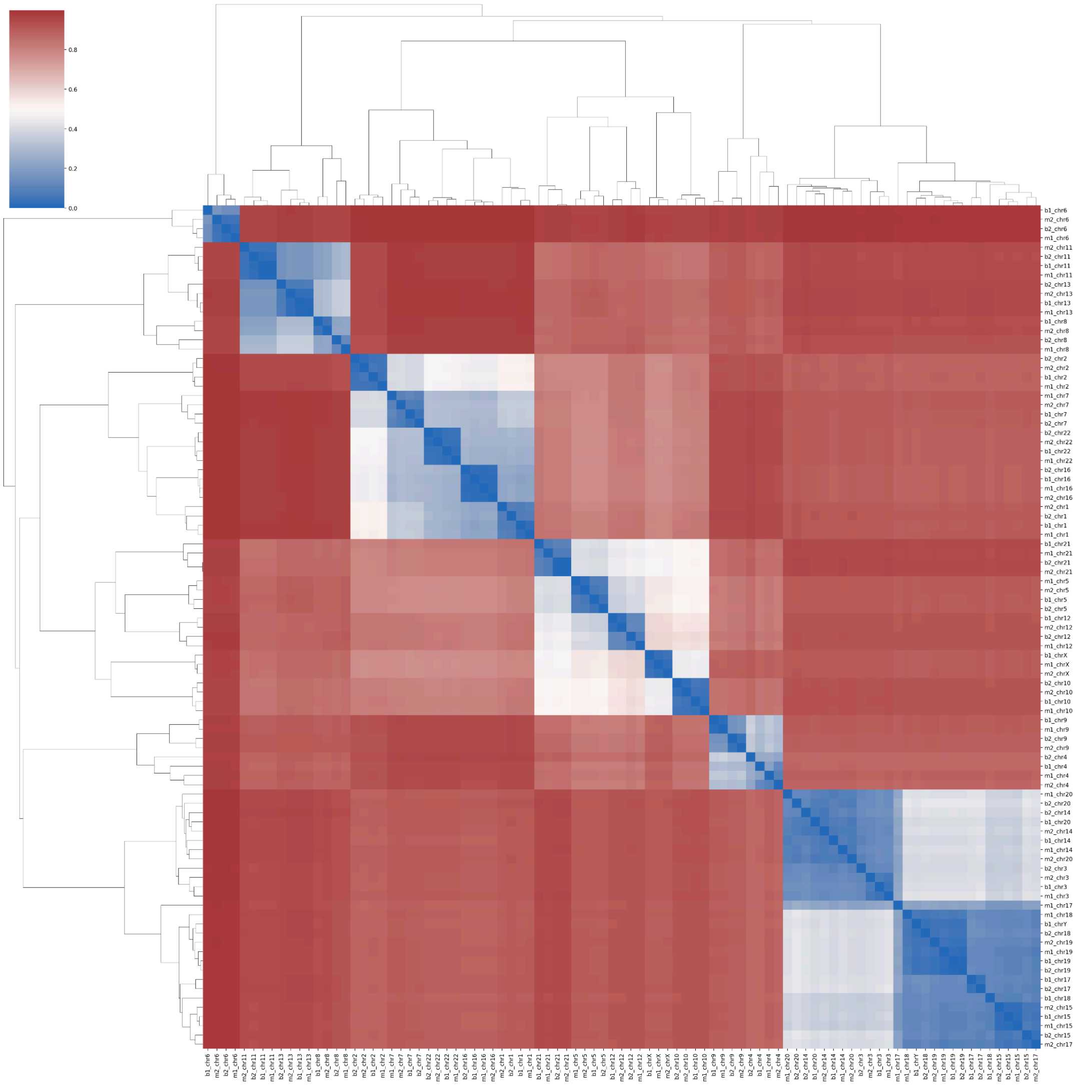


b1 = calJac240_primary
b2 = calJac240_alternate
m1 = calJac220_primary
m2 = calJac220_alternate

Note that ClusterSourMash reveals considerable differences between 4 copies of a homologous chromosome in a number of cases. Such differences reflect centromere haplotypes (cenhaps), which are stably maintained as centromeric lineages because of repressed meiotic recombination in and around centromeres[42]. For example, one copy of cen6 is dramatically different from the other 3 (Fig. **SN7, left edge**). Also, 2 distinct types of cens 18 and 19 are notable.

We next tried to generate chromosome-specific kmers for each marmoset centromere. The S3S4 dimer arrays from *GCA_049354665.1* assembly were used to obtain chromosome-specific kmers as previously described for the macaque genome[14]. Specificity parameters were: (1) count in a target chromosome assembly >= 30, (2) count in any other chromosome <= 10% of the target count. The lists of kmers obtained were examined, and one kmer per chromosome was selected that had an optimal copy number vs. background ratio (aiming for a higher copy number on the lowest background). The counts of these selected kmers are shown in **Table SN5**.

**Table SN5. The counts of selected chromosome-specific kmers in marmoset centromeres.**

| active SF | sum cnt | kmer | chr1 | chr2 | chr3 | chr4 | chr5 | chr6 | chr7 | chr8 | chr9 | chr10 | chr11 | chr12 | chr13 | chr14 | chr15 | chr16 | chr17 | chr18 | chr19 | chr20 | chr21 | chr22 | chrX | chrY |
|---|---|---|---|---|---|---|---|---|---|---|---|---|---|---|---|---|---|---|---|---|---|---|---|---|---|---|
| SF1-4 | 899 | b1_chr1-1 | 0.98 | 0.00 | 0.00 | 0.00 | 0.00 | 0.00 | 0.00 | 0.00 | 0.00 | 0.00 | 0.00 | 0.00 | 0.00 | 0.00 | 0.00 | 0.01 | 0.00 | 0.00 | 0.00 | 0.00 | 0.00 | 0.00 | 0.00 | 0.00 |
| SF1-4 | 538 | b1_chr2-1 | 0.00 | 0.97 | 0.00 | 0.00 | 0.00 | 0.00 | 0.00 | 0.00 | 0.00 | 0.00 | 0.00 | 0.00 | 0.00 | 0.00 | 0.00 | 0.01 | 0.00 | 0.00 | 0.00 | 0.00 | 0.00 | 0.02 | 0.00 | 0.00 |
| SF1-1 | 466 | b1_chr3-1 | 0.00 | 0.00 | 0.99 | 0.00 | 0.00 | 0.00 | 0.00 | 0.00 | 0.00 | 0.00 | 0.00 | 0.00 | 0.00 | 0.00 | 0.00 | 0.00 | 0.00 | 0.00 | 0.00 | 0.00 | 0.00 | 0.00 | 0.00 | 0.00 |
| SF1-2 | 357 | b1_chr4-1 | 0.00 | 0.00 | 0.00 | 0.97 | 0.00 | 0.00 | 0.00 | 0.00 | 0.03 | 0.00 | 0.00 | 0.00 | 0.00 | 0.00 | 0.00 | 0.00 | 0.00 | 0.00 | 0.00 | 0.00 | 0.00 | 0.00 | 0.00 | 0.00 |
| SF1-5 | 564 | b1_chr5-1 | 0.00 | 0.00 | 0.00 | 0.00 | 0.98 | 0.00 | 0.00 | 0.00 | 0.00 | 0.00 | 0.00 | 0.02 | 0.00 | 0.00 | 0.00 | 0.00 | 0.00 | 0.00 | 0.00 | 0.00 | 0.00 | 0.00 | 0.00 | 0.00 |
| SF1-6 | 2569 | b1_chr6-1 | 0.00 | 0.00 | 0.00 | 0.00 | 0.00 | 0.98 | 0.00 | 0.00 | 0.00 | 0.00 | 0.00 | 0.00 | 0.00 | 0.00 | 0.00 | 0.00 | 0.00 | 0.00 | 0.00 | 0.00 | 0.00 | 0.00 | 0.01 | 0.00 |
| SF1-4 | 598 | b1_chr7-1 | 0.01 | 0.00 | 0.00 | 0.00 | 0.00 | 0.00 | 0.98 | 0.00 | 0.00 | 0.00 | 0.00 | 0.00 | 0.00 | 0.00 | 0.00 | 0.00 | 0.00 | 0.00 | 0.00 | 0.00 | 0.00 | 0.01 | 0.00 | 0.00 |
| SF1-3 | 1963 | b1_chr8-1 | 0.00 | 0.00 | 0.00 | 0.00 | 0.00 | 0.00 | 0.00 | 1.00 | 0.00 | 0.00 | 0.00 | 0.00 | 0.00 | 0.00 | 0.00 | 0.00 | 0.00 | 0.00 | 0.00 | 0.00 | 0.00 | 0.00 | 0.00 | 0.00 |
| SF1-2 | 2359 | b1_chr9-1 | 0.00 | 0.00 | 0.00 | 0.04 | 0.00 | 0.00 | 0.00 | 0.00 | 0.96 | 0.00 | 0.00 | 0.00 | 0.00 | 0.00 | 0.00 | 0.00 | 0.00 | 0.00 | 0.00 | 0.00 | 0.00 | 0.00 | 0.00 | 0.00 |
| SF1-5 | 857 | b1_chr10-1 | 0.00 | 0.00 | 0.03 | 0.00 | 0.00 | 0.00 | 0.00 | 0.00 | 0.00 | 0.96 | 0.00 | 0.00 | 0.00 | 0.00 | 0.00 | 0.00 | 0.00 | 0.00 | 0.00 | 0.00 | 0.00 | 0.00 | 0.00 | 0.00 |
| SF1-3 | 412 | b1_chr11-1 | 0.00 | 0.00 | 0.00 | 0.00 | 0.00 | 0.00 | 0.00 | 0.00 | 0.00 | 0.00 | 1.00 | 0.00 | 0.00 | 0.00 | 0.00 | 0.00 | 0.00 | 0.00 | 0.00 | 0.00 | 0.00 | 0.00 | 0.00 | 0.00 |
| SF1-5 | 1975 | b1_chr12-1 | 0.00 | 0.00 | 0.00 | 0.00 | 0.00 | 0.00 | 0.08 | 0.00 | 0.00 | 0.00 | 0.00 | 0.90 | 0.00 | 0.00 | 0.01 | 0.00 | 0.00 | 0.01 | 0.00 | 0.00 | 0.00 | 0.00 | 0.00 | 0.00 |
| SF1-3 | 1278 | b1_chr13-1 | 0.00 | 0.00 | 0.00 | 0.00 | 0.00 | 0.00 | 0.00 | 0.02 | 0.00 | 0.00 | 0.00 | 0.00 | 0.97 | 0.00 | 0.00 | 0.00 | 0.00 | 0.00 | 0.00 | 0.00 | 0.00 | 0.00 | 0.00 | 0.00 |
| SF1-1 | 96 | b1_chr14-1 | 0.00 | 0.00 | 0.00 | 0.00 | 0.00 | 0.00 | 0.00 | 0.00 | 0.00 | 0.00 | 0.00 | 0.00 | 0.00 | 0.99 | 0.00 | 0.00 | 0.00 | 0.00 | 0.00 | 0.01 | 0.00 | 0.00 | 0.00 | 0.00 |
| SF1-1 | 117 | b1_chr15-1 | 0.00 | 0.00 | 0.00 | 0.00 | 0.00 | 0.00 | 0.00 | 0.00 | 0.00 | 0.00 | 0.00 | 0.00 | 0.00 | 0.00 | 0.98 | 0.00 | 0.00 | 0.01 | 0.01 | 0.00 | 0.00 | 0.00 | 0.00 | 0.00 |
| SF1-4 | 569 | b1_chr16-1 | 0.00 | 0.00 | 0.00 | 0.00 | 0.00 | 0.00 | 0.00 | 0.00 | 0.00 | 0.00 | 0.00 | 0.00 | 0.00 | 0.00 | 0.00 | 1.00 | 0.00 | 0.00 | 0.00 | 0.00 | 0.00 | 0.00 | 0.00 | 0.00 |
| SF1-1 | 172 | b1_chr17-1 | 0.00 | 0.00 | 0.00 | 0.00 | 0.00 | 0.00 | 0.00 | 0.00 | 0.00 | 0.00 | 0.00 | 0.00 | 0.04 | 0.00 | 0.00 | 0.00 | 0.96 | 0.00 | 0.00 | 0.00 | 0.00 | 0.00 | 0.00 | 0.00 |
| SF1-1 | 323 | b1_chr18-1 | 0.00 | 0.00 | 0.00 | 0.00 | 0.00 | 0.00 | 0.00 | 0.00 | 0.00 | 0.00 | 0.00 | 0.00 | 0.00 | 0.00 | 0.01 | 0.00 | 0.00 | 0.97 | 0.01 | 0.00 | 0.00 | 0.00 | 0.00 | 0.01 |
| SF1-1 | 39 | b1_chr19-1 | 0.00 | 0.00 | 0.00 | 0.00 | 0.00 | 0.00 | 0.00 | 0.00 | 0.00 | 0.00 | 0.00 | 0.00 | 0.00 | 0.00 | 0.03 | 0.00 | 0.00 | 0.00 | 0.97 | 0.00 | 0.00 | 0.00 | 0.00 | 0.00 |
| SF1-1 | 34 | b1_chr20-1 | 0.00 | 0.00 | 0.00 | 0.00 | 0.00 | 0.00 | 0.00 | 0.00 | 0.00 | 0.00 | 0.00 | 0.00 | 0.00 | 0.00 | 0.00 | 0.00 | 0.00 | 0.00 | 0.00 | 1.00 | 0.00 | 0.00 | 0.00 | 0.00 |
| SF1-5 | 278 | b1_chr21-1 | 0.00 | 0.00 | 0.00 | 0.00 | 0.00 | 0.00 | 0.00 | 0.00 | 0.00 | 0.00 | 0.00 | 0.00 | 0.00 | 0.00 | 0.00 | 0.00 | 0.00 | 0.00 | 0.00 | 0.00 | 0.99 | 0.00 | 0.00 | 0.00 |
| SF1-4 | 1611 | b1_chr22-1 | 0.00 | 0.00 | 0.00 | 0.00 | 0.00 | 0.00 | 0.00 | 0.00 | 0.00 | 0.00 | 0.00 | 0.00 | 0.00 | 0.00 | 0.00 | 0.00 | 0.00 | 0.00 | 0.00 | 0.00 | 0.00 | 1.00 | 0.00 | 0.00 |
| SF1-4 | 342 | b1_chrX-1 | 0.00 | 0.00 | 0.01 | 0.00 | 0.00 | 0.00 | 0.00 | 0.00 | 0.00 | 0.01 | 0.00 | 0.01 | 0.00 | 0.00 | 0.00 | 0.00 | 0.00 | 0.00 | 0.00 | 0.00 | 0.00 | 0.00 | 0.98 | 0.00 |
| SF1-1 | 80 | b1_chrY-1 | 0.00 | 0.00 | 0.00 | 0.00 | 0.00 | 0.00 | 0.00 | 0.00 | 0.00 | 0.00 | 0.00 | 0.00 | 0.00 | 0.00 | 0.00 | 0.00 | 0.00 | 0.00 | 0.00 | 0.03 | 0.00 | 0.00 | 0.00 | 0.98 |

Note: the sum cnt column shows the total count in the assembly. The values in the cells show the proportion of each marmoset chromosome.

Note that SF1-1 (acrocentric) centromeres have only relatively low-copy kmers, because they share almost identical dimers, but even they have some significant chromosome-specific expansions in their live arrays. For instance, in cenY this expansion is located towards the q-end of the active array (**Fig. SN8**). That correlates with observations of the general architecture of acrocentric centromeric arrays: their p-sides appear more uniform in structure, and the q-sides more chromosome-specific. However, in other acrocentric chromosomes, the expansions can be scattered or located in the middle, but they never span the extreme p-side of the array.

**Fig. SN8. A. A typical localization of cenY-specific kmer on the q-side of the centromere.** All cenY-specific kmers obtained are concentrated in the same region. The chromosome is flipped to achieve the correct orientation. B. SubSF tracks of acrocentric chromosomes. Note similar-looking p-arm ends within SF1-1A and SF1-1B groups, and differently looking q-arm ends (except for the 17/19 pair).

A.

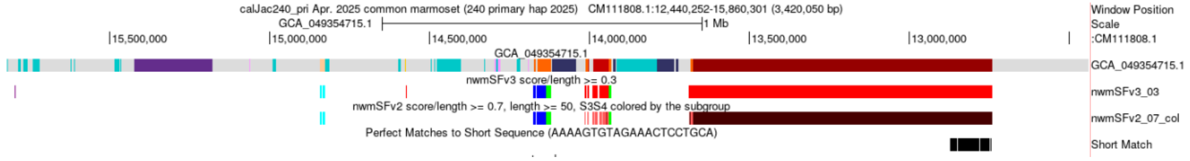


B.

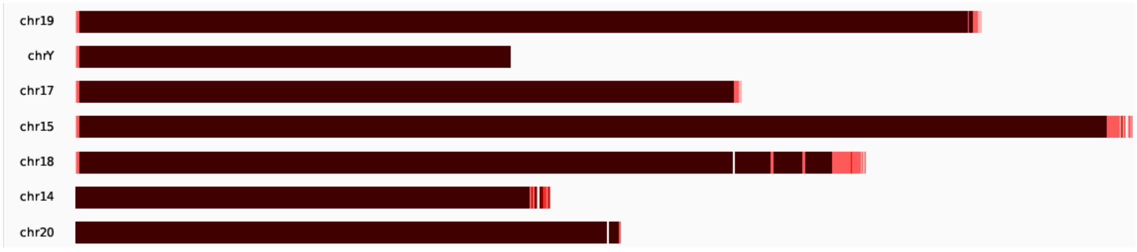

#### Divergence variation within the same dimhap array (chr14)

While examining the alignments of dimer sequences extracted from individual chromosomes, we have noted that some centromeres have more divergent dimers at the flanks and more homogeneous ones in the middle. Unexpectedly, this was not because the flanking regions were formed by slightly different dimhaps with different levels of divergence. The latter is well documented in humans and primates[10,31]. An example from chr14, which shows a virtual absence of specific clades in such arrays, is shown in Fig. **SN9**. It shows 3 parts of the same alignment taken from the middle of the array and from 2 flanks. On the flanks, which are much more divergent, one cannot see any significant clades that share the same colored consensus differences. In some cases, the same mutations appear in 2 or 3 dimers (usually located close together), but there are no consistent polymorphic sites.

**Fig. SN9. Alignment of the dimers from the chromosome 14 active region.**

Middle (cen14)

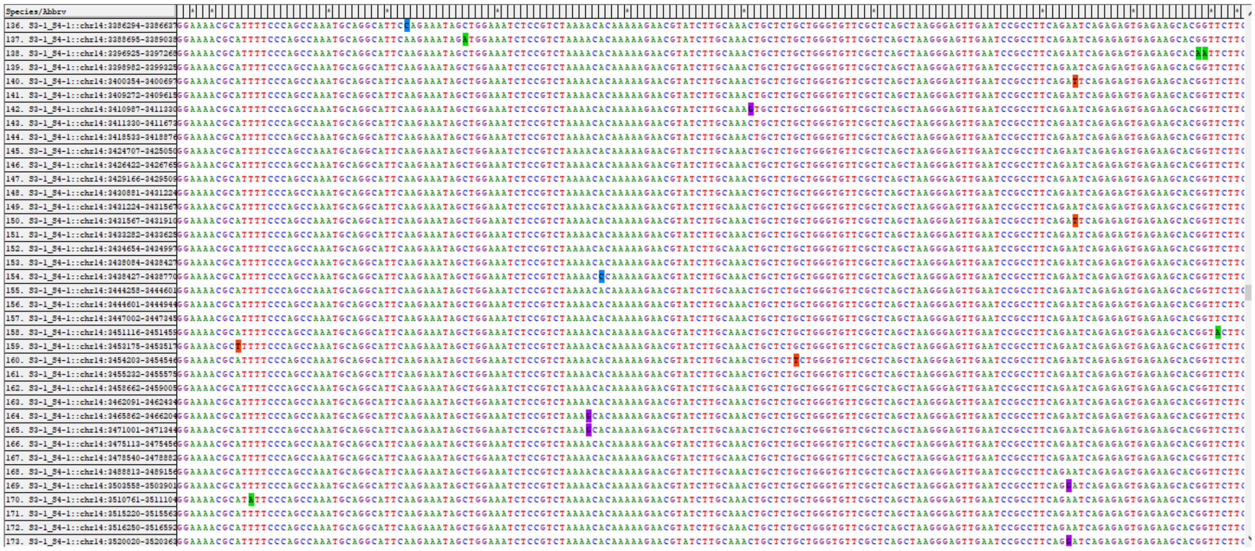

Top (cen14)

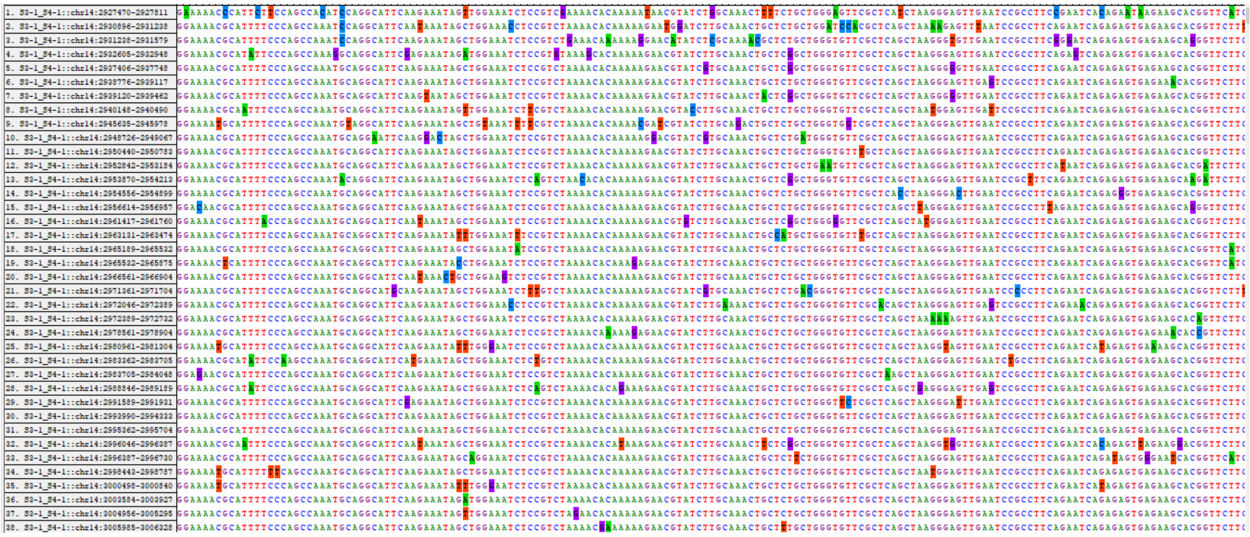

Bottom (cen14)

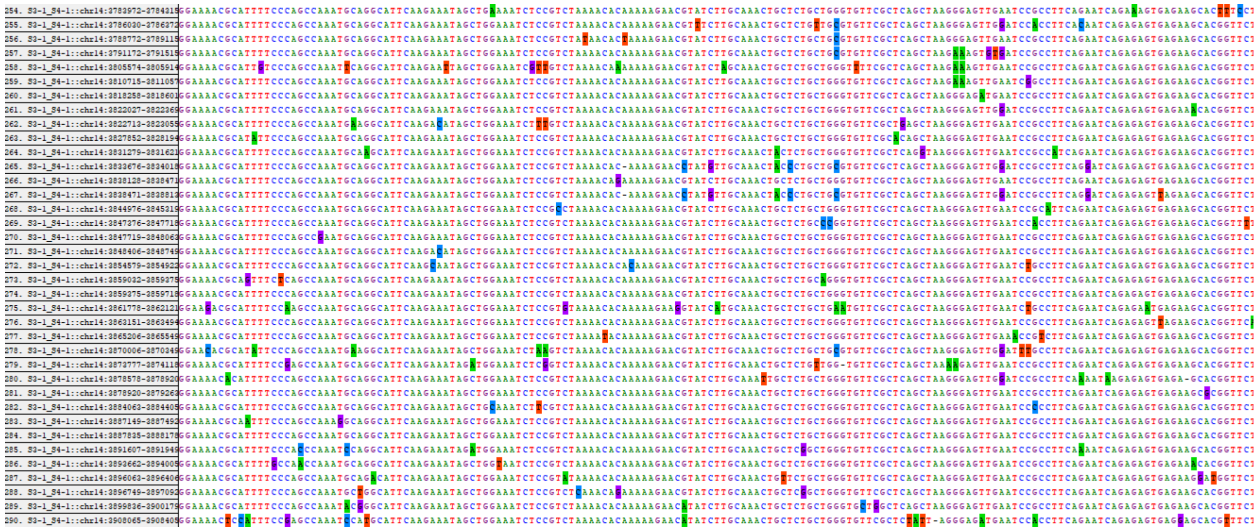

Accordingly, the consensus dimers derived from the flanks and the center, using various methods, were identical (Fig. **SN10**).

**Fig. SN10. Dimer similarity to intra-array consensus sequence along the length of the active array in chr14 (ASat divergence track). Localization of the CDR aligns with the high-identity region in centromere 14.**

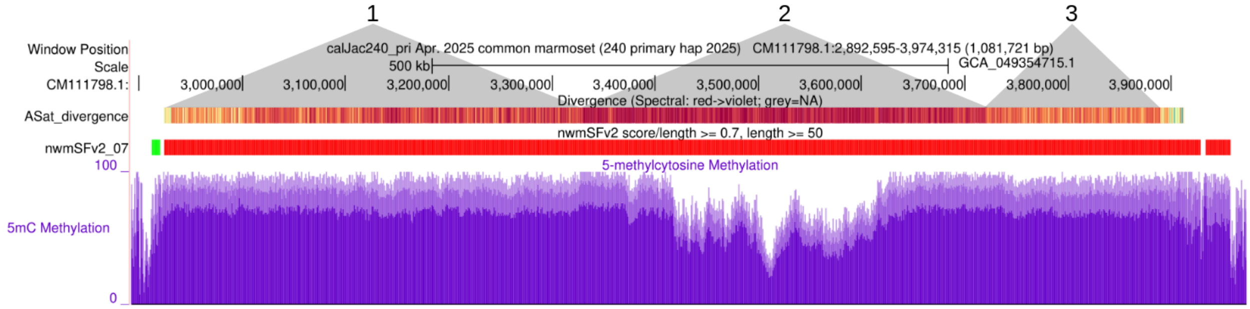


One can see 2 regions of reduced similarity on the flanks and a high-similarity region in the middle, as illustrated above. The average similarity in the regions was as follows:

1 0.967
2 0.986
3 0.949

Coordinates of the whole region and the 3 sub-regions were:

all CM111798.1:2,888,631-3,972,449
1 CM111798.1:3,001,263-3,320,032
2 CM111798.1:3,327,684-3,721,258
3 CM111798.1:3,724,659-3,886,169

Consensus sequences, derived from the 3 regions separately, were identical regardless of the method (simple majority or >50%). Note that the CDR, the methylation dip which marks the kinetochore location, aligns with the high-identity region.

#### AS HOR in cenX

The 28mer HOR was readily revealed with NTRprism and is clearly visible on the humSF track due to regularly spaced hits. CJ_cenXHOR kmer CAGAAAACGCATTTTTTCAGC hits once in the HOR and may be used to detect it (Fig. **SN11**). The HOR consists of 14 S3-5_S4-5 dimers. One can see that it occupies only part of the cenX active array; the rest has no HOR. Notably, the CDR overlaps the HOR region.

**Fig. SN11. Visualization of the 28mer HOR in marmoset cenX.**

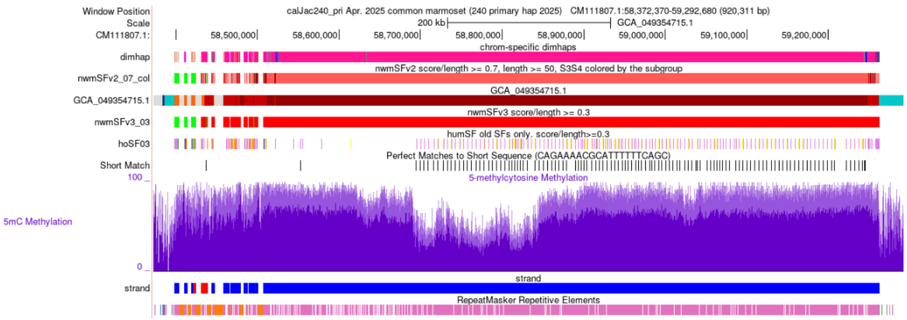


#### Strand reversals in the active arrays

In humans, active arrays in centromeres rarely contain inverted regions, and AS almost always remains on either the direct or reverse strand[10]. This is not the case in the flanking inactive AS regions, where inversions are common. Notably, in macaques, the inversions in active regions are much more frequent[14].

It appears that marmosets follow the human pattern. No inversions were found in the active regions, and they were frequent in the dead layers (Fig. **SN12**). For comparison, only one inversion is known in the active regions in humans (chr1)[10]. Also, there is an imbalance between direct and reverse orientations. In humans, about half of the centromeres have active regions in both direct and reverse orientations. The significance of this observation is unknown.

**Fig. SN12. Visualization of inversions in GCA_049354655.1_calJac240_alt (autosomes only).** In chr1 and chr21, AS goes on the reverse strand (red); one can see frequent reversals in the dead layers on the flanks. Not a single inversion is present in the central regions of centromeres, which are occupied by the active arrays.

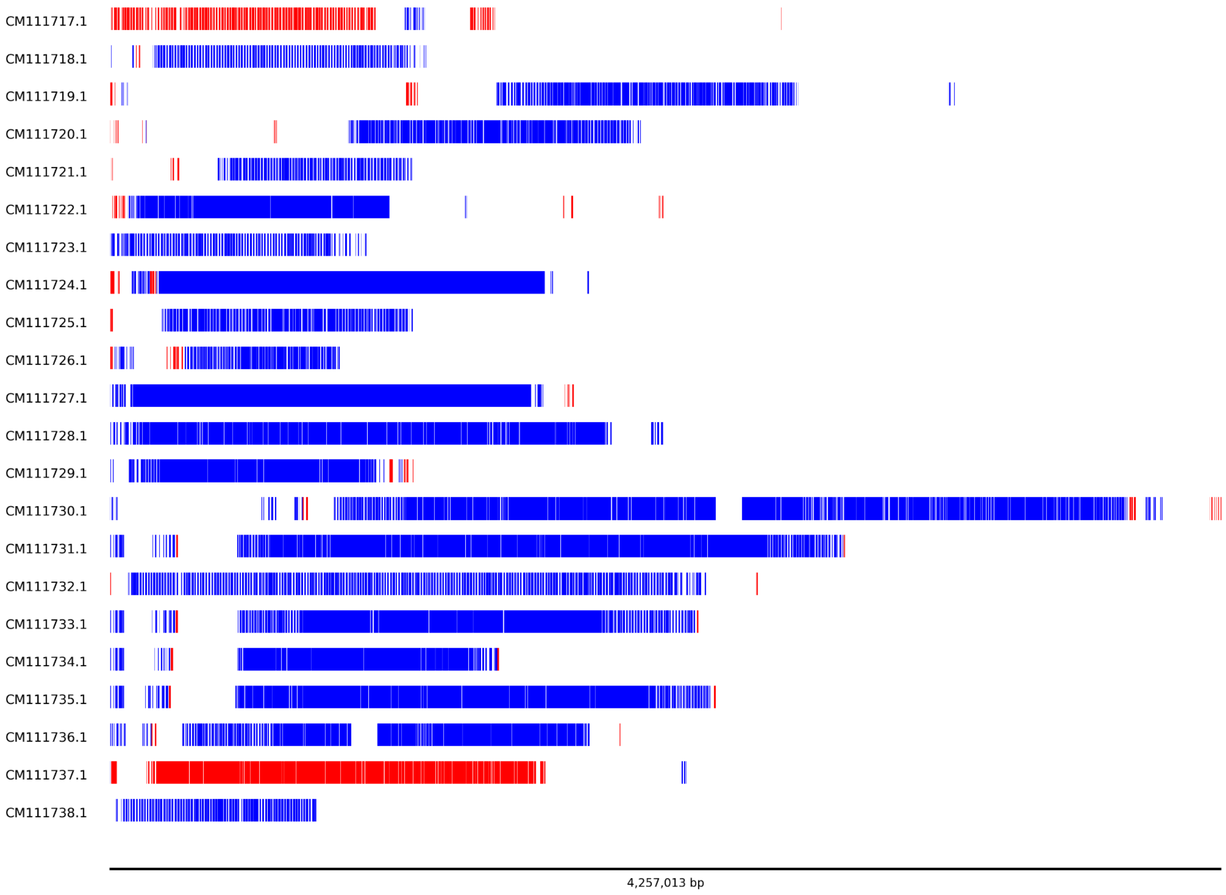


#### What were the centromeres of the Last Common Ancestor (LCA) of apes and NWM?

To answer this question, we (1) tried to annotate the NWM-specific arrays using ape-shared monomers as annotation standards, and (2) constructed and examined the all-encompassing tree of AS monomers to establish the phylogenetic relationships of the NWM-specific monomers with the ones shared with the ape lineage. The last shared monomer species was expected to form the centromeres of the hypothetical ancestral taxon.

The first test was performed using only the annotation standards for AS monomers typical of primates before the African apes, as classifying too distant monomer species is heavily affected by random coincidences. Thus, only the monomers for SF4 and older SFs were used, and w/o any filtering threshold, to achieve more complete coverage. Specifically, AS-SFs-hmmer3.0.290621.hmm were used to make an hmm file with "old" SFs only (Qa Pa Ta Ea Fa Ia Aa Ja Ba Ca Oa Na Ka Ha Ga FD La). The file was run against the GCA_0493546665.1 assembly. Both the unfiltered and 0.3-filtered versions were installed on the hub at https://genome.ucsc.edu/s/fedorrik/marmoset.

Under these conditions, the marmoset centromeres were mostly recognized as SF10 (Ba) and SF11 (Ja) monomers (Fig. **SN13**). However, other much younger species, which could not have possibly been ancestral to NWM monomers, were also present. Thus, this method, which worked very convincingly with the OWM centromeres[14], was not very effective for NWM centromeres, yielding only a very approximate answer. Presumably, that was because the NWM-specific repeats had split from the ape tree much earlier and had much more time to accumulate divergence. Hence, the classification was very much affected by coincidental mutation matching.

**Fig. SN13. Annotation of marmoset centromeres using only the standards shared with the ape lineage.** One can see that the prevailing colors are lilac (APE-SF11) and orange (APE-SF10). The genome-wide quantification is shown at the bottom.

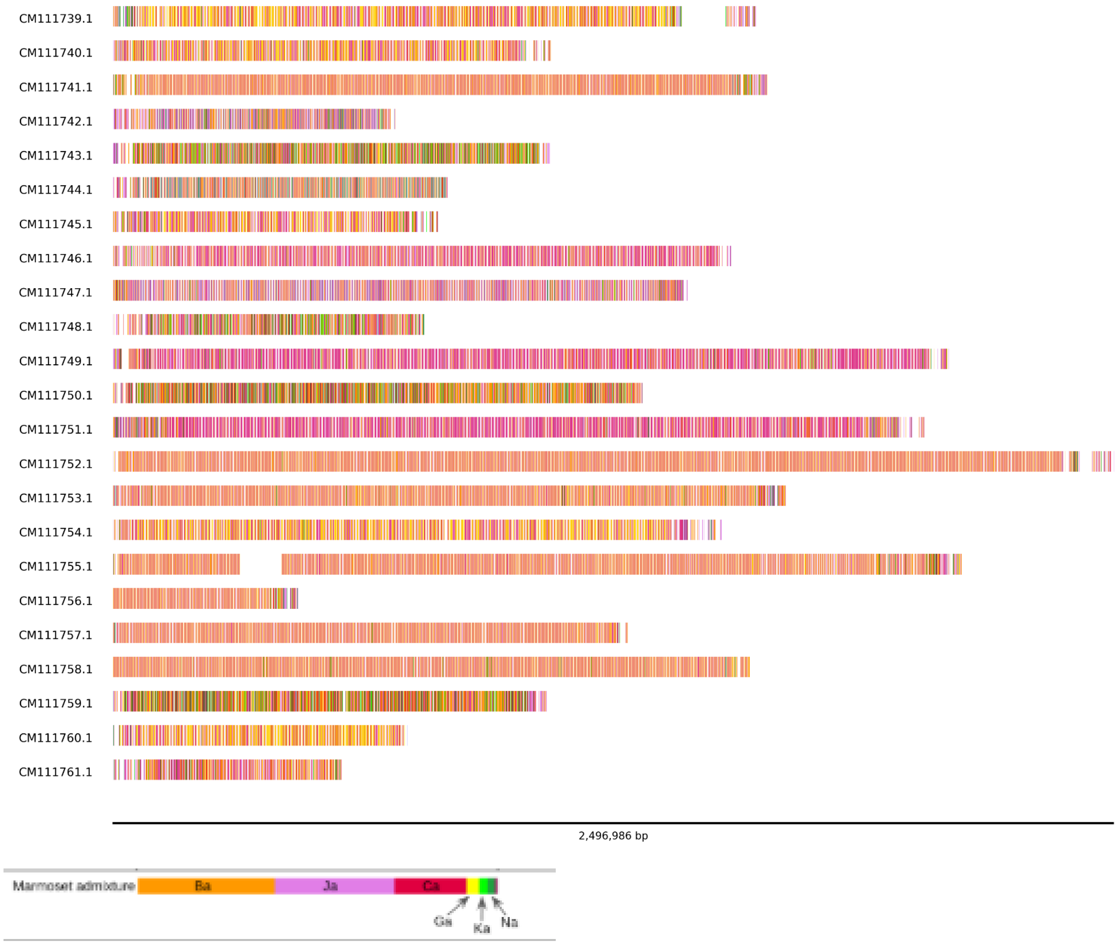


The second method that relied on consensus sequences which have filtered out most of the noise, gave a more convincing result. The tree of all primate monomer species representative of all known SFs was compiled here for the first time and is shown in Fig. **SN14**. It shows that all the NWM-specific monomer types diverge from the tree after Ja (APE-SF11) monomer and before the divergence of Ba and Ca types, which are shared between OWM and apes, but do not appear in NWM-SF annotation in significant numbers. Therefore, we conclude that the LCA of NWM and the OWM/APE branch had APE-SF11 centromeres.

**Fig. SN14. The minimum evolution phylogenetic tree of all primate AS SF consensus monomers.** The top of the tree is formed by monomers that form centromeres in African apes (cyan blocks), which represent APE-SFs 1-3 ("new SFs"). The blue circles (R1 and R2) represent the monomers of orangutan SF5 centromeres. These 2 monomers are the ancestors of all new SF monomers. Yellow circles (Ga) represent the centromeres of gibbons, Ha monomers (brown circles) correspond to the centromeres of the unknown ape ancestor phylogenetically located between OWM and apes. The green empty circles, blocks, and a triangle correspond to the OWM-specific centromeres (empty circles represent monomeric classes of extinct OWM ancestors).

Monomer Ka represents the dead layer, which remains of the centromeres of OWM/APE LCA, and monomers Na and Oa form a dimeric dead layer immediately preceding Ka in OWM and ape pericentromeres. These are followed by Ca and Ba layers. These 3 layers (NaOa, Ca, and Ba) likely identify 3 entirely extinct taxa that separate NWM and OWM LCAs in the primate tree. Ja represents the centromeres of the NWM LCA species, and the NWM AS species are designated with purple blocks and empty circles.   One can see that the marmoset p-monomer is clearly more related to the a-monomer from 11mer HOR. Also, the j-monomer of the HOR is likely to be the ancestor of both S3 and S4 monomers, as it is close to the root of the S3/S4 branch.

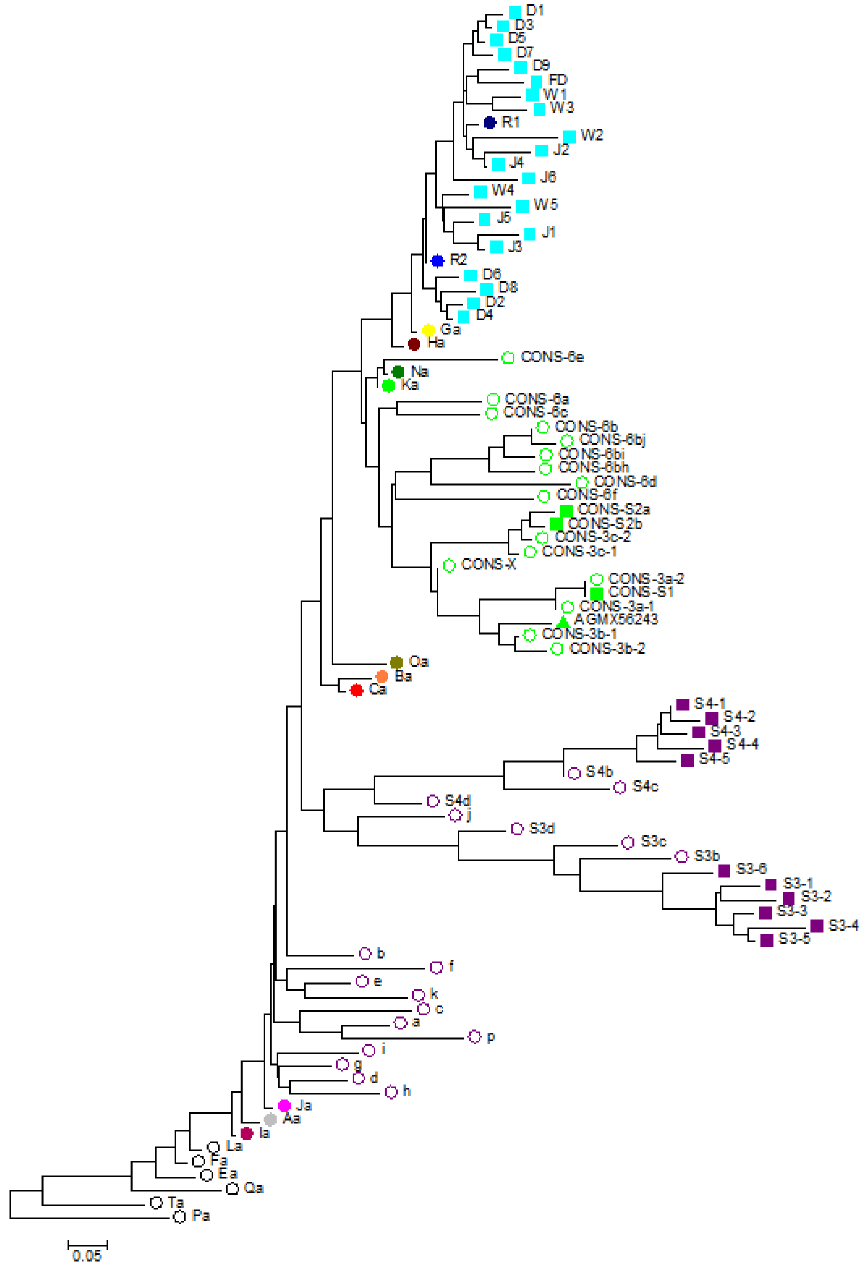


#### Annotation of other NWM primate chromosome-level assemblies

The annotation tool was applied to additional Callitrichidae and Cebidae assemblies (*Saguinus oedipus (GCA_031835075.1)*, *Leontopithecus rosalia (GCA_028533165.1)*, and *Saimiri boliviensis (GCA_048565385.1)*). Comparative analysis revealed that relatively more divergent species (*S. oedipus* and *S. boliviensis*) have satellite types (or their derivatives) already inactive in marmoset (NWM-SF2,

S3bS4b dimers) in their active centromeres (see Fig. **SN15** for the NWM tree). The more closely related *L. rosalia* shares the NWM-SF1 active dimer classification (S3S4) with marmoset but lacks the same marmoset-specific subSFs and exhibits a mixed subSF classification, indicating significant centromere evolution since the species' divergence.

**Fig. SN15. NWM phylogenetic tree**

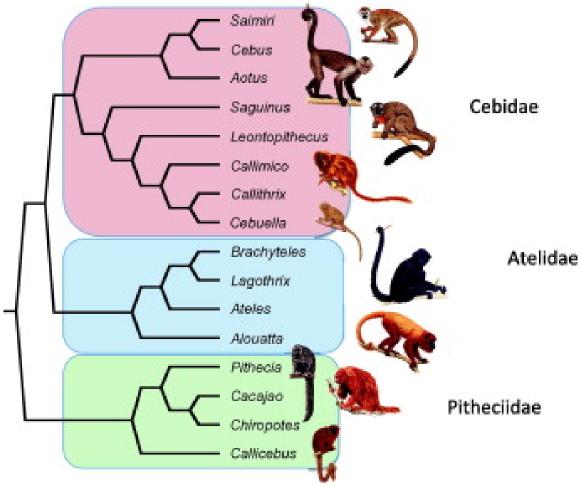


**Table SN6. Comparison of annotation statistics for 4 Cebidae species**

| | Saguinus oedipus | Leontopithecus rosalia | Saimiri boliviensis | Callithrix jacchus (marmoset) |
|---|---|---|---|---|
| dimer | S3bS4b | S3S4 / S3bS4b | S3bS4b | S3S4 |
| sum | 793,405 | 26,820 no full cens? | 256,108 | 192,188 |
| S3-1 | 47 | 68 | 38 | 32,489 |
| S4-1 | 31 | 202 | 13 | 31,904 |
| S3-2 | 17 | 18 | 10 | 5,839 |
| S4-2 | 13 | 229 | 15 | 6,042 |
| S3-3 | 8 | 165 | 21 | 16,929 |
| S4-3 | 14 | 161 | 9 | 15,984 |
| S3-4 | 18 | 55 | 25 | 15,648 |
| S4-4 | 16 | 155 | 18 | 15,677 |
| S3-5 | 42 | 942 | 26 | 14,110 |
| S4-5 | 123 | 7,026 | 17 | 17,860 |
| S3-6 | 85 | 6,115 | 83 | 6,568 |
| S4-6 | 11 | 69 | 16 | 3,206 |
| S3b | 394,260 | 2,987 | 125,980 | 2,054 |
| S4b | 394,193 | 2,615 | 127,004 | 2,584 |
| S3c | 471 | 1,058 | 1,013 | 1,206 |
| S4c | 293 | 1,021 | 164 | 1,414 |
| S3d | 885 | 300 | 120 | 227 |
| S4d | 989 | 281 | 75 | 197 |

#### <u>Annotation of Macaca (OWM) chromosome Y centromere</u>

As we proposed a hypothesis on the relationship of AS in active centromeres of acrocentric chromosomes with the presence/absence of rDNA in their short arms, which included a possible explanation of the delayed cenY phenomenon, we were interested in a complete survey of the delayed/non-delayed status of primate cenYs in all major primate branches (NWM, OWM, and apes). As we recently reported, the analysis of ape cenYs and the NWM cenY is presented here; we only missed the OWM cenY, which had not been annotated previously. Although we developed the AS annotation tool for OWM as part of centromere characterization in the recently published T2T genome of *Macaca fascicularis*[14], that genome lacked the Y chromosome, so it was left unexamined. Since then, however, the assemblies of M. mulatta (MMU, *CM111680.2*), M fascicularis (MFA, *NC_132903.1*), and M. nemestrina (MNE, *NC_092146.1*) chromosomes Y have become available on the Genome Ark site[43], so we took advantage of these data and annotated them for this paper.

It appeared that, in all 3 genomes, cenY active regions were formed by a 20-mer HOR composed of monX AS with 2 monomers, likely falsely recognized as 3a and 3c (Fig. **SN16**). This is in stark contrast to all other active macaque centromeres, which are formed by dimers of S1S2 monomers[14]. As both monX arrays and the arrays of the 3abc trimer represent the inactive, older OWM-specific layers, it follows that macaque cenY is delayed, as are cenYs in humans and chimps[14,31].

**Fig. SN16. Annotation of macaque cenYs**

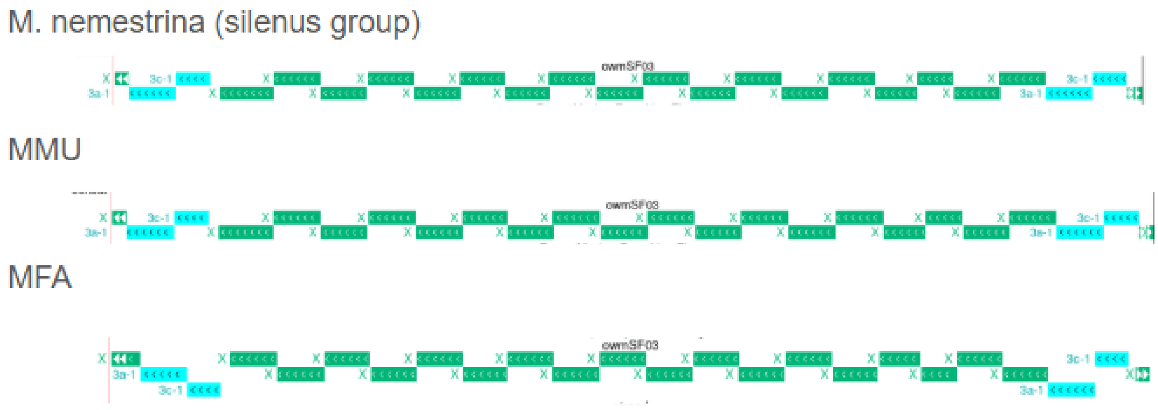
